# Pan-genome analysis identifies intersecting roles for *Pseudomonas* specialized metabolites in potato pathogen inhibition

**DOI:** 10.1101/783258

**Authors:** Alba Pacheco-Moreno, Francesca L. Stefanato, Jonathan J. Ford, Christine Trippel, Simon Uszkoreit, Laura Ferrafiat, Lucia Grenga, Ruth Dickens, Nathan Kelly, Alexander D. H. Kingdon, Liana Ambrosetti, Sergey A. Nepogodiev, Kim C. Findlay, Jitender Cheema, Martin Trick, Govind Chandra, Graham Tomalin, Jacob G. Malone, Andrew W. Truman

## Abstract

Agricultural soil harbors a diverse microbiome that can form beneficial relationships with plants, including the inhibition of plant pathogens. *Pseudomonas* spp. are one of the most abundant bacterial genera in the soil and rhizosphere and play important roles in promoting plant health. However, the genetic determinants of this beneficial activity are only partially understood. Here, we genetically and phenotypically characterize the *Pseudomonas fluorescens* population in a commercial potato field, where we identify strong correlations between specialized metabolite biosynthesis and antagonism of the potato pathogens *Streptomyces scabies* and *Phytophthora infestans*. Genetic and chemical analyses identified hydrogen cyanide and cyclic lipopeptides as key specialized metabolites associated with *S. scabies* inhibition, which was supported by *in planta* biocontrol experiments. We show that a single potato field contains a hugely diverse and dynamic population of *Pseudomonas* bacteria, whose capacity to produce specialized metabolites is shaped both by plant colonization and defined environmental inputs.

## Introduction

Plant pathogenic microorganisms are responsible for major crop losses worldwide and represent a substantial threat to food security. Potato scab is one of the main diseases affecting potato quality (1), and presents a significant economic burden to farmers around the world. The Gram-positive bacterium *Streptomyces scabies*, which is the causal organism of potato scab, is ubiquitous and presents a threat in almost all soils (2, 3). Properly managed irrigation is a reasonably effective control measure for potato scab. However, scab outbreaks still regularly occur in irrigated soil, and with increasing pressures on water use it is clear that alternative approaches to the control of scab are needed. An attractive potential alternative involves the exploitation of soil microorganisms that suppress or kill plant pathogens, known as biocontrol agents (4, 5).

Many soil-dwelling *Pseudomonas* species form beneficial relationships with plants, positively affecting nutrition and health (6–8) and exhibiting potent antagonistic behavior towards pathogenic microorganisms (4, 9, 10). *Pseudomonas* influence the plant environment using a diverse range of secondary metabolites (11–14) and secreted proteins (15, 16). As such, *Pseudomonas* sp. have been identified as key biocontrol organisms in numerous plant-microbe systems (17, 18), and these bacteria have potential applications as agricultural biocontrol agents and biofertilizers (4, 19). Many soil pseudomonads belong to the *P. fluorescens* group, which consists of over 50 sub-species and exhibits huge phenotypic and genetic diversity (6, 10, 20–22), with a core genome of about 1,300 genes and a pan-genome of over 30,000 genes (23). These bacteria use a variety of mechanisms to colonize the plant rhizosphere (24), communicate with plants (7), and suppress a range of plant pathogens (9), including bacteria (25), fungi (26) and insects (27), although a single strain is unlikely to have all of these attributes. Specialized metabolites are critical to many of these ecological functions, and the *Pseudomonas* specialized metabolome is one of the richest and best characterized of any bacterial genus (11–13).

Various studies have associated pseudomonads with potato scab suppression. A significant increase in the abundance of *Pseudomonas* taxa has been observed for irrigated fields, correlating with reduced levels of potato scab (28). Naturally scab-suppressive soils have also been shown to contain a greater proportion of *Pseudomonas* when compared to scab-conducive soils (29, 30), and phenazine production by *P. fluorescens* can contribute to scab control (14, 25, 31). Differences between soil microbial populations that enable effective pathogen suppression are routinely assessed using amplicon sequencing (29, 32). However, the heterogeneity of the *P. fluorescens* group limits the usefulness of these methods for observing changes at the species or even the genus level. To effectively determine the relationship between the soil *Pseudomonas* population and disease suppression, it is important to accurately survey genotypic and phenotypic variability at the level of individual isolates, and to determine how this variation is linked to agriculturally relevant environmental changes (33).

To investigate the genetic bases for *S. scabies* inhibition by *P. fluorescens* and to assess whether the scab-suppressive effects of irrigation derive from increased populations of biocontrol genotypes in the soil or on the plant, we focused on the *Pseudomonas* population from a potato field susceptible to potato scab. We first employed a phenotype-genotype correlation analysis across *P. fluorescens* strains isolated from a single potato field. We hypothesized that an unbiased correlation analysis would identify genetic loci and biosynthetic gene clusters that may be overlooked by screening for bioactive small molecules or by focusing on the biosynthetic repertoire of a limited number of strains. Here, we correlated phylogeny, phenotypes, specialized metabolism and accessory genome loci, then investigated the importance of strong correlations by genetic manipulation of selected wild isolates. In total, 432 *Pseudomonas* strains were phenotyped (with 69 whole genomes sequenced). This approach also enabled us to answer a number of ancillary questions: how diverse is the *P. fluorescens* population from a single field location? Do the phenotypes associated with a *P. fluorescens* strain correlate with its biosynthetic capacity? What does irrigation do to both the population structure of the *P. fluorescens* group and to the wider bacterial community? Using this approach, we identify the *P. fluorescens* genes, gene clusters and natural products that are required for potato pathogen suppression *in vitro*. We use this data to inform the discovery that the cyclic lipopeptide tensin is a key determinant of *in planta* pathogen suppression by a *Pseudomonas* species. We show that irrigation induces profound and repeatable changes in the microbiome, both on a global level and within the *P. fluorescens* species group. Finally, we propose a model for the relationship between irrigation, pathogen suppression, and population-level shifts within the plant-associated *P. fluorescens* population.

## Results

### Irrigation induces a significant change in the soil microbiome

The ability of irrigation to protect root vegetables against *S. scabies* infection is agriculturally important and widespread, but poorly understood. It is likely that the irrigated soil microbiota plays a role in mediating scab suppression, but how this occurs is unclear. We therefore assessed the impact of irrigation on the total bacterial population of a commercial potato field in the United Kingdom. Multiple soil samples were taken from two sites (A1 and B1) within this field, immediately prior to potato planting in January. Following potato planting, one site was irrigated as normal (site A), while the second was protected from irrigation (site B). Tuber-associated soil was sampled from both sites in May (A2 and B2) when tubers were just forming, and the plants were most susceptible to *S. scabies* infection. Total genomic DNA was then extracted from replicate samples of each site after each sampling event, and 16S rRNA amplicon sequencing was used to examine the bacterial population in each of these sites (Figure 1, Figure S1).

**Figure 1.**
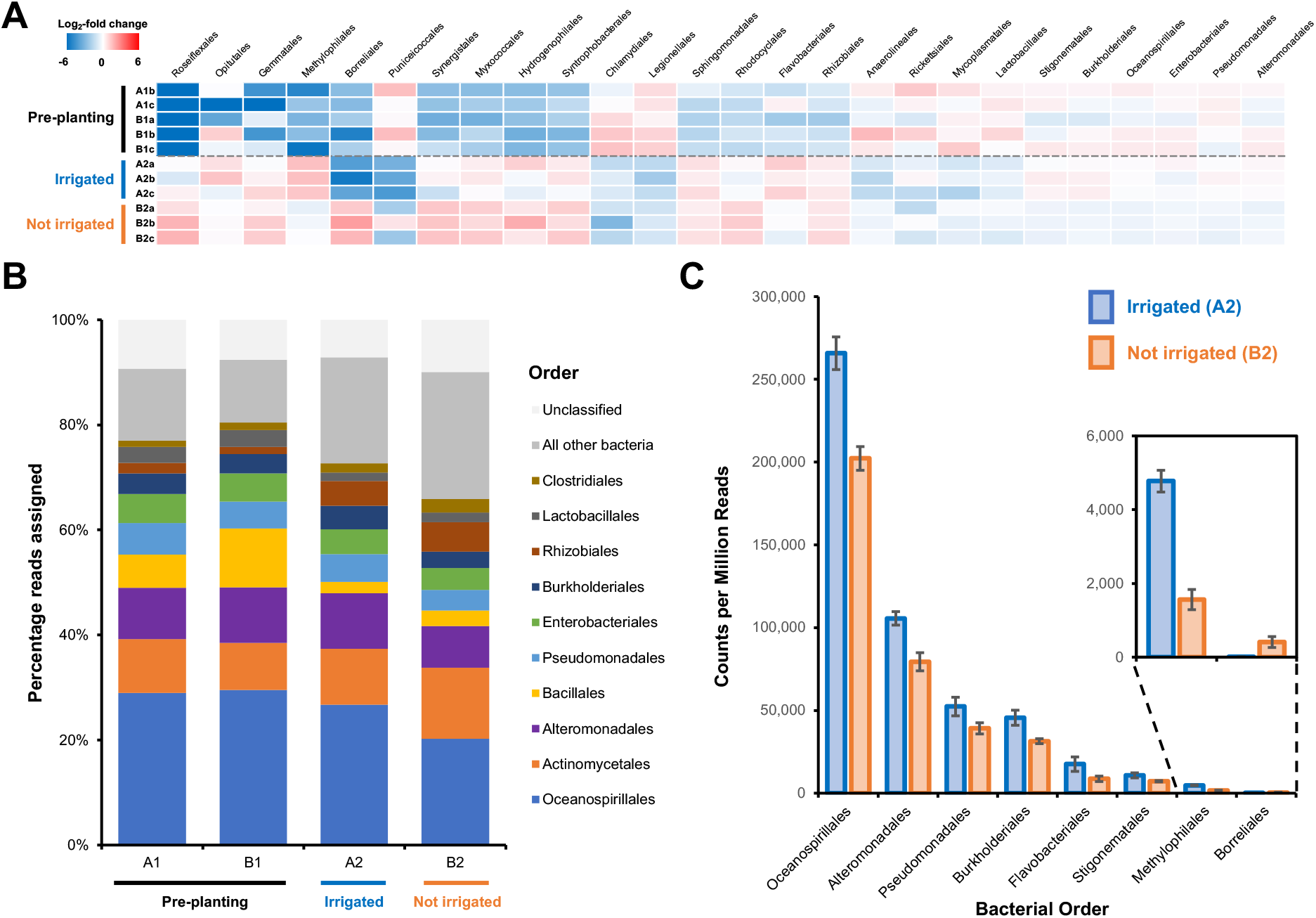
Effect of irrigation on the microbial population of a potato field. (A) The 26 bacterial orders whose populations were determined to significantly differ across one or more sampling sites using voom with a false discovery rate of 0.05. Data is shown as a heatmap of the log2-fold change with respect to the overall average counts per million for a given order. Sample A1a was omitted from the analysis due to possible contamination leading to an atypical bacterial population (Figure S1). (B) Overall average population of each sample site showing the 10 most abundant bacterial orders across all sites. (C) The eight bacterial orders whose populations were determined to significantly differ between irrigated and non-irrigated sites, represented as counts per million reads. Error bars represent the standard deviation of triplicate data.

We used voom (34) with a false discovery rate (FDR) of 0.05 to assess population changes across the four sampling sites. In total, changes were observed for 26 bacterial orders (Figure 1A), with the most significant changes observed between January and May regardless of irrigation. This partially reflects an increase in bacterial orders that have previously been associated with the potato root microbiome (35, 36), including *Rhizobiales*, *Sphingomonadales* and *Flavobacteriales*. Significant population changes (FDR<0.05) were also observed for 8 bacterial orders between the irrigated (A2) and non-irrigated (B2) sites (Figure 1B and 1C), including a larger proportion of *Pseudomonadales* bacteria in the irrigated site. In contrast, despite the potential for microbial heterogeneity across the fertilized field prior to planting, no significant changes were observed between pre-planting sites A1 and B1.

### Phenotypic, phylogenetic and genomic analysis of the *P. fluorescens* field population

Taxonomic identifications using 16S rRNA amplicon analysis showed order-level changes to the field microbiome between sites (Figure S1) but were unable to accurately capture diversity within genera or species groups. Therefore, biologically relevant variation within the populations of genetically diverse species groups such as *P. fluorescens* is potentially overlooked. To investigate the diversity of the fluorescent pseudomonad population, we isolated 240 individual *Pseudomonas* strains from our pre- and post-irrigation field sites (Supporting Dataset 1). These strains were screened for multiple phenotypes including motility, protease production, fluorescence (siderophore production) and on-plate suppression of *S. scabies* using a cross-streak assay (Figures S2 and S3). Each phenotype was scored on an ordinal scale between 0 (no phenotype observed) and 3 (strong phenotype). The cross-streak assay provided a rapid read-out of bacterial antagonism for both contact-dependent and diffusible mechanisms of growth inhibition. On-plate suppression of *S. scabies* was a surprisingly rare trait, with 79% of *Pseudomonas* isolates out-competed by *S. scabies* in this assay. To determine whether this suppressive activity correlated with specific genetic loci, 69 isolates were selected for whole genome sequencing, where almost half (32 strains) exhibited on-plate suppression of *S. scabies* and the remaining strains represented a diverse selection (based on phenotypic variation and 16S rRNA sequencing) of non-suppressive strains. We hypothesized that a comparative analysis of a similar number of genomes from suppressive and non-suppressive strains would identify those biosynthetic gene clusters (BGCs) that play important roles in suppressive activity.

The phylogeny of the 69 sequenced strains was analyzed alongside various model pseudomonads, including representatives of the eight phylogenomic *P. fluorescens* groups defined by Garrido-Sanz *et al.* (23). Our sequenced strain collection spans much of this characterized global phylogenetic diversity and contains representatives of at least five of the eight *P. fluorescens* phylogenomic groups (23), as well as strains belonging to the *P. putida* and *P. syringae* groups (Figure S4). This genetic heterogeneity was also reflected in the diverse specialized metabolome of these strains, as predicted by a detailed analysis of the BGCs encoded in their genomes. Each genome was subjected to antiSMASH 5.0 analysis (37), which was further refined by extensive manual annotation to improve the accuracy of predicted pathway products. This second annotation step was particularly important for BGCs that are atypically distributed across two distinct genomic loci (e.g. viscosin and pyoverdine). Our analysis was further expanded to include BGCs not identified by antiSMASH 5.0, including BGCs for hydrogen cyanide (HCN) (38), microcin B17-like pathways (39) and the auxin indole-3-acetic acid (IAA) (40, 41). This was achieved by searching the genomes with a curated set of known *Pseudomonas* BGCs using MultiGeneBlast (42) (see Supplementary Information for further details). This manual annotation provided a level of resolution superior to that provided by automated cluster-searching algorithms alone and provided confidence that the majority of natural product biosynthetic potential had been identified. Within a given pathway type (e.g. non-ribosomal peptide synthetases, NRPSs), likely pathway products were assigned where possible (e.g. cyclic lipopeptides, CLPs) or assigned a code when a conserved uncharacterized BGC was identified (e.g. NRPS 1). All BGCs were mapped to strain phylogeny (Figure 2).

**Figure 2.**
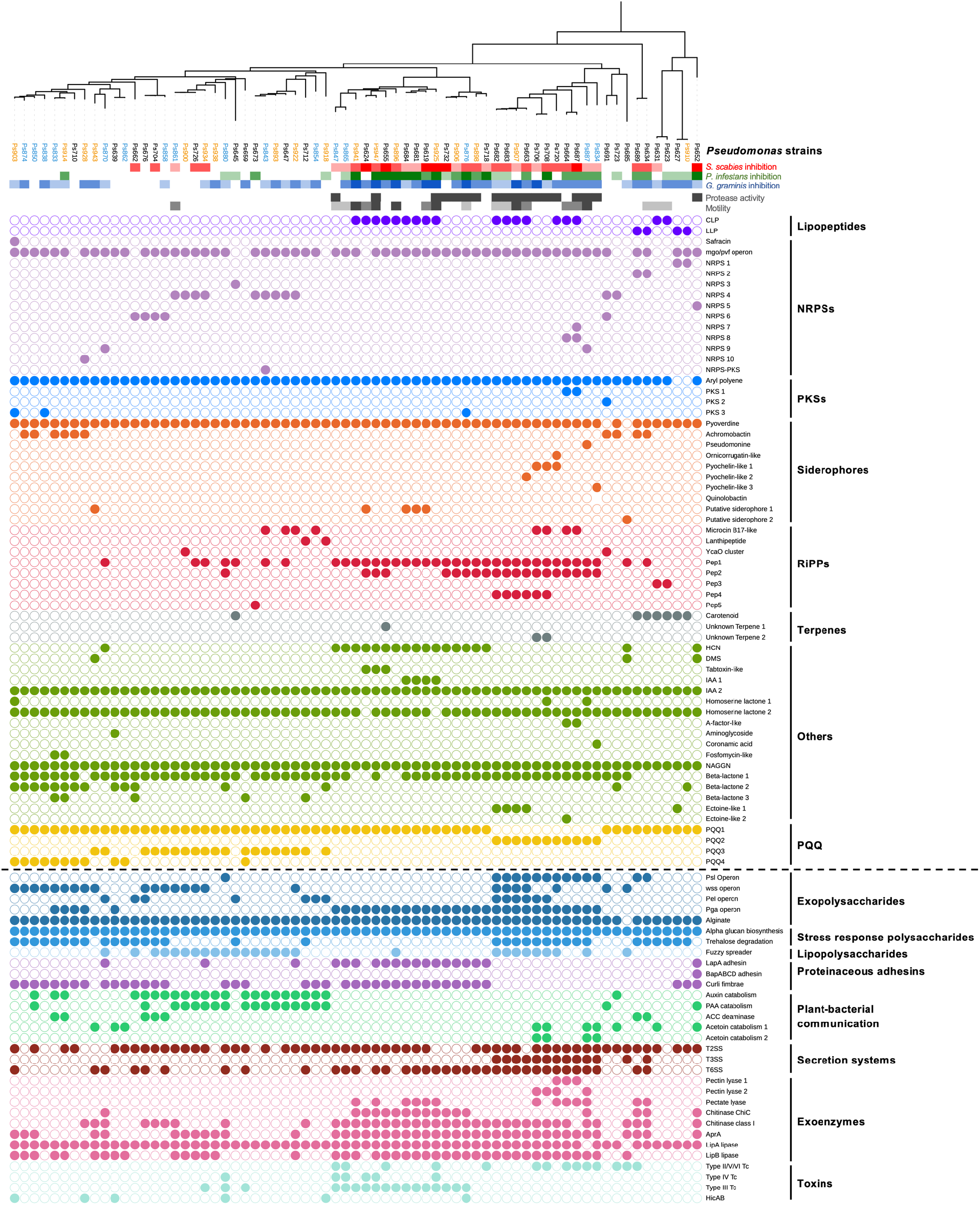
Comparison of phylogeny, *S. scabies* suppression (red colour scales), *P. infestans* suppression (green colour scale), *G. graminis* pv. *tritici* (take-all) suppression (blue colour scale), phenotypes (grey colour scales), natural product biosynthetic gene clusters (filled circles = presence of a gene or gene cluster) and the accessory genome (separated from BGCs by a dotted line). In the phylogenetic tree of *Pseudomonas* strains, blue strains were collected from irrigated plots while orange strains were collected from unirrigated plots. All other strains were collected from the pre-irrigation plots. Figure S4 shows the relationship between these strains and the *Pseudomonas* phylogenomic groups defined by Garrido-Sanz *et al.* (23).

Multiple BGCs were commonly found across the sequenced strains (Figure 2, Supporting Dataset 1), including BGCs predicted to make CLPs (43), arylpolyenes (44) and HCN (38). In addition to pyoverdine BGCs (45) in almost all strains, numerous other siderophore BGCs were identified, including pathways predicted to make achromobactin (46), ornicorrugatin (47), pyochelin-like molecules (48) (Figure S5) and a pseudomonine-like molecule (49). A variety of polyketide synthase (PKS), terpene and NRPS BGCs with no characterised homologues were also identified (Figure 2). Furthermore, BGCs were identified that were predicted to make compounds related to microcin B17 (39), fosfomycin-like antibiotics (50), lanthipeptides (51), safracin (52), a carbapenem (53) and an aminoglycoside (54) (Figure S6). Each of these natural product classes is predicted to have potent biological activity and some are rarely found in pseudomonads.

In addition to these potentially antibacterial and cytotoxic compounds, all genomes contain BGCs predicted to produce the plant auxin IAA, while 23 genomes contained genes for IAA catabolism (55). All 69 strains had at least one BGC for the production of the electron-transport cofactor pyrroloquinoline quinone (56) (PQQ, Figure S7), reported to function as a plant growth promoter (57). Surprisingly, BGCs for numerous well characterized *Pseudomonas* specialized metabolites were not found, including phenazine, pyrrolnitrin or 2,4-diacylphloroglucinol BGCs (11). In total, 787 gene clusters were identified that could be sub-divided into 61 gene cluster families (Figure 2).

The *P. fluorescens* species group possesses a highly diverse array of non-essential accessory genes and gene clusters. These are often critical to the lifestyle of a given strain, and can include motility determinants, proteases, secretion systems, polysaccharides, toxins and metabolite catabolism pathways (17). These accessory genome loci were identified using MultiGeneBlast (details in Supplementary Information), which revealed a high degree of genomic diversity across strains. Specialized metabolism BGCs and accessory genome loci were mapped to strain phylogeny (Figure 2), which indicated that for some loci (e.g. the *psl* operon, auxin catabolism, HCN biosynthesis) there is a close, but not absolute, relationship between phylogeny and the presence of a gene cluster.

### Correlation analysis identifies potential genetic determinants of *S. scabies* inhibition

We hypothesised that genes associated with suppression of *S. scabies* could be identified by a correlation analysis between *S. scabies* cross-streak inhibition and the presence of BGC families or accessory genes. We therefore calculated Pearson correlation coefficients for each BGC with *S. scabies* inhibition (Figure S8). The top 10 positively correlating genotypes and phenotypes (Figure 3) comprised four BGC families (Pep1, CLP, Pep2 and HCN) (Figure S9), four accessory genome loci (chitinase ChiC, protease AprA, chitinase class 1 and the Pga operon) and two phenotypes (motility and secreted protease production). The production of HCN and/or CLPs by *Pseudomonas* strains have been previously associated with the suppression of various plant pathogens including fungi (26, 58) and oomycetes (59, 60), and can also contribute to insect killing (27, 61), but have not been linked to the suppression of bacteria. A variety of genotypes associated with plant-microbe interactions were moderately negatively correlated with suppression (ρ< −0.3), including BGCs for PQQ biosynthesis and catabolism of the plant auxins IAA and phenylacetic acid (PAA) (Figure 3A).

**Figure 3.**
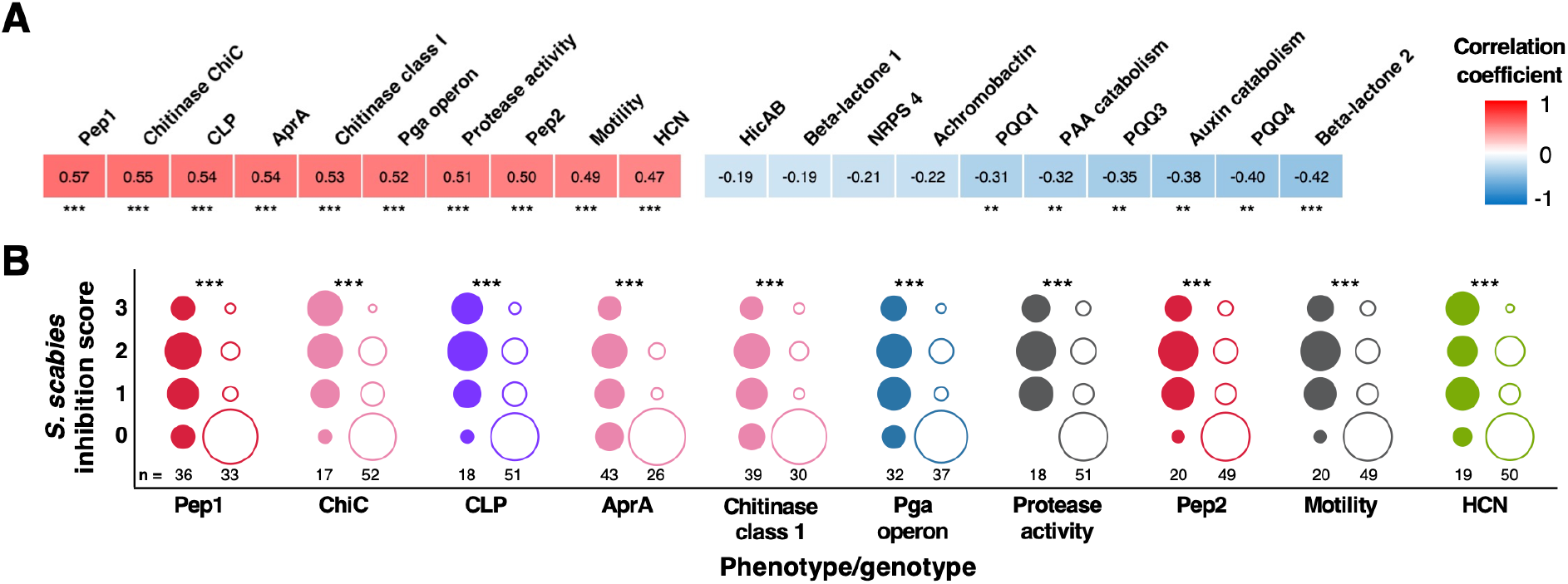
Correlation of BGCs and accessory genome loci with *S. scabies* inhibition. (A) Heatmap showing the 10 genotypes and phenotypes that correlated most strongly (positively and negatively) with on-plate suppression of *S. scabies*. Stars represent the statistical significance of a correlation using a two-tailed Mann-Whitney test (p<0.05 = *, p<0.01 = **, p<0.001 = ***). (B) Distributions of *S. scabies* suppressive activity for top 10 positive correlations. Circles are stacked from no (0) to high (3) inhibition, where filled and empty circles represent strains with and without a given genotype/phenotype, respectively. The number of strains (total = 69) in each class is listed, and the area of a circle specifies the proportion of strains with given suppressive activity.

Interestingly, while certain BGC loci (e.g. CLP) positively correlated with both suppression and motility, this relationship was not seen for every locus (e.g. HCN correlates with suppression but is less strongly correlated with motility). Correlation does not equate to causation, especially considering the significant evolutionary association seen for some BGCs (Figure 3C). The importance of correlating BGCs to *S. scabies* suppression was therefore investigated experimentally using a genetically tractable subset of suppressive isolates.

### Production and role of CLPs in suppression of *S. scabies*

The strong positive correlation between putative CLP gene clusters and *S. scabies* suppression prompted us to investigate whether CLPs play a role in suppressive activity. *Pseudomonas* CLPs have previously been associated with a wide array of functions, including fungal growth inhibition, plant colonisation and promotion of swarming motility (43, 62), although there are no reports of *Pseudomonas* CLPs functioning as inhibitors of streptomycete growth. However, prior work has shown that surfactin, a CLP from *Bacillus subtilis,* inhibits *Streptomyces coelicolor* aerial hyphae development (63), while iturin A, a CLP from *Bacillus* sp. sunhua, inhibits *S. scabies* development (64).

To determine the identity of each CLP we combined bioinformatic predictions of the NRPS products (37) with experimental identification using liquid chromatography – tandem mass spectrometry (LC-MS/MS). In every strain that contained a CLP BGC, a molecule with an expected mass and MS/MS fragmentation pattern was identified (Figure 4, Figures S10-S16). This showed that *P. fluorescens* strains from a single field have the collective capacity to make viscosin (*m/z* 1126.69, identical retention time to a viscosin standard) (65), a viscosin isomer (*m/z* 1126.69, different retention time to viscosin standard) (Figures S10 and S11), as well as compounds with BGCs, exact masses and MS/MS fragmentation consistent with tensin (*m/z* 1409.85, Figure S12) (66), anikasin (*m/z* 1354.81, Figure S13) (67), and putisolvin II (*m/z* 1394.85, Figure S14) (68). In addition, an array of related metabolites were observed that differed by 14 or 28 Da, which is characteristic of different lipid chain lengths. This analysis also proved that the linear lipopeptides syringafactin A (*m/z* 1082.74) and cichofactin (*m/z* 555.38, [M+2H]^2+^) were made by strains harbouring BGCs predicted to make these phytotoxins (69, 70) (Figures S15 and S16). The metabolic capacity of all strains was mapped using mass spectral networking (71, 72), which showed that CLPs were strongly associated with strains that inhibit *S. scabies* (Figure 4).

**Figure 4.**
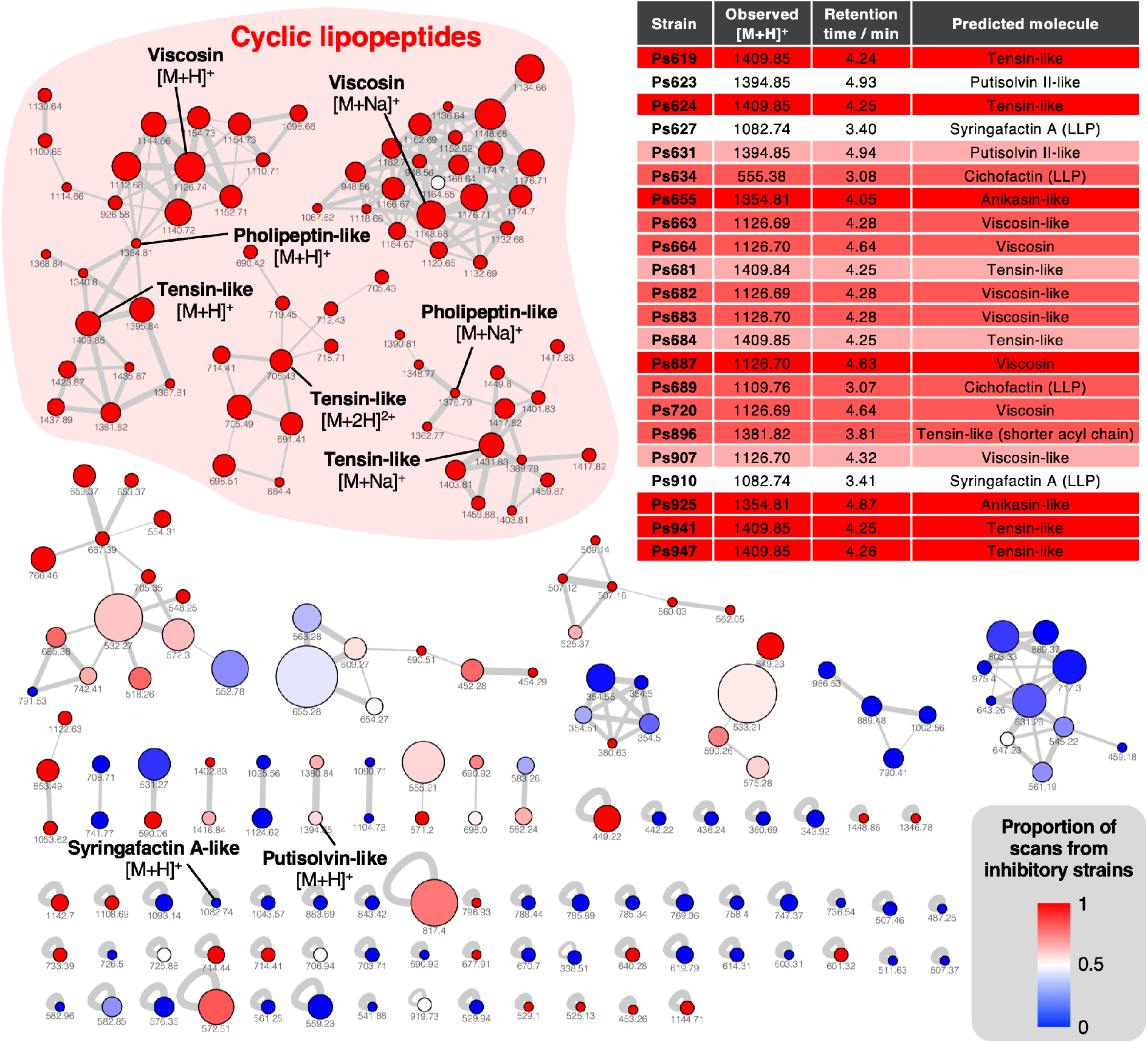
Mass spectral networking analysis of LC-MS/MS data from the *Pseudomonas* strains used in this study. Node area is proportional to the number of distinct strains where MS/MS data was acquired for a given metabolite. Node colour reflects the proportion of MS/MS scans for a given node that come from strains with a *S. scabies* inhibition score ≥1. Nodes are labelled with the corresponding parent masses and nodes that relate to lipopeptides are labelled (multiple networks arise from differential fragmentation of [M+H]^+^, [M+2H]^2+^ and [M+Na]^+^ ions). Line thickness is proportional to the cosine similarity score calculated by Global Natural Product Social Molecular Networking (GNPS) (72). The table shows production of lipopeptides by strains containing lipopeptide BGCs. Colour-coding reflects level of *S. scabies* inhibition by each strain with same scale as Figure 2 (LLP = linear lipopeptide; all others are CLPs).

To assess the potential role of CLPs in mediating the interaction between *P. fluorescens* and *S. scabies*, an NRPS gene predicted to be involved in the biosynthesis of a viscosin-like molecule in Ps682 was deleted by allelic replacement (Figure 5A). The resulting Ps682 *Δvisc* strain was unable to make the viscosin-like molecule (*m/z* 1126.69, Figure 5B), or to undergo swarming motility (Figure S17). This is in agreement with earlier work on the role of viscosin in the motility of *P. fluorescens* SBW25 (62) and the observation that possession of a CLP BGC was the genotype that most strongly correlated with motility (ρ = 0.65, Figure S8). A cross-streak assay with *S. scabies* revealed an active role for this CLP in on-plate *S. scabies* inhibition (Figure 5D). Wild type (WT) Ps682 appeared to specifically colonise the *S. scabies* streak, whereas Ps682 *Δvisc* was unable to restrict *S. scabies* growth. Alternatively, it was possible that this instead could reflect diffusible inhibition of *Streptomyces* development by WT Ps682, leading to a “bald” *S. scabies* phenotype (73). To distinguish between these possible inhibition modes, a constitutively-expressed *lux* operon was integrated into the chromosomal *att::*Tn*7* site (74) of Ps682 to visualise this interaction by bioluminescence. This clearly showed viscosin-dependent *Pseudomonas* colonisation of the *Streptomyces* streaks (Figure 5C).

**Figure 5.**
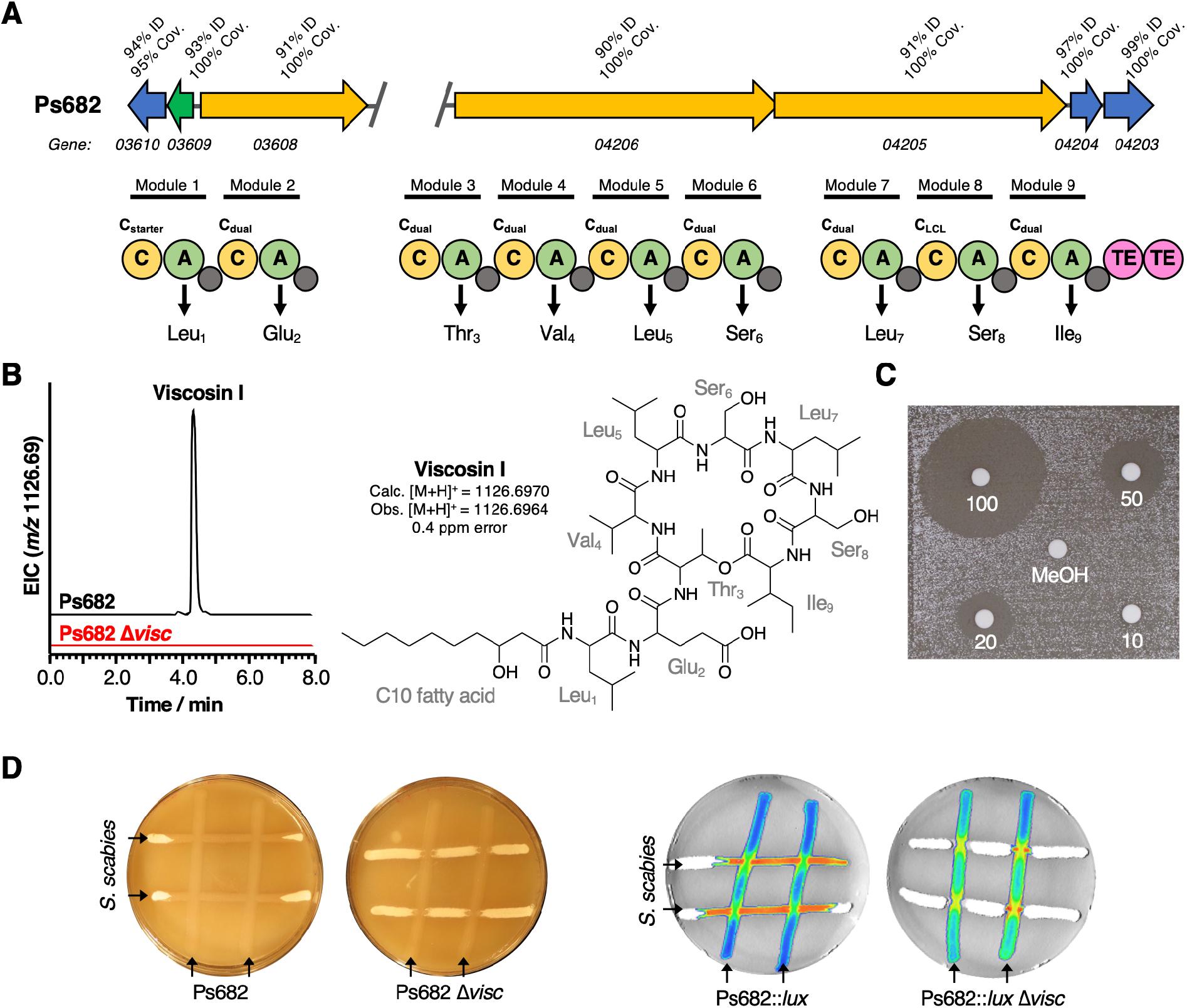
The role of the Ps682 CLP BGC in *S. scabies* suppression. (A) BGC displaying identity/coverage scores in comparison to the viscosin BGC in *P. fluorescens* SBW25. Genes encoding regulatory proteins are green, transporter genes are blue, and NRPS genes are yellow. The NRPS organization is shown, where C = condensation domain, A = adenylation domain, TE = thioesterase domain, and grey circles are peptidyl carrier protein domains. Amino acids incorporated by each module are displayed, along with predicted condensation domain specificity. (B) LC-MS analysis of viscosin I production in the WT strain and a mutant (Ps682 Δ*visc*) with an in-frame deletion of NRPS gene 04206. NMR and MS/MS data for viscosin I is shown in Figures S19-S29. (C) Disk diffusion assay of viscosin I against *S. scabies*. Concentrations are indicated (µg/mL), alongside a methanol control. (D) On-plate *S. scabies* suppression activity of Ps682 alongside Ps682 *Δvisc* shown as cross-streaks using strains with and without the *lux* operon. Bioluminescence was detected using a NightOWL camera (Berthold Technologies).

To quantitatively assess the antagonistic effect of the Ps682 CLP, it was purified and structurally characterized using MS/MS (Figure S18) and nuclear magnetic resonance (NMR) spectroscopy (^1^H, ^13^C, COSY HSQC, TOCSY, HMBC, Figures S19-S29, Table S4). NMR analysis revealed that the molecule has an identical amino acid composition to viscosin (3-hydroxydecanoic acid-Leu1-Glu2-Thr3-Val4-Leu5-Ser6-Leu7-Ser8-Ile9, Figure 5B), which was fully supported by detailed high-resolution MS (calculated viscosin [M+H]^+^ = 1126.6970, observed [M+H]^+^ = 1126.6964) and MS/MS fragmentation data (Figure S18). The LC retention time of this CLP is different to viscosin, but is almost identical to WLIP (Figure S18), which is a viscosin isomer that has a D-Leu5 residue instead of L-Leu5 (75). However, comparison of NMR data in DMF-d7 revealed some minor shift differences between published WLIP spectra (75) and the Ps682 CLP, such as the γ-CH_2_ group of Glu2 (WLIP = δ_H_ 2.54 ppm, δ_C_ 30.3 ppm; Ps682 CLP = δ_H_ 2.24 ppm, δ_C_ 34.8 ppm). Therefore, we could not conclusively confirm the absolute configuration of the Ps682 CLP and thus named it viscosin I (for viscosin Isomer). A disk diffusion assay of purified viscosin I with *S. scabies* (Figure 5D) demonstrated that it directly inhibited *S. scabies* growth with a minimum inhibitory concentration of approximately 20 µg/mL. Long-term growth of *S. scabies* in the presence of viscosin I (Figure S30) indicated that the inhibition of *S. scabies* is temporary and growth partially resumes after several days. These data show that in addition to its role as a surfactant, viscosin I functions by inhibiting the growth rate of *S. scabies*, consistent with the on-plate data for Ps682 *Δvisc*.

### HCN and CLP production both contribute to on-plate *S. scabies* inhibition

Pan-genome analysis showed that HCN production was predicted for a significant number of suppressive strains (ρ = 0.47, Figure S8), where 17 of the 19 strains containing HCN gene clusters were inhibitory towards *S. scabies* (Figure 2). The HCN pathway is encoded by the *hcnABC* gene cluster (Figure S9) and has previously been associated with insect and fungal pathogen inhibition in other *Pseudomonas* strains (27) (59, 76). HCN is toxic to a wide variety of organisms, but not to *Pseudomonas* owing to their branched aerobic respiratory chain that has at least five terminal oxidases, including a cyanide-insensitive oxidase (77, 78). We confirmed that nearly every strain with the *hcnABC* gene cluster produced HCN (18 out of 19) using the Feigl-Anger colorimetric detection reagent (79) (Supporting Dataset 1) and used this assay to identify HCN producers across the original collection of 240 *Pseudomonas* strains. This wider analysis showed that HCN production strongly correlated with *S. scabies* inhibition (ρ = 0.52, Figure S31), in accordance with our analysis of the sequenced strains.

To examine the role of HCN in *S. scabies* suppression and whether it exhibited a synergistic effect with CLP production, Ps619 was investigated as this strain produces both HCN and a tensin-like CLP (Figure 2 and Figure 6A). A tensin BGC has not previously been reported, but the predicted amino acid specificity, mass (Figure 6A) and MS/MS fragmentation (Figure S12) indicated that seven isolates produce tensin-like CLPs (Figure 4). The *hcn* and *ten* gene clusters were inactivated by in-frame deletions to generate single and double mutants of Ps619, and the resulting Δ*hcn*, Δ*ten* and Δ*hcn*Δ*ten* mutants were subjected to cross-streak assays (Figure 6B). A comparison of wild type, single and double mutants showed that HCN inhibits *S. scabies* growth and development across the entire plate, while tensin is important for *Pseudomonas* motility and helps the *Pseudomonas* to grow competitively at the cross-streak interface. Furthermore, this suppressive effect is additive: the Ps619 Δ*hcn* and Δ*ten* single mutants both retained some inhibitory activity towards *S. scabies*, whereas the Ps619 Δ*hcn*Δ*ten* double mutant could not inhibit *S. scabies*.

**Figure 6.**
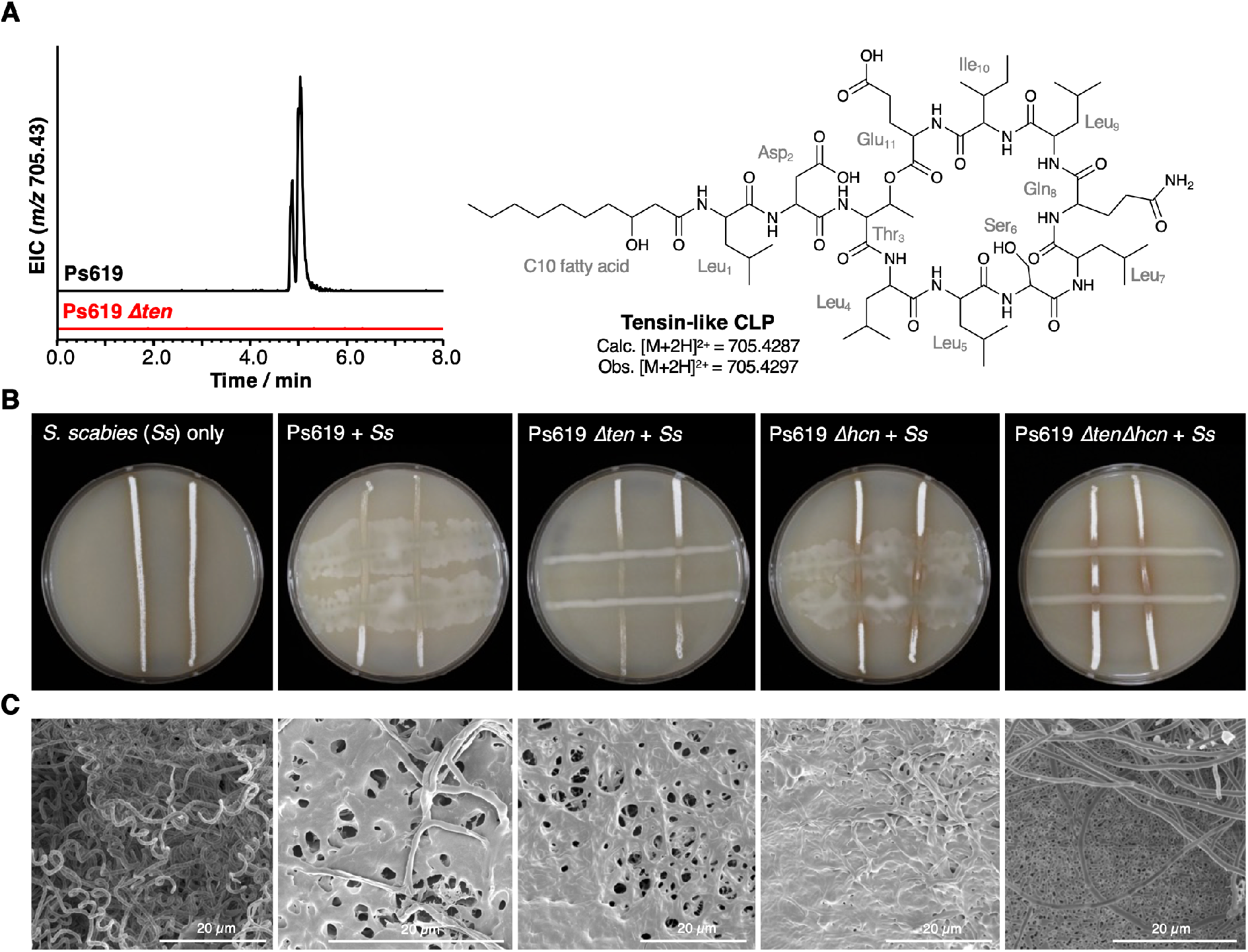
The role of the Ps619 CLP and HCN gene clusters in *S. scabies* suppression. (A) Predicted structure of the tensin-like molecule and LC-MS analysis of CLP production in WT Ps619 and a mutant (Ps619 *Δten*) with an in-frame deletion of NRPS gene 02963 (see Figure S12 for BGC information). (B) Cross streak assays of Ps619 and associated mutants with *S. scabies*. See Figure S32 for assays with drier plates. (C) Cryo-SEM images of the interfacial region between the Ps619 strains and *S. scabies*. The order of images is identical to the cross-streaks in panel B.

In drier growth conditions expected to favour streptomycete growth and limit motility, the role of tensin-mediated motility was abrogated, yet tensin and HCN still possessed an additive inhibitory effect at the microbial interface (Figure S32). Notably, Ps619 Δ*hcn* was able to induce a developmental defect in *S. scabies* at the microbial interface that was not present in Ps619 Δ*ten* or Ps619 Δ*hcn*Δ*ten*, showing that the tensin-like CLP induces a developmental defect in *S. scabies* that is independent of *Pseudomonas* motility, comparable to the inhibitory effect of isolated viscosin I. This analysis also clearly showed that at areas distant from the bacterial interaction, *S. scabies* grew more vigorously when cultured with Δ*hcn* strains, consistent with the volatility of HCN enabling a long-range inhibitory effect. A similar volatile effect was seen when Ps619 strains were separated from *S. scabies* by a barrier, where only those strains producing HCN inhibited growth and development (Figure S32).

To further probe how tensin and HCN affected the interaction between Ps619 strains and *S. scabies*, the interfacial regions of cross-streaks were imaged using Cryo-scanning electron microscopy (cryo-SEM). WT Ps619 was able to colonise the *S. scabies* streak meaning that the interfacial region imaged was further from the cross-streak intersection than all other co-cultures (Figure 6C). Here, Ps619 inhibited *S. scabies* development, which appears as a mixture of deformed aerial hyphae and vegetative growth reminiscent of a “bald” phenotype (80). Cryo-SEM indicated that both the Ps619 Δ*hcn* and Δ*ten* mutants induced a similar partially bald phenotype in *S. scabies*, but the Δ*hcn*Δ*ten* double mutant was unable to trigger the same developmental defect, as *S. scabies* could develop aerial mycelia close to the microbial interface (Figure 6C and Figure S33). This appears as a clear boundary between Ps619 Δ*hcn*Δ*ten* (single cells in background, bottom right panel of Fig 6C) and *S. scabies* (hyphae in the foreground). The volatile HCN can inhibit growth and development at a distance, whereas CLP inhibition of development only occurs close to the microbial interface. Both inhibitory mechanisms enable Ps619 to obtain a competitive advantage at the microbe-microbe interface (Figure 6C and Figure S33), while the CLP also functions as a surfactant enabling Ps619 motility, promoting *Pseudomonas* invasion of the *Streptomyces* cross-streak.

### Tensin is a key determinant of *in planta* inhibition of potato scab

To examine the *in planta* biocontrol properties of Ps619 and Ps682, and to determine the contribution of HCN and CLPs to activity, potato scab suppression assays were carried out in glasshouse trials. Maris Piper potatoes were infected with *S. scabies* 87-22 and scored for disease severity after 16 weeks using the method of Andrade *et al*. (81). A subset of plants was also treated with *Pseudomonas* spp. and associated BGC mutants. Ps619 conferred significant protection against potato scab, where disease severity was reduced to levels similar to uninfected control plants (Figure 7). This suppressive ability was lost for Ps619 Δ*ten* and Ps619 Δ*hcn*Δ*ten*, resulting in disease severity similar to scab-infected tubers. In contrast, Ps619 Δ*hcn* was just as effective as WT Ps619 at suppressing potato scab, which differed from the on-plate results for HCN. The significance of these results was supported by an independent *in planta* biocontrol experiment, where equivalent results were obtained for each strain (Figure S34). This result indicates that tensin plays an important role in the biocontrol of potato scab. In contrast with its on-plate suppressive activity, potato scab assays showed no significant antagonistic activity for Ps682 against *S. scabies* infection. Unfortunately, this meant that the role of viscosin I could not be determined *in planta*.

**Figure 7.**
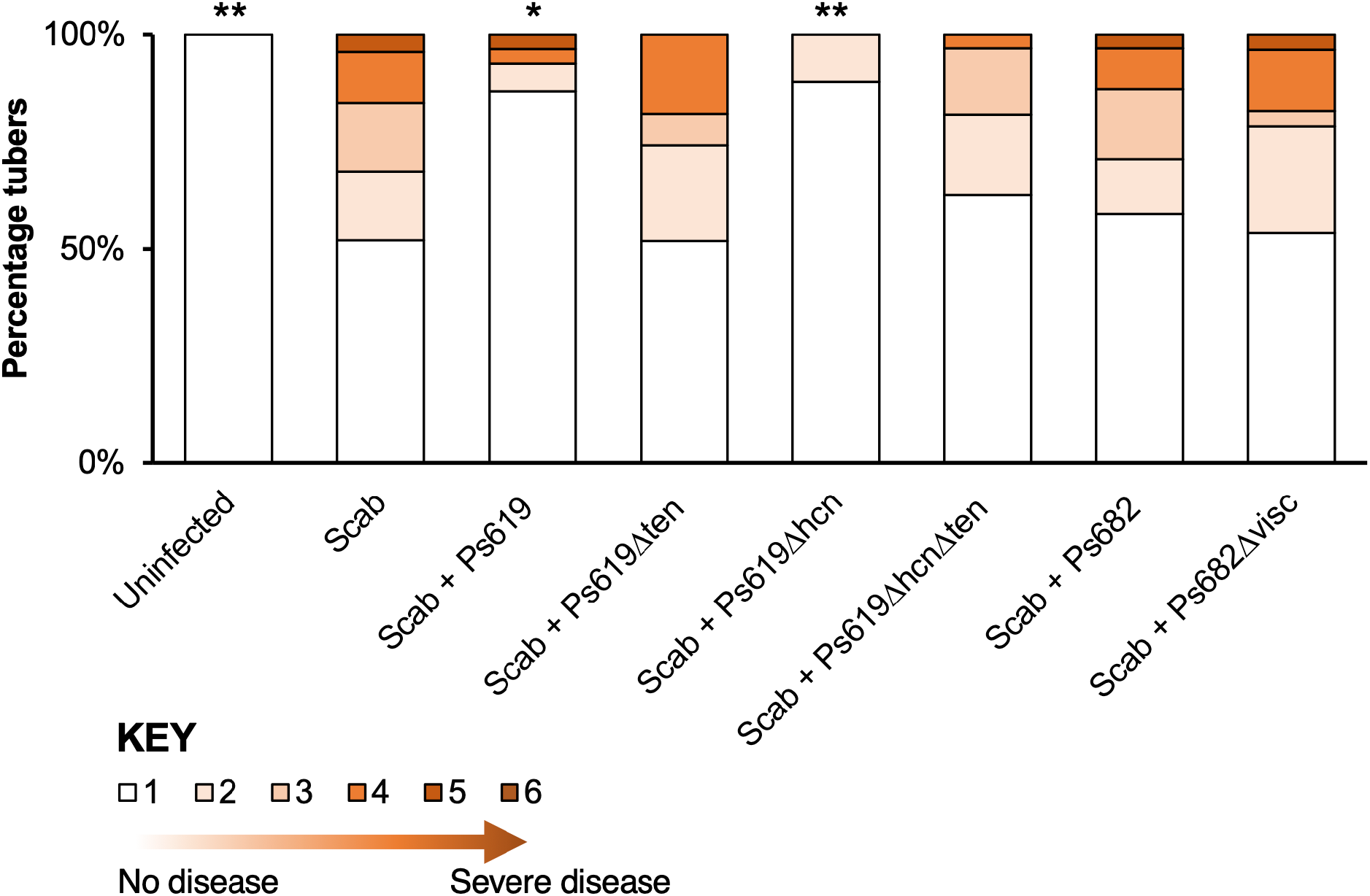
Potato scab biocontrol assay. The bar chart shows the percentage of diseased tubers following infection with *S. scabies* (“Scab”) along with treatment by Ps619, Ps682 and associated mutants. Tubers were scored using a disease severity index from 1 to 6 according to the method of Andrade *et al*. (81). Statistical analyses were calculated by taking into account the average disease index of each plant (n=4). *p* values were calculated using Dunnett’s multiple comparison test and asterisks indicate *p* < 0.05 (*), 0.01 (**) as compared to Scab treatment only. Results of a repeat biocontrol experiment and further statistical statistics are shown in Figure S34.

### A subset of *P. fluorescens* strains are generalist pathogen suppressors

To determine whether the strains and metabolites we identified have suppressive activity towards a range of plant pathogens, we investigated the ability of the potato field strain collection to suppress the growth of *Phytophthora infestans*, the oomycete that causes potato blight (82), and *Gaeumannomyces graminis* var. *tritici*, the fungus that causes take-all disease of cereal crops (17). These assays revealed strong congruence between the genotypes that correlated with suppression of each pathogen (Figure S35). HCN and CLPs have both previously been identified as inhibitors of oomycete and fungal growth (26, 59). To assess whether these natural products are critical for inhibition of *P. infestans* and *G. graminis* by Ps619 and Ps682, the HCN/CLP mutants were tested for inhibitory activity (Figure S35). Surprisingly, neither HCN or tensin were required for Ps619 inhibition of either *G. graminis* or *P. infestans* (Figure S35), indicating the production of at least one other secreted inhibitory factor. In contrast, inactivation of the viscosin I pathway in Ps682 abolished activity towards both pathogens (Figure 8C). These data indicate that a subset of pseudomonads can function as generalist pathogen suppressors, possessing multiple growth inhibition mechanisms (e.g. Ps619) and/or by producing molecules with broad bioactivity (e.g. Ps682).

**Figure 8.**
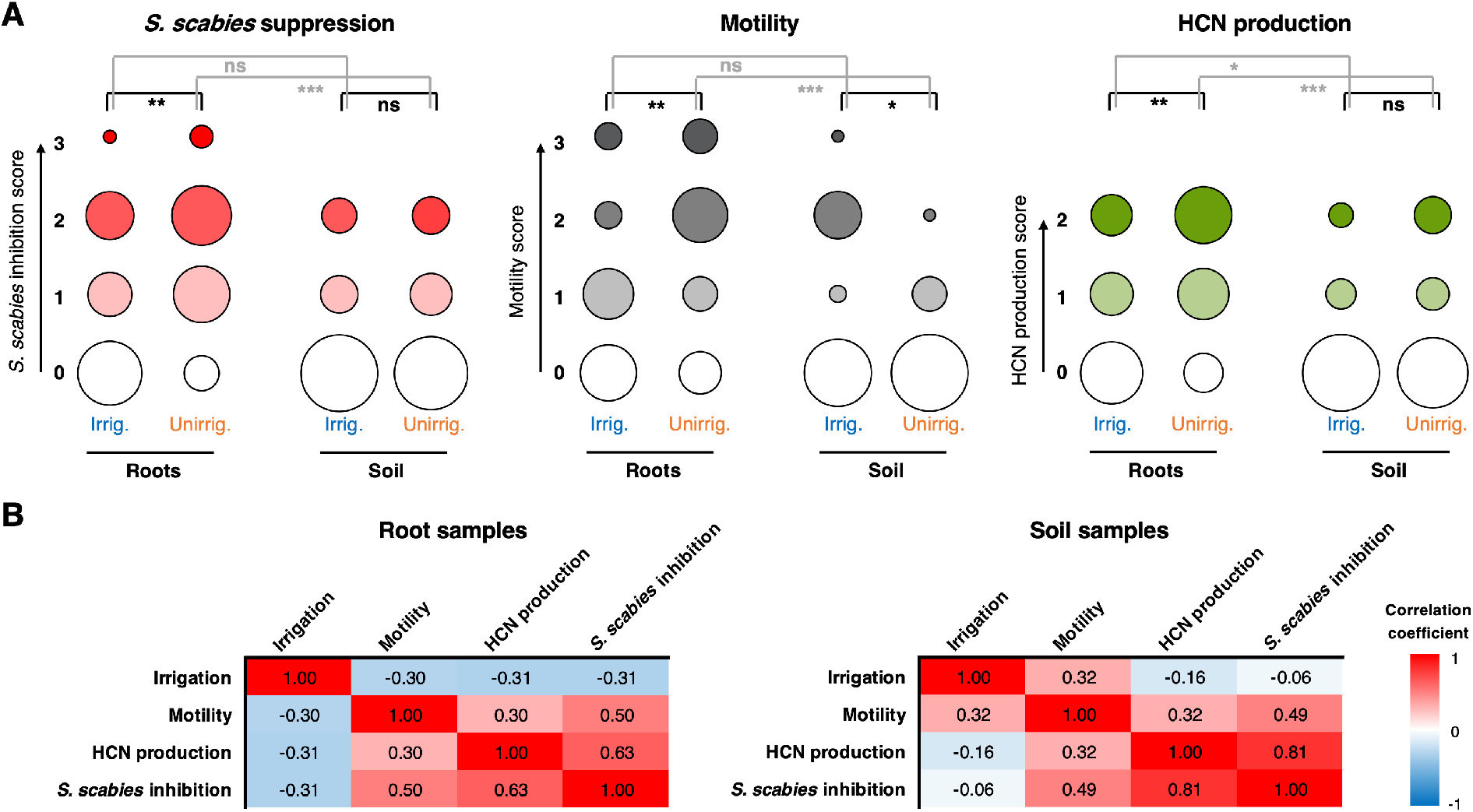
The effect of irrigation and environment (soil versus root) on the *P. fluorescens* population. (A) Plots showing the proportion of strains exhibiting a particular phenotype from each environment (n = 48 for each condition). HCN production was scored on a scale of 0-2 based on a qualitative assessment of the colour change in the Feigl-Anger assay. The size of each circle is proportional to the number of strains with a given phenotypic score. Statistical comparisons were carried out two-tailed Mann-Whitney tests where ns (not significant) = p≥0.05, * = p<0.05, ** = p<0.01, *** = p<0.001. (B) Pearson correlation scores for phenotypes from strains isolated from roots (n = 96) and soil (n = 96).

Multiple other genome loci are strongly correlated with pathogen suppression (Figure 3A, Figure S35), including chitinases (83) and the extracellular metalloprotease AprA (84). Phenotypically, extracellular protease activity also positively correlates with suppression. The BGC that correlated most strongly with *S. scabies* suppression was Pep1 (“Peptide 1”), while the related Pep2 also correlated strongly (Figure 3A). These were identified by antiSMASH as putative “bacteriocin” BGCs, and encode short DUF2282 peptides alongside DUF692 and DUF2063 proteins (Figure S9). The DUF692 protein family includes dioxygenases involved in methanobactin (85) and 3-thiaglutamate (86) biosynthesis. Other studies indicate that DUF692 and DUF2063 proteins may be involved in heavy metal and/or oxidative stress responses (87–89). Further work is required to determine the significance of both the Pep BGCs and the accessory genome loci for pathogen inhibition.

### The effect of irrigation on the soil *Pseudomonas* population

Irrigation is currently the only effective way to control potato scab, so we hypothesised that this may lead to an increase in the number of inhibitory bacteria associated with the soil and/or tuber, especially as the *Pseudomonadales* population moderately increased in irrigated soil (Figure 1C). However, a greater number of strongly suppressive strains (inhibition score ≥ 2) were isolated from non-irrigated sites (7/60 strains) than from irrigated sites (1/60 strains). A similar pattern was observed for strongly motile (score ≥ 2) strains (6 non-irrigated versus 2 irrigated). Analysis of the BGCs in our sequenced strains revealed a similar result, where 5/18 unirrigated strains contained CLP BGCs versus 0/16 irrigated strains. This counter-intuitive observation led us to hypothesise that irrigation enables non-motile, non-suppressive bacteria to survive and colonise plant roots, whereas highly motile bacteria that produce multiple biological weapons can more effectively colonise plants in drier, more ‘hostile’ conditions.

To test these hypotheses, we sampled irrigated and unirrigated sites in a neighbouring field two years after the first sampling event. 48 strains were isolated from bulk soil and the rhizospheres of tuber-forming potato plants, with and without irrigation, providing a total of 192 *P. fluorescens* strains (Supporting Dataset 1). These strains were scored for motility, HCN production and *S. scabies* suppression (Figure 8A). Our results were in strong agreement with the first sample set, including strong positive correlations between *S. scabies* inhibition, motility and HCN production (Figure 8B). A negative correlation was observed between irrigation and *S. scabies* suppression on the plant roots, but not in the surrounding soil. This appeared to be driven primarily by differences in the unirrigated samples, where a substantially greater proportion of suppressive isolates were associated with roots than with the surrounding soil. We observed a strong positive correlation between motility and root association for unirrigated samples, while the reverse was true for irrigated plants (Figure 8A). This effect of irrigation on the distribution of motile bacteria was striking – in dry plants the motile population was almost entirely associated with roots, while in irrigated plants a comparable proportion of motile bacteria were found in the soil and roots (Figure 8A).

This analysis therefore supports the root colonisation hypothesis, where a lack of irrigation leads to a more specialized pseudomonad population colonising the root. Upon irrigation, the difference between the bulk soil and root pseudomonad populations is much less significant. The mechanism for this population change is not yet defined and these changes are counter-intuitive in relation to the suppression of potato scab upon irrigation, given there is a drop in suppressive strains colonising the potato root following irrigation. Irrigation did lead to moderately more motile pseudomonads in bulk soil versus unirrigated conditions, but this was not associated with more suppressive strains or HCN producers (Figure 8A, 8B). The mechanism and significance of this irrigation effect requires further investigation. It is possible that a protective microbiome in irrigated conditions actually contains a mixture of *S. scabies*-suppressive “biocontrol” *Pseudomonas* strains alongside other non-motile pseudomonads that interact with the plant in important ways due to traits usually absent from the “biocontrol” strains, such as their ability to produce PQQ and catabolise auxins (55, 57). Profound irrigation-associated changes in antibiotic-producing *Pseudomonas* populations have previously been observed for the wheat rhizosphere (90, 91).

## Discussion

Prior studies on the suppression of potato scab have indicated a potential biocontrol role for *Pseudomonas* bacteria (14, 25, 28, 29). Fluorescent pseudomonads form multiple beneficial relationships with plants, including growth promotion and biocontrol (7, 9). However, there is limited understanding of the genetic factors that are critical for such activity, and little is known about the diversity of the *P. fluorescens* species group within a given agricultural field or how this population is shaped by environmental changes. In this study, we integrated genomics, metabolomics, phenotypic analysis, molecular biology and *in planta* assays to identify the genetic determinants of *Pseudomonas* antagonism towards *S. scabies*. This population level approach shows that the *P. fluorescens* population in a single field is highly complex, heterogeneous and dynamic (Figures 2, 3 and 8), where the overall genotypic diversity is similar to the global diversity of *P. fluorescens* (23). Pan-genome analysis and metagenomics represent increasingly powerful routes to understanding the genetic determinants of biological activity in plant-associated microbes (10, 22, 92–95).

Multiple BGCs and accessory genome loci were identified that correlated with on-plate inhibition of *S. scabies* growth and development (Figure 3), including BGCs for CLPs and HCN. These loci also correlated with inhibition of *P. infestans* and *G. graminis*, and their contribution to suppression was validated genetically. This confirmed a role for both molecules in *S. scabies* inhibition (Figures 5 and 6), representing a new function for these *Pseudomonas* specialized metabolites. Co-culture assays and cryo-SEM imaging (Figure 6) showed that HCN and a tensin-like CLP produced by Ps619 arrest the formation of streptomycete aerial hyphae and subsequent sporulation, providing the pseudomonad with a competitive advantage at the microbial interface.

*In planta* experiments confirmed that Ps619 could suppress potato scab and that CLP production was a key determinant of this inhibitory effect (Figure 7). In contrast, HCN production was not a requirement for potato scab suppression by Ps619. It is possible that HCN is not produced in sufficient amounts during root colonisation for *S. scabies* inhibition, or that it instead has an alternative natural role, such as metal chelation (96). The roles of the CLPs are reminiscent of the interaction between *B. subtilis* and *S. coelicolor*, where the CLP surfactin functions as a surfactant required for the formation of aerial structures in *B. subtilis* and arrests aerial development in *S. coelicolor* (63). Collectively, these results are surprising given that streptomycetes themselves use surfactants to assist in the erection of aerial mycelia (97, 98) and points to secondary antagonistic roles for these molecules beyond the reduction of surface tension. This is strongly supported by the inhibitory effect of purified viscosin I towards *S. scabies* (Figure 5C).

HCN and CLPs have also been associated with insect (27) and nematode (76) killing, as well as the suppression of pathogenic fungi (26, 99). This indicates that a subset of pseudomonads are generalist suppressors of pathogens (and presumably also non-pathogenic organisms) due to the production of these broad range antimicrobials. Genetic analysis indicates that these strains are more likely to produce multiple suppressive metabolites and proteins. Evidence for this is provided by the inhibition of both *P. infestans* and *G. graminis* by Ps619 Δ*hcn*Δ*ten* (Figure S35). A study of the inhibitory properties of bacteria associated with the *Arabidopsis* leaf microbiome showed that a large proportion of the total inhibitory activity was due to *Pseudomonadales* bacteria and that a subset of individual strains were active against a wide array of bacteria (100).

Unexpectedly, irrigation led to a decrease in the proportion of suppressive pseudomonads on potato roots (Figure 8A) even though irrigation is one of the most effective ways to suppress potato scab. One possible reason for this discrepancy is that irrigation enables non-suppressive *Pseudomonas* spp. with low motility to be transported to plant roots more effectively. Recruitment of non-suppressive pseudomonads to the rhizosphere may benefit the plant in other ways, such as immune system priming (101, 102) or modulation of auxin biosynthesis. For example, Cheng *et al.* (8) showed that auxin biosynthesis was linked to plant growth promotion and induced systemic resistance by *P. fluorescens* Pf.SS101. An alternative hypothesis is that changes in the overall relative abundance of soil *Pseudomonas* over *Streptomyces* resulting from irrigation may override the observed shift towards less-suppressive *Pseudomonas* genotypes. In support of this, drought-induced enrichment for commensal *Streptomyces* and depletion of *Proteobacteria* in sorghum and rice plants have been shown to be reversed by irrigation (103, 104). In this model, irrigation may reduce the relative fitness of *S. scabies* versus *Pseudomonas* spp., while the microbiome of irrigated roots simultaneously becomes less optimal for disease suppression.

Our data show that Ps619 is highly effective at inhibiting potato scab, yet Ps619-like strains are naturally less abundant in irrigated conditions. Therefore, possible future efforts to control potato scab could combine irrigation with pre-treatment with effective biocontrol strains, like Ps619, to ensure tubers are colonized by a significant proportion of biocontrol strains. Such a strategy could reduce the quantity of water required for effective scab suppression. While our study was focused on fluorescent pseudomonads, interactions between these bacteria and the wider microbiome (Figure 1) may also have a key role in potato scab suppression.

Moving forward, systematic analyses of individual organisms within microbiomes will continue to help answer questions relating to microbial communities and host interactions that are difficult to address using global ‘omics approaches alone. For example, the role of many bacterial specialized metabolites in nature is poorly understood, especially for prolific producers such as the pseudomonads and the streptomycetes (105). Future studies could examine whether the host selects for bacterial populations enriched in specific BGCs and whether environmental stimuli modulate the abundance of these BGCs. Synthetic microbial communities based on well-characterized natural communities could then be used to test hypotheses on the role of specialized metabolites in shaping the community or modulating the health of the host organism.

## Materials and Methods

### Strains and growth conditions

All strains used in this study are listed in Table S1. Unless otherwise stated, chemicals were purchased from Sigma-Aldrich, enzymes from New England Biolabs, and molecular biology kits from GE Healthcare and Promega. All *P. fluorescens* strains were grown at 28 °C in L medium (Luria base broth, Formedium) and *Escherichia coli* at 37 °C in lysogeny broth (LB) (106). 1.3% agar was added for solid media. Gentamicin was used at 25 μg/mL, carbenicillin at 100 μg/mL and tetracycline (Tet) at 12.5 μg/mL. *S. scabies* spore suspensions were prepared using established procedures (107).

### Soil sample collection

Soil samples were collected from potato fields at RG Abrey Farms (East Wretham, Norfolk, U.K., 52.4644° N, 0.8299° E). The first sampling was conducted in 2015 from two adjacent plots in a single field. Soil samples were taken on 22^nd^ January 2015, immediately prior to planting. One plot was then covered loosely in polythene to protect it from irrigation. The same field sites were sampled again in May at the point of maximum scab impact, once potato tubers had begun to form. In this case soil samples were taken from the base of the plants, near the root system. For each sampling event, a total of 12 samples were taken from three parallel potato beds at regularly spaced intervals approximately one metre apart. For the second experiment, 12 irrigated and 12 non-irrigated potato plants were uprooted from field sites in June 2017 and returned to the laboratory in large pots. Bulk soil samples were taken from these pots alongside an equivalent number of rhizosphere-associated samples, which were defined as isolated root systems gently shaken to remove bulk soil before processing as below. Samples were collected in sterile 50 mL tubes and stored at 4 °C.

### Isolation of soil *Pseudomonas*

Sample processing was conducted at 4 °C throughout. 10 mL of sterile phosphate-buffered saline (PBS, per litre: 8 g NaCl, 0.2 g KCl, 1.44 g Na_2_HPO_4_, 0.24 g KH_2_PO_4_, pH 7.4) were added to 50 mL tubes containing 20 g of soil or root material, and vortexed vigorously for 10 minutes. Samples were then filtered through a sterile muslin filter to remove larger debris. The resulting suspension of soil and organic matter was centrifuged at 1000 rpm for 30 s to pellet remaining soil particles, before serial dilution in PBS and plating on *Pseudomonas* selective agar. The selection media comprised *Pseudomonas* agar base (Oxoid, UK) supplemented with CFC (cetrimide/fucidin/cephalosporin) *Pseudomonas* selective supplement (Oxoid, UK). Plates were incubated at 28 °C until colonies arose, then isolated single colonies were patched on fresh CFC agar and incubated overnight at 28 °C before streaking to single colonies on King’s B (KB) agar plates (108). Six isolates were selected at random per soil sample and subjected to phenotypic/genomic analysis.

### Amplicon sequencing

Genomic DNA was isolated from 3 g of pooled soil samples using the FastDNA™ SPIN Kit for soil (MP Biomedicals, UK) following the manufacturer’s instructions. Genomic DNA concentration and purity was determined by NanoDrop spectrophotometry as above. Microbial 16S rRNA genes were amplified from soil DNA samples with barcoded universal prokaryotic primers (F515/R806) targeting the V4 region(109), and then subjected to Illumina® MiSeq sequencing (600-cycle, 2×300 bp) at the DNA Sequencing Facility, Department of Biochemistry, University of Cambridge (UK). The data was analysed using the MiSeq Reporter Metagenomics Workflow (Illumina, UK) to acquire read counts for all taxonomic ranks from phylum to genus. MiSeq data were visualized and analysed using Degust 3.1.0 (http://degust.erc.monash.edu/) and Pheatmap (https://CRAN.R-project.org/package=pheatmap) in R 3.5.1.

### Phenotypic assays

All phenotyping assays were conducted at least twice independently, and where disagreements were recorded in the ordinal data, additional repeats were conducted until a firm consensus was reached.

*Swarming motility:* 0.5% KB agar plates were poured and allowed to set and dry for 1 hour in a sterile flow chamber. Plates were then inoculated with 2 μL spots of overnight cultures and incubated overnight at room temperature. The motility of each isolate was tested in triplicate and scored from 0 (no motility) to 3 (high motility).

*Secreted protease activity:* 5 μL of overnight cultures were spotted onto KB plates containing 1.0% skimmed milk powder. Plates were incubated at 28 °C and photographed after 24 hours, with individual isolates scored as protease positive (score = 2) or negative (score = 0).

*Hydrogen cyanide (HCN) production:* An adaptation of the method described in Castric *et al.* (110) was used. *Pseudomonas* isolates were inoculated into 150 μL of liquid KB medium in individual wells of a flat bottomed 96-well plate. The plates were then overlaid with Feigl-Anger reagent paper (79), prepared as follows. Whatman 3MM chromatography paper was soaked in Feigl-Anger detection reagent (5 mg/mL copper(II) ethyl acetoacetate and 5 mg/mL 4,4’-methylenebis(N,N-dimethylaniline) dissolved in chloroform). After complete solvent evaporation, the paper was placed under the plate lid and strains were grown at 28 °C overnight with gentle shaking. The intensity of blue staining on the paper was then scored from 0 (no colour) to 3 (high blue intensity). The same method was applied to isolates growing on agar medium.

*S. scabies inhibition:* Two parallel lines of *S. scabies* 87-22 spores were streaked onto SFM plates (107) using a sterile toothpick. These lines were then cross streaked with overnight cultures of *Pseudomonas* isolates. Plates were incubated at 30 °C and the relative performance of each species was assessed daily for 5 days.

*P. infestans inhibition:* Assays were conducted with *P. infestans* #2006-3920A (6-A1) (The Sainsbury Laboratory, UK). This was maintained on rye agar medium supplemented with 2% sucrose (C-RSA) (111) at 21 °C. C-RSA was filtered through muslin fabric to enable clearer observation of oomycete growth. Three 10 μL drops of overnight cultures per *Pseudomonas* isolate strain were placed equidistantly 15 mm from the edge of C-RSA plates. A 3 mm plug from the leading edge of a *P. infestans* culture was then placed in the centre of each plate and incubated for a further 7 days at 21 °C before scoring and imaging.

*Take-all inhibition:* Assays were conducted with *Gaeumannomyces graminis var. tritici* strain NZ.66.12 (*Ggt*) (17). Three 10 μL drops of overnight *Pseudomonas* cultures per strain were placed equidistantly 15 mm from the edge of potato dextrose agar (PDA) plates and incubated for 24 hours at 28 °C. *Ggt* NZ.66.12 was cultured on PDA agar for five days at room temperature. A 3 mm plug from the leading edge of the NZ.66.12 culture was then placed in the centre of each plate and incubated for a further five days at 22 °C before the extent of *Ggt* inhibition was assessed.

### DNA extraction and Illumina® genome sequencing

Single colonies of each isolate to be sequenced were picked from L agar plates and grown overnight in L medium. DNA was then extracted from 2 mL of cell culture using a GenElute™ Bacterial Genomic DNA Kit (Sigma-Aldrich, USA). DNA samples were subjected to an initial quality check using a Nanodrop spectrophotometer (Thermo Scientific, Wilmington, DE, USA) before submission for Nextera library preparation and paired-end read sequencing on the Illumina® MiSeq platform (600­cycle, 2×300 bp) at the DNA Sequencing Facility, Department of Biochemistry, University of Cambridge (UK). Reads from 35 pseudomonads collected in February 2015 were assembled into genomes using MaSuRCA v3.2.6 (112) with the following settings:

GRAPH_KMER_SIZE=auto; USE_LINKING_MATES=1; LIMIT_JUMP_COVERAGE=60;

CA_PARAMETERS = ovlMerSize=30 cgwErrorRate=0.25 ovlMemory=4GB;

NUM_THREADS=16; JF_SIZE=100000000; DO_HOMOPOLYMER_TRIM=0.

Reads from 32 samples collected in May 2015 were assembled into genomes using SPAdes v3.6.2 (113) with k-mer flag set to -k 21,33,55,77,99,127. All assembly tasks were conducted using 16 CPUs on a 256 GB compute node within the Norwich Bioscience Institutes (NBI) High Performance Computing cluster. An additional strain from May 2015 (Ps925) was sequenced and assembled by MicrobesNG (http://www.microbesng.uk), which is supported by the BBSRC (grant number BB/L024209/1). The 69 assembled genome sequences were annotated using Prokka (114), which implements Prodigal (115) as an open reading frame calling tool. Assembly qualities were assessed using CheckM (116). Genome assemblies are available at the European Nucleotide Archive (http://www.ebi.ac.uk/ena/) with the project accession PRJEB34261.

### Phylogenetic and bioinformatic analysis

The *gyrB* housekeeping gene sequence was identified in each newly sequenced genome by BLAST comparison with the sequence of *gyrB* from *P. fluorescens* SBW25. The full-length *gyrB* sequences from these strains and several reference strains were aligned using MUSCLE 3.8.31 (117) with default settings, then a maximum likelihood tree was calculated using RAxML 8.2.12 (118) on the CIPRES portal (119) with the following parameters: raxmlHPC-HYBRID-AVX -T 4 -f a -N autoMRE -n result -s infile.txt -c 25 -m GTRCAT -p 12345 -k -x 12345. Genomes were subjected to bioinformatic analysis as described in the Supplementary Information. Phylogenetic trees and presence/absence data for accessory genes were visualised using Interactive Tree of Life (iTOL) (120), with *Pseudomonas aeruginosa* PAO1 *gyrB* as the outgroup.

### Molecular biology procedures

Cloning was carried out in accordance with standard molecular biology techniques. *P. fluorescens* deletion mutants were constructed by allelic exchange as described previously (121). Up- and downstream flanking regions (approx. 500 bp) to the target genes were amplified using primers listed in Table S2. PCR products in each case were ligated into pTS1 (122) between XhoI and BamHI. The resulting deletion vectors were transformed into the target strains by electroporation, and single crossovers selected on L + Tet and re-streaked to isolate single colonies. 100 mL cultures in L medium from single crossovers were grown overnight at 28 °C, then plated onto L + 10% sucrose plates to counter-select for double crossovers. Individual colonies from these plates were then patched onto L plates ± Tet, with Tet-sensitive colonies tested for gene deletion by colony PCR using primers external to the deleted gene in each case (Table S2).

Luminescent-tagged strains were produced by introduction of the *Aliivibrio fischeri luxCDABE* cassette into the neutral *att::Tn7* site in *Pseudomonas* chromosomes using the Tn7-based expression system described in Choi *et al.* (74). Strains were electroporated with plasmids pUC18-mini*Tn7*T-Gm-*lux* and the helper pTNS2, and transformant colonies were grown on solid L medium + gentamicin for 2-3 days at 28 °C. Integration of the *lux* cassette into *Pseudomonas* genomes was confirmed by PCR and with a luminometer. Luminescent cells were then tracked using the NightOWL visualisation system (Berthold Technologies, Germany). All plasmids used in this study are reported in Table S3.

### Liquid chromatography - mass spectrometry detection of lipopeptides

*Pseudomonas* isolates were grown overnight in L medium (10 mL) for 16 hours at 28 °C. 100 μL of each culture was used to inoculate 40 mm diameter KB agar plates. Plates were incubated for 24 hours at 28 °C, before the agar from each plate was decanted into a sterile 50 mL tube and extracted with 10 mL 50% EtOH with occasional vortexing for 3 hours. 2 mL was taken from each sample and centrifuged in 2 mL tubes for 5 min at 16,000 g. The supernatant was collected and stored at −80 °C. Samples were diluted with an equal volume of water, then subjected to LC-MS analysis using a Shimadzu Nexera X2 UHPLC coupled to a Shimadzu ion-trap time-of-flight (IT-TOF) mass spectrometer. Samples (5 μL) were injected onto a Phenomenex Kinetex 2.6 μm XB-C18 column (50 x 2.1mm, 100 Å), eluting with a linear gradient of 5 to 95% acetonitrile in water + 0.1% formic acid over 6 minutes with a flow-rate of 0.6 mL/min at 40 °C. To compare the retention times of viscosin I (Ps682), WLIP (*Pseudomonas* sp. LMG 2338) and viscosin (*P. fluorescens* SBW25), extracts were prepared from their producing organisms as described above. The same chromatography conditions as above were used, but with a linear gradient of 5 to 100% acetonitrile in water + 0.1% formic acid over 15 minutes.

Positive mode mass spectrometry data was collected between *m/z* 300 and 2000 with an ion accumulation time of 10 ms featuring an automatic sensitivity control of 70% of the base peak. The curved desolvation line temperature was 300 °C and the heat block temperature was 250 °C. MS/MS data was collected in a data-dependent manner using collision-induced dissociation energy of 50% and a precursor ion width of 3 Da. The instrument was calibrated using sodium trifluoroacetate cluster ions prior to every run.

A molecular network was created using the online workflow at the Global Natural Product Social Molecular Networking (GNPS) site (https://gnps.ucsd.edu/) (72). The data was filtered by removing all MS/MS peaks within +/- 17 Da of the precursor *m/z*. The data was then clustered with MS-Cluster with a parent mass tolerance of 1 Da and a MS/MS fragment ion tolerance of 0.5 Da to create consensus spectra. Consensus spectra that contained less than 2 spectra were discarded. A network was then created where edges were filtered to have a cosine score above 0.6 and more than 4 matched peaks. Further edges between two nodes were kept in the network if each of the nodes appeared in each other’s respective top 10 most similar nodes. The spectra in the network were then searched against GNPS spectral libraries. The library spectra were filtered in the same manner as the input data. All matches kept between network spectra and library spectra were required to have a score above 0.7 and at least 4 matched peaks. Networks were visualised using Cytoscape v3.8.2 (123) and the data was manually filtered to remove duplicate nodes (same *m/z* and retention time). The data is available as a MassIVE dataset at ftp://massive.ucsd.edu/MSV000084283/ and the GNPS analysis is available here: https://gnps.ucsd.edu/ProteoSAFe/status.jsp?task=51ac5fe596424cf88cfc17898985cac2

High-resolution mass spectra were acquired on a Synapt G2-Si mass spectrometer equipped with an Acquity UPLC (Waters). Aliquots of the samples were injected onto an Acquity UPLC® BEH C18 column, 1.7 μm, 1×100 mm (Waters) and eluted with a gradient of acetonitrile/0.1% formic acid (B) in water/0.1% formic acid (A) with a flow rate of 0.08 mL/min at 45 °C. The concentration of B was kept at 1% for 1 min followed by a gradient up to 40% B in 9 min, ramping to 99% B in 1 min, kept at 99% B for 2 min and re-equilibrated at 1% B for 4 min. MS data were collected in positive mode with the following parameters: resolution mode, positive ion mode, scan time 0.5 s, mass range *m/z* 50-1200 calibrated with sodium formate, capillary voltage = 2.5 kV; cone voltage = 40 V; source temperature = 125 °C; desolvation temperature = 300 °C. Leu-enkephalin peptide was used to generate a lock-mass calibration with 556.2766, measured every 30 s during the run. For MS/MS fragmentation, a data directed analysis (DDA) method was used with the following parameters: precursor selected from the 4 most intense ions; MS2 threshold: 5,000; scan time 0.5 s; no dynamic exclusion. In positive mode, collision energy (CE) was ramped between 10-30 at low mass (*m/z* 50) and 15-60 at high mass (*m/z* 1200).

### Purification and characterisation of viscosin I

Pre-cultures of Ps682 were grown in 10 mL LB medium for 16 hours and 600 μL aliquots were used to inoculate 140 mm diameter KB agar plates. Fifteen plates were inoculated and incubated for 24 hours. The agar was decanted and extracted with 500 mL ethyl acetate for 2 hours with occasional mixing. The organic fraction was filtered off, washed with 3 x 200 mL water, dried over MgSO_4_, and then solvent was removed *in vacuo*. The resulting material was dissolved in MeOH and applied to a 12 g C18 flash chromatography column (Biotage), pre-equilibrated in 70% MeOH. Separation proceeded by a gradient of 70-100% MeOH over 10 column volumes. Each fraction was subject to LC-MS analysis using a Shimadzu Nexera X2 UHPLC coupled to a Shimadzu IT-TOF mass spectrometer, as described above.

Solvent was removed from viscosin I-containing fractions using a rotary evaporator and then a Genevac (SP Scientific). Fractions were then dissolved in MeOH to 1 mg/mL and further purified using a Thermo Dionex Ultimate 3000 HPLC system. 200 μL aliquots were injected onto a Phenomenex C18 Luna column (5 μm, 250 mm x 10 mm) and eluted with a linear gradient of 5% to 95% acetonitrile/H_2_O over 30 minutes, with a flow rate of 4 mL/min and UV absorption data collected at 210 nm. LC-MS analysis (as described above) was used to identify pure fractions, which were combined and dried *in vacuo* to yield 1.4 mg viscosin I as a white powder.

Viscosin I (1.4 mg) was dissolved in *N,N*-dimethylformamide-d_7_ (DMF-d_7_) and nuclear magnetic resonance (NMR) spectra were acquired on a Bruker Avance Neo 600 MHz spectrometer equipped with a TCI cryoprobe. The experiments were carried out at 298 K with the residual DMF solvent used as an internal standard (δ_H_/δ_C_ 2.75/29.76). The residual solvent signal from H_2_O was suppressed through a presaturation sequence in 1D ^1^H. Resonances were assigned through 1D ^1^H and DEPT135 experiments, and 2D COSY, HSQCed, HMBC, TOCSY and HSQC-TOCSY experiments. Spectra were analysed using Bruker TopSpin 3.5 and Mestrelab Research Mnova 14.0 software. NMR data are reported in Figures S19-S29 and Table S4.

### Disk Diffusion Assays

A *S. scabies* 87-22 spore suspension was diluted 1:100 in sterile MQ water and 60 μL aliquots were applied to instant potato medium (20 g/L Smash™ Instant Mash, 20 g/L agar) on 100 mm square plates. The spore solution was evenly distributed using a sterile cotton bud and the plate was dried for 30 min. Viscosin I was diluted in MeOH to produce a range of concentrations from 20 to 100 μg/mL. Each concentration was applied to a 6 mm filter paper disk, in 5 x 20 μL applications at 10 min intervals, and then dried for 30 min. The disks were then applied to the surface of the agar plate, which was incubated at 30 °C and imaged daily.

### Scanning electron microscopy

Small pieces of the *Pseudomonas-Streptomyces* co-culture samples were excised from the surface of agar plates and mounted on an aluminium stub using Tissue Tek^R^ (BDH Laboratory Supplies, Poole, England). The stub was then immediately plunged into liquid nitrogen slush at approximately −210 °C to cryo-preserve the material. The sample was transferred onto the cryostage of an ALTO 2500 cryo-transfer system (Gatan, Oxford, England) attached to an FEI Nova NanoSEM 450 (FEI, Eindhoven, The Netherlands). Sublimation of surface frost was performed at −95 °C for three minutes before sputter coating the sample with platinum for 3 min at 10 mA, at colder than −110 °C. After sputter-coating, the sample was moved onto the cryo-stage in the main chamber of the microscope, held at −125 °C. The sample was imaged at 3 kV and digital TIFF files were stored.

### Potato scab biocontrol assays

*In planta* assays were performed as described previously (124, 125) with some modifications. Briefly, 2 L of GYM (4 g/L glucose, 4 g/L yeast extract, 10 g/L malt extract, 2 g/L CaCO_3_, pH 7.2) was inoculated with a *S. scabies* 87-22 starter culture from a spore suspension and incubated for 48 h at 30 °C, 250 rpm. Each culture was then centrifuge at 16,994 x *g* for 15 min and washed twice with PBS (2 L). *Pseudomonas* strains Ps619, Ps682 and associated mutants were grown overnight at 28 °C, 250 rpm in 50 mL L medium (Luria base broth, Formedium), centrifuged at 1,520 x *g* for 15 min, washed twice with PBS (20 mL) and adjusted to OD600 = 0.2 for the final inoculation.

50 mL of autoclaved vermiculite and 50 mL of bacterial culture were mixed to constitute the final inoculum. 5 L pots were filled with steam-sterilised substrate (John Innes Cereal Mix) and inoculum was applied into the pots and mixed with the soil. Different combinations of bacterial inocula were made accordingly following the same method. Potato seeds cv. Maris Piper obtained from VCS Potatoes Ltd (Suffolk, UK) were surfaced disinfected by immersion in 1% sodium hypochlorite for 15 min. Tubers were then rinsed with water, air-dried and placed in pots. Pots were watered until saturation according to their growth stage. Potato plants were grown in a glasshouse with a light cycle of 16h/8h at 18-20 °C. Two independent experiments were run between 17^th^ July - 6th November 2020 and 31^st^ July - 20th November 2020, and three - four plants were used per treatment. Tubers were collected, washed, weighed and scored accordingly to a one to six scale as described in Andrade *et al*. (81). Potato plants were dried for four days at 30 °C and aerial parts and tuber weights were both recorded, as well as tuber number. Treatment differences were carried out based on the disease index (DI) of each plant (n=4 for each treatment). The DI was calculated as the mean of all the scored tubers per plant and *p* values were calculated using Dunnett’s multiple comparison test.

## Acknowledgements

Financial support was provided by a Royal Society University Research Fellowship to Andrew Truman, Biotechnology and Biological Sciences Research Council (BBSRC) BIO, MET, PH and MfN Institute Strategic Programme grants to the John Innes Centre (JIC), a JIC Institute Development Grant and NPRONET Proof of Concept grant. Jonathan Ford was supported by a BBSRC DTP studentship and Alba Pacheco-Moreno was supported by a BBSRC iCASE PhD studentship, both awarded to the Norwich Research Park. We thank Prof. Jonathan Jones (The Sainsbury Laboratory) for providing *P. infestans* isolates and Dr Tim Mauchline (Rothamsted Research) for providing *Ggt* NZ.66.12. We thank the JIC Bioimaging, Metabolomics and NMR facilities for their contribution to this publication, Dr Carlo de Oliveira Martins (JIC) for assistance with mass spectrometry, and Dr Natalia Miguel-Vior (JIC) for assistance with potato scab biocontrol assays.

## References

1. Larkin, R. P., Honeycutt, C. W., Griffin, T. S., Olanya, O. M., Halloran, J. M. and He, Z. (2011) Effects of different potato cropping system approaches and water management on soilborne diseases and soil microbial communities. Phytopathology 101, 58–67. DOI: 10.1094/PHYTO-04-10-0100.

2. Lerat, S., Simao-Beaunoir, A.-M. snd Beaulieu, C. (2009) Genetic and physiological determinants of *Streptomyces scabies* pathogenicity. Molecular Plant Pathology 10, 579–585. DOI: 10.1111/j.1364-3703.2009.00561.x.

3. Bignell, D. R. D., Huguet-Tapia, J. C., Joshi, M. V., Pettis, G. S. and Loria, R. (2010) What does it take to be a plant pathogen: genomic insights from *Streptomyces* species. Antonie van Leeuwenhoek 98, 179–194. DOI: 10.1007/s10482-010-9429-1.

4. Weller, D. M. (2007) *Pseudomonas* biocontrol agents of soilborne pathogens: looking back over 30 years. Phytopathology 97, 250–256. DOI: 10.1094/PHYTO-97-2-0250.

5. Köhl, J., Kolnaar, R. and Ravensberg, W. J. (2019) Mode of Action of Microbial Biological Control Agents Against Plant Diseases: Relevance Beyond Efficacy. Front Plant Sci 10. DOI: 10.3389/fpls.2019.00845.

6. Loper, J. E., Hassan, K. A., Mavrodi, D. V., Davis, E. W., Lim, C. K., Shaffer, B. T., Elbourne, L. D. H., Stockwell, V. O., Hartney, S. L., Breakwell, K., Henkels, M. D., Tetu, S. G., Rangel, L. I., Kidarsa, T. A., Wilson, N. L., van de Mortel, J. E., Song, C., Blumhagen, R., Radune, D., Hostetler, J. B., Brinkac, L. M., Durkin, A. S., Kluepfel, D. A., Wechter, W. P., Anderson, A. J., Kim, Y. C., Pierson, L. S., Pierson, E. A., Lindow, S. E., Kobayashi, D. Y., Raaijmakers, J. M., Weller, D. M., Thomashow, L. S., Allen, A. E. and Paulsen, I. T. (2012) Comparative Genomics of Plant-Associated *Pseudomonas* spp.: Insights into Diversity and Inheritance of Traits Involved in Multitrophic Interactions. PLoS Genet 8, e1002784. DOI: 10.1371/journal.pgen.1002784.s027.

7. Zamioudis, C., Mastranesti, P., Dhonukshe, P., Blilou, I. and Pieterse, C. M. J. (2013) Unraveling Root Developmental Programs Initiated by Beneficial *Pseudomonas* spp. Bacteria. Plant Physiol. 162, 304–318. DOI: 10.1104/pp.112.212597.

8. Cheng, X., Etalo, D. W., van de Mortel, J. E., Dekkers, E., Nguyen, L., Medema, M. H. and Raaijmakers, J. M. (2017) Genome-wide analysis of bacterial determinants of plant growth promotion and induced systemic resistance by *Pseudomonas fluorescens*. Environmental Microbiology 19, 4638–4656. DOI: 10.1111/1462-2920.13927.

9. Haas, D. and Défago, G. (2005) Biological control of soil-borne pathogens by fluorescent pseudomonads. Nat Rev Micro 3, 307–319. DOI: 10.1038/nrmicro1129.

10. Biessy, A., Novinscak, A., Blom, J., Léger, G., Thomashow, L. S., Cazorla, F. M., Josic, D. and Filion, M. (2019) Diversity of phytobeneficial traits revealed by whole-genome analysis of worldwide-isolated phenazine-producing *Pseudomonas* spp. Environmental Microbiology 21, 437–455. DOI: 10.1111/1462-2920.14476.

11. Gross, H. and Loper, J. E. (2009) Genomics of secondary metabolite production by *Pseudomonas* spp. Nat. Prod. Rep. 26, 1408–1446. DOI: 10.1039/b817075b.

12. Nguyen, D. D., Melnik, A. V., Koyama, N., Lu, X., Schorn, M., Fang, J., Aguinaldo, K., Lincecum, T. L., Jr, Ghequire, M. G. K., Carrión, V. J., Cheng, T. L., Duggan, B. M., Malone, J. G., Mauchline, T. H., Sanchez, L. M., Kilpatrick, A. M., Raaijmakers, J. M., De Mot, R., Moore, B. S., Medema, M. H. and Dorrestein, P. C. (2016) Indexing the *Pseudomonas* specialized metabolome enabled the discovery of poaeamide B and the bananamides. Nature Microbiology 2, 1–9. DOI: 10.1038/nmicrobiol.2016.197.

13. Stringlis, I. A., Zhang, H., Pieterse, C. M. J., Bolton, M. D. and de Jonge, R. (2018) Microbial small molecules - weapons of plant subversion. Nat. Prod. Rep. 35, 410–433. DOI: 10.1039/c7np00062f.

14. Arseneault, T., Goyer, C. and Filion, M. (2013) Phenazine Production by *Pseudomonas* sp. LBUM223 Contributes to the Biological Control of Potato Common Scab. Phytopathology 103, 995–1000. DOI: 10.1094/PHYTO-01-13-0022-R.

15. Ghequire, M. G. K. and De Mot, R. (2014) Ribosomally encoded antibacterial proteins and peptides from *Pseudomonas*. FEMS Microbiol Rev 38, 523–568. DOI: 10.1111/1574-6976.12079.

16. Rangel, L. I., Henkels, M. D., Shaffer, B. T., Walker, F. L., Davis, E. W., Stockwell, V. O., Bruck, D., Taylor, B. J. and Loper, J. E. (2016) Characterization of Toxin Complex Gene Clusters and Insect Toxicity of Bacteria Representing Four Subgroups of *Pseudomonas fluorescens*. PLoS ONE 11, e0161120. DOI: 10.1371/journal.pone.0161120.

17. Mauchline, T. H., Chedom-Fotso, D., Chandra, G., Samuels, T., Greenaway, N., Backhaus, A., McMillan, V., Canning, G., Powers, S. J., Hammond-Kosack, K. E., Hirsch, P. R., Clark, I. M., Mehrabi, Z., Roworth, J., Burnell, J. and Malone, J. G. (2015) An analysis of *Pseudomonas* genomic diversity in take-all infected wheat fields reveals the lasting impact of wheat cultivars on the soil microbiota. Environmental Microbiology 17, 4764–4778. DOI: 10.1111/1462-2920.13038.

18. Wei, Z., Gu, Y., Friman, V.-P., Kowalchuk, G. A., Xu, Y., Shen, Q. and Jousset, A. (2019) Initial soil microbiome composition and functioning predetermine future plant health. Sci Adv 5, eaaw0759. DOI: 10.1126/sciadv.aaw0759.

19. Kwak, Y.-S. and Weller, D. M. (2013) Take-all of Wheat and Natural Disease Suppression: A Review. Plant Pathol. J. 29, 125–135. DOI: 10.5423/PPJ.SI.07.2012.0112.

20. Silby, M. W., Cerdeño-Tárraga, A. M., Vernikos, G. S., Giddens, S. R., Jackson, R. W., Preston, G. M., Zhang, X.-X., Moon, C. D., Gehrig, S. M., Godfrey, S. A. C., Knight, C. G., Malone, J. G., Robinson, Z., Spiers, A. J., Harris, S., Challis, G. L., Yaxley, A. M., Harris, D., Seeger, K., Murphy, L., Rutter, S., Squares, R., Quail, M. A., Saunders, E., Mavromatis, K., Brettin, T. S., Bentley, S. D., Hothersall, J., Stephens, E., Thomas, C. M., Parkhill, J., Levy, S. B., Rainey, P. B. and Thomson, N. R. (2009) Genomic and genetic analyses of diversity and plant interactions of *Pseudomonas fluorescens*. Genome Biol. 10, R51. DOI: 10.1186/gb-2009-10-5-r51.

21. Gomila, M., Peña, A., Mulet, M., Lalucat, J. and García-Valdés, E. (2015) Phylogenomics and systematics in *Pseudomonas*. Front. Microbiol. 6, 214. DOI: 10.3389/fmicb.2015.00214.

22. Melnyk, R. A., Hossain, S. S. and Haney, C. H. (2019) Convergent gain and loss of genomic islands drive lifestyle changes in plant-associated *Pseudomonas*. ISME J. 13, 1575–1588. DOI: 10.1038/s41396-019-0372-5.

23. Garrido-Sanz, D., Meier-Kolthoff, J. P., Göker, M., Martín, M., Rivilla, R. and Redondo-Nieto, M. (2016) Genomic and Genetic Diversity within the *Pseudomonas fluorescens* Complex. PLoS ONE 11, e0150183. DOI: 10.1371/journal.pone.0150183.

24. Little, R. H., Woodcock, S. D., Campilongo, R., Fung, R. K. Y., Heal, R., Humphries, L., Pacheco-Moreno, A., Paulusch, S., Stigliano, E., Vikeli, E., Ward, D. and Malone, J. G. (2019) Differential Regulation of Genes for Cyclic-di-GMP Metabolism Orchestrates Adaptive Changes During Rhizosphere Colonization by *Pseudomonas fluorescens*. Front. Microbiol. 10, 1089. DOI: 10.3389/fmicb.2019.01089.

25. Arseneault, T., Goyer, C. and Filion, M. (2015) *Pseudomonas fluorescens* LBUM223 Increases Potato Yield and Reduces Common Scab Symptoms in the Field. Phytopathology 105, 1311–1317. DOI: 10.1094/PHYTO-12-14-0358-R.

26. Michelsen, C. F., Watrous, J., Glaring, M. A., Kersten, R., Koyama, N., Dorrestein, P. C. and Stougaard, P. (2015) Nonribosomal peptides, key biocontrol components for *Pseudomonas fluorescens* In5, isolated from a Greenlandic suppressive soil. mBio 6, e00079. DOI: 10.1128/mBio.00079-15.

27. Flury, P., Vesga, P., Péchy-Tarr, M., Aellen, N., Dennert, F., Hofer, N., Kupferschmied, K. P., Kupferschmied, P., Metla, Z., Ma, Z., Siegfried, S., de Weert, S., Bloemberg, G., Höfte, M., Keel, C. J. and Maurhofer, M. (2017) Antimicrobial and Insecticidal: Cyclic Lipopeptides and Hydrogen Cyanide Produced by Plant-Beneficial *Pseudomonas* Strains CHA0, CMR12a, and PCL1391 Contribute to Insect Killing. Front Microbiol 8, 100. DOI: 10.3389/fmicb.2017.00100.

28. Elphinstone, J. G., Thwaites, R., Stalham, M. and Wale, S. (2009) Integration of precision irrigation and non-water based measures to suppress common scab of potato (Potato Council Project Report). https://ahdb.org.uk/integration-of-precision-irrigation-and-non-water-based-measures-to-suppress-common-scab-of-potato [accessed June 2021]

29. Rosenzweig, N., Tiedje, J. M., Quensen, J. F., III, Meng, Q. and Hao, J. J. (2012) Microbial communities associated with potato common scab-suppressive soil determined by pyrosequencing analyses. Plant Disease 96, 718–725. DOI: 10.1094/PDIS-07-11-0571.

30. Meng, Q., Yin, J., Rosenzweig, N., Douches, D. and Hao, J. J. (2012) Culture-based assessment of microbial communities in soil suppressive to potato common scab. Plant disease 96, 712–717. DOI: 10.1094/PDIS-05-11-0441.

31. Arseneault, T., Goyer, C. and Filion, M. (2016) Biocontrol of Potato Common Scab is Associated with High *Pseudomonas fluorescens* LBUM223 Populations and Phenazine-1-Carboxylic Acid Biosynthetic Transcript Accumulation in the Potato Geocaulosphere. Phytopathology 106, 963–970. DOI: 10.1094/PHYTO-01-16-0019-R.

32. Fierer, N. (2017) Embracing the unknown: disentangling the complexities of the soil microbiome. Nat Rev Micro 15, 579–590. DOI: 10.1038/nrmicro.2017.87.

33. Mauchline, T. H. and Malone, J. G. (2017) Life in earth - the root microbiome to the rescue? Curr. Opin. Microbiol. 37, 23–28. DOI: 10.1016/j.mib.2017.03.005.

34. Law, C. W., Chen, Y., Shi, W. and Smyth, G. K. (2014) voom: Precision weights unlock linear model analysis tools for RNA-seq read counts. Genome Biol. 15, R29. DOI: 10.1186/gb-2014-15-2-r29.

35. Weinert, N., Piceno, Y., Ding, G.-C., Meincke, R., Heuer, H., Berg, G., Schloter, M., Andersen, G. and Smalla, K. (2011) PhyloChip hybridization uncovered an enormous bacterial diversity in the rhizosphere of different potato cultivars: many common and few cultivar-dependent taxa. FEMS Microbiol. Ecol. 75, 497–506. DOI: 10.1111/j.1574-6941.2010.01025.x.

36. Pfeiffer, S., Mitter, B., Oswald, A., Schloter-Hai, B., Schloter, M., Declerck, S. and Sessitsch, A. (2017) Rhizosphere microbiomes of potato cultivated in the High Andes show stable and dynamic core microbiomes with different responses to plant development. FEMS Microbiol. Ecol. 93, fiw242. DOI: 10.1093/femsec/fiw242.

37. Blin, K., Shaw, S., Steinke, K., Villebro, R., Ziemert, N., Lee, S. Y., Medema, M. H. and Weber, T. (2019) antiSMASH 5.0: updates to the secondary metabolite genome mining pipeline. Nucleic Acids Res. 47, W81–W87. DOI: 10.1093/nar/gkz310.

38. Pessi, G. and Haas, D. (2000) Transcriptional control of the hydrogen cyanide biosynthetic genes hcnABC by the anaerobic regulator ANR and the quorum-sensing regulators LasR and RhlR in *Pseudomonas aeruginosa*. Journal of Bacteriology 182, 6940–6949. DOI: 10.1128/JB.182.24.6940-6949.2000.

39. Metelev, M., Serebryakova, M., Ghilarov, D., Zhao, Y. and Severinov, K. (2013) Structure of microcin B-like compounds produced by *Pseudomonas syringae* and species specificity of their antibacterial action. Journal of Bacteriology 195, 4129–4137. DOI: 10.1128/JB.00665-13.

40. Palm, C. J., Gaffney, T. and Kosuge, T. (1989) Cotranscription of genes encoding indoleacetic acid production in *Pseudomonas syringae* subsp. *savastanoi*. Journal of Bacteriology 171, 1002–1009. DOI: 10.1128/jb.171.2.1002-1009.1989.

41. McClerklin, S. A., Lee, S. G., Harper, C. P., Nwumeh, R., Jez, J. M. and Kunkel, B. N. (2018) Indole-3-acetaldehyde dehydrogenase-dependent auxin synthesis contributes to virulence of *Pseudomonas syringae* strain DC3000. PLoS Pathog. 14, e1006811. DOI: 10.1371/journal.ppat.1006811.

42. Medema, M. H., Takano, E. and Breitling, R. (2013) Detecting sequence homology at the gene cluster level with MultiGeneBlast. Molecular Biology and Evolution 30, 1218– 1223. DOI: 10.1093/molbev/mst025.

43. Raaijmakers, J. M., de Bruijn, I., Nybroe, O. and Ongena, M. (2010) Natural functions of lipopeptides from *Bacillus* and *Pseudomonas*: more than surfactants and antibiotics. FEMS Microbiol Rev 34, 1037–1062. DOI: 10.1111/j.1574-6976.2010.00221.x.

44. Cimermancic, P., Medema, M. H., Claesen, J., Kurita, K., Wieland Brown, L. C., Mavrommatis, K., Pati, A., Godfrey, P. A., Koehrsen, M., Clardy, J., Birren, B. W., Takano, E., Sali, A., Linington, R. G. and Fischbach, M. A. (2014) Insights into secondary metabolism from a global analysis of prokaryotic biosynthetic gene clusters. Cell 158, 412–421. DOI: 10.1016/j.cell.2014.06.034.

45. Cézard, C., Farvacques, N. and Sonnet, P. (2015) Chemistry and biology of pyoverdines, *Pseudomonas* primary siderophores. Curr. Med. Chem. 22, 165–186.

46. Berti, A. D. and Thomas, M. G. (2009) Analysis of achromobactin biosynthesis by *Pseudomonas syringae* pv. *syringae* B728a. Journal of Bacteriology 191, 4594–4604. DOI: 10.1128/JB.00457-09.

47. Matthijs, S., Budzikiewicz, H., Schäfer, M., Wathelet, B. and Cornelis, P. (2008) Ornicorrugatin, a New Siderophore from *Pseudomonas fluorescens* AF76. Zeitschrift für Naturforschung C 63, 8–12. DOI: 10.1515/znc-2008-1-202.

48. Patel, H. M. and Walsh, C. T. (2001) In Vitro Reconstitution of the *Pseudomonas aeruginosa* Nonribosomal Peptide Synthesis of Pyochelin: Characterization of Backbone Tailoring Thiazoline Reductase and N-Methyltransferase Activities. Biochemistry 40, 9023–9031. DOI: 10.1021/bi010519n.

49. Mercado-Blanco, J., van der Drift, K. M., Olsson, P. E., Thomas-Oates, J. E., van Loon, L. C. and Bakker, P. A. (2001) Analysis of the pmsCEAB gene cluster involved in biosynthesis of salicylic acid and the siderophore pseudomonine in the biocontrol strain *Pseudomonas fluorescens* WCS374. Journal of Bacteriology 183, 1909–1920. DOI: 10.1128/JB.183.6.1909-1920.2001.

50. Kim, S. Y., Ju, K.-S., Metcalf, W. W., Evans, B. S., Kuzuyama, T. and van der Donk, W. A. (2012) Different biosynthetic pathways to fosfomycin in *Pseudomonas syringae* and *Streptomyces* species. Antimicrob. Agents Chemother. 56, 4175–4183. DOI: 10.1128/AAC.06478-11.

51. Repka, L. M., Chekan, J. R., Nair, S. K. and van der Donk, W. A. (2017) Mechanistic Understanding of Lanthipeptide Biosynthetic Enzymes. Chem. Rev. 117, 5457–5520. DOI: 10.1021/acs.chemrev.6b00591.

52. Velasco, A., Acebo, P., Gomez, A., Schleissner, C., Rodríguez, P., Aparicio, T., Conde, S., Muñoz, R., la Calle, de, F., Garcia, J. L. and Sánchez-Puelles, J. M. (2005) Molecular characterization of the safracin biosynthetic pathway from *Pseudomonas fluorescens* A2-2: designing new cytotoxic compounds. Mol. Microbiol. 56, 144–154. DOI: 10.1111/j.1365-2958.2004.04433.x.

53. Coulthurst, S. J., Barnard, A. M. L. and Salmond, G. P. C. (2005) Regulation and biosynthesis of carbapenem antibiotics in bacteria. Nat. Rev. Micro. 3, 295–306. DOI: 10.1038/nrmicro1128.

54. Kudo, F. and Eguchi, T. (2009) Biosynthetic genes for aminoglycoside antibiotics. The Journal of Antibiotics 62, 471–481. DOI: 10.1038/ja.2009.76.

55. Leveau, J. H. J. and Gerards, S. (2008) Discovery of a bacterial gene cluster for catabolism of the plant hormone indole 3-acetic acid. FEMS Microbiol. Ecol. 65, 238– 250. DOI: 10.1111/j.1574-6941.2008.00436.x.

56. Puehringer, S., Metlitzky, M. and Schwarzenbacher, R. (2008) The pyrroloquinoline quinone biosynthesis pathway revisited: a structural approach. BMC Biochem. 9, 8. DOI: 10.1186/1471-2091-9-8.

57. Choi, O., Kim, J., Kim, J.-G., Jeong, Y., Moon, J. S., Park, C. S. and Hwang, I. (2008) Pyrroloquinoline quinone is a plant growth promotion factor produced by *Pseudomonas fluorescens* B16. Plant Physiol. 146, 657–668. DOI: 10.1104/pp.107.112748.

58. Zachow, C., Jahanshah, G., de Bruijn, I., Song, C., Ianni, F., Pataj, Z., Gerhardt, H., Pianet, I., Lämmerhofer, M., Berg, G., Gross, H. and Raaijmakers, J. M. (2015) The Novel Lipopeptide Poaeamide of the Endophyte *Pseudomonas poae* RE*1-1-14 Is Involved in Pathogen Suppression and Root Colonization. Mol. Plant Microbe Interact. 28, 800–810. DOI: 10.1094/MPMI-12-14-0406-R.

59. Hunziker, L., Bönisch, D., Groenhagen, U., Bailly, A., Schulz, S. and Weisskopf, L. (2015) *Pseudomonas* strains naturally associated with potato plants produce volatiles with high potential for inhibition of *Phytophthora infestans*. Applied and Environmental Microbiology 81, 821–830. DOI: 10.1128/AEM.02999-14.

60. Hultberg, M., Bengtsson, T. and Liljeroth, E. (2010) Late blight on potato is suppressed by the biosurfactant-producing strain *Pseudomonas koreensis* 2.74 and its biosurfactant. BioControl 55, 543–550. DOI: 10.1007/s10526-010-9289-7.

61. Jang, J. Y., Yang, S. Y., Kim, Y. C., Lee, C. W., Park, M. S., Kim, J. C. and Kim, I. S. (2013) Identification of orfamide A as an insecticidal metabolite produced by *Pseudomonas protegens* F6. J. Agric. Food Chem. 61, 6786–6791. DOI: 10.1021/jf401218w.

62. Alsohim, A. S., Taylor, T. B., Barrett, G. A., Gallie, J., Zhang, X.-X., Altamirano-Junqueira, A. E., Johnson, L. J., Rainey, P. B. and Jackson, R. W. (2014) The biosurfactant viscosin produced by *Pseudomonas fluorescens* SBW25 aids spreading motility and plant growth promotion. Environmental Microbiology 16, 2267–2281. DOI: 10.1111/1462-2920.12469.

63. Straight, P. D., Willey, J. M. and Kolter, R. (2006) Interactions between *Streptomyces coelicolor* and *Bacillus subtilis*: Role of surfactants in raising aerial structures. Journal of Bacteriology 188, 4918–4925. DOI: 10.1128/JB.00162-06.

64. Han, J. S., Cheng, J. H., Yoon, T. M., Song, J., Rajkarnikar, A., Kim, W. G., Yoo, I. D., Yang, Y. Y. and Suh, J. W. (2005) Biological control agent of common scab disease by antagonistic strain *Bacillus* sp. sunhua. J. Appl. Microbiol. 99, 213–221. DOI: 10.1111/j.1365-2672.2005.02614.x.

65. de Bruijn, I., de Kock, M. J. D., Yang, M., de Waard, P., van Beek, T. A. and Raaijmakers, J. M. (2007) Genome-based discovery, structure prediction and functional analysis of cyclic lipopeptide antibiotics in *Pseudomonas* species. Mol. Microbiol. 63, 417–428. DOI: 10.1111/j.1365-2958.2006.05525.x.

66. Nielsen, T. H., Thrane, C., Christophersen, C., Anthoni, U. and Sørensen, J. (2000) Structure, production characteristics and fungal antagonism of tensin - a new antifungal cyclic lipopeptide from *Pseudomonas fluorescens* strain 96.578. J. Appl. Microbiol. 89, 992–1001.

67. Götze, S., Herbst-Irmer, R., Klapper, M., Görls, H., Schneider, K. R. A., Barnett, R., Burks, T., Neu, U. and Stallforth, P. (2017) Structure, Biosynthesis, and Biological Activity of the Cyclic Lipopeptide Anikasin. ACS Chem. Biol. 12, 2498–2502. DOI: 10.1021/acschembio.7b00589.

68. Kuiper, I., Lagendijk, E. L., Pickford, R., Derrick, J. P., Lamers, G. E. M., Thomas-Oates, J. E., Lugtenberg, B. J. J. and Bloemberg, G. V. (2004) Characterization of two *Pseudomonas putida* lipopeptide biosurfactants, putisolvin I and II, which inhibit biofilm formation and break down existing biofilms. Mol. Microbiol. 51, 97–113. DOI: 10.1046/j.1365-2958.2003.03751.x.

69. Götze, S., Arp, J., Lackner, G., Zhang, S., Kries, H., Klapper, M., García-Altares, M., Willing, K., Günther, M. and Stallforth, P. (2019) Structure elucidation of the syringafactin lipopeptides provides insight in the evolution of nonribosomal peptide synthetases. Chem. Sci. 10, 10979–10990. DOI: 10.1039/C9SC03633D.

70. Pauwelyn, E., Huang, C.-J., Ongena, M., Leclère, V., Jacques, P., Bleyaert, P., Budzikiewicz, H., Schäfer, M. and Höfte, M. (2013) New linear lipopeptides produced by *Pseudomonas cichorii* SF1-54 are involved in virulence, swarming motility, and biofilm formation. Mol. Plant Microbe Interact. 26, 585–598. DOI: 10.1094/MPMI-11-12-0258-R.

71. Wang, M., Carver, J. J., Phelan, V. V., Sanchez, L. M., Garg, N., Peng, Y., Nguyen, D. D., Watrous, J., Kapono, C. A., Luzzatto-Knaan, T., Porto, C., Bouslimani, A., Melnik, A. V., Meehan, M. J., Liu, W.-T., Crüsemann, M., Boudreau, P. D., Esquenazi, E., Sandoval-Calderón, M., Kersten, R. D., Pace, L. A., Quinn, R. A., Duncan, K. R., Hsu, C.-C., Floros, D. J., Gavilan, R. G., Kleigrewe, K., Northen, T., Dutton, R. J., Parrot, D., Carlson, E. E., Aigle, B., Michelsen, C. F., Jelsbak, L., Sohlenkamp, C., Pevzner, P., Edlund, A., McLean, J., Piel, J., Murphy, B. T., Gerwick, L., Liaw, C.-C., Yang, Y.-L., Humpf, H.-U., Maansson, M., Keyzers, R. A., Sims, A. C., Johnson, A. R., Sidebottom, A. M., Sedio, B. E., Klitgaard, A., Larson, C. B., P, C. A. B., Torres-Mendoza, D., Gonzalez, D. J., Silva, D. B., Marques, L. M., Demarque, D. P., Pociute, E., O’Neill, E. C., Briand, E., Helfrich, E. J. N., Granatosky, E. A., Glukhov, E., Ryffel, F., Houson, H., Mohimani, H., Kharbush, J. J., Zeng, Y., Vorholt, J. A., Kurita, K. L., Charusanti, P., McPhail, K. L., Nielsen, K. F., Vuong, L., Elfeki, M., Traxler, M. F., Engene, N., Koyama, N., Vining, O. B., Baric, R., Silva, R. R., Mascuch, S. J., Tomasi, S., Jenkins, S., Macherla, V., Hoffman, T., Agarwal, V., Williams, P. G., Dai, J., Neupane, R., Gurr, J., Rodríguez, A. M. C., Lamsa, A., Zhang, C., Dorrestein, K., Duggan, B. M., Almaliti, J., Allard, P.-M., Phapale, P., Nothias, L.-F., Alexandrov, T., Litaudon, M., Wolfender, J.-L., Kyle, J. E., Metz, T. O., Peryea, T., Nguyen, D.-T., VanLeer, D., Shinn, P., Jadhav, A., Müller, R., Waters, K. M., Shi, W., Liu, X., Zhang, L., Knight, R., Jensen, P. R., Palsson, B. O., Pogliano, K., Linington, R. G., Gutiérrez, M., Lopes, N. P., Gerwick, W. H., Moore, B. S., Dorrestein, P. C. and Bandeira, N. (2016) Sharing and community curation of mass spectrometry data with Global Natural Products Social Molecular Networking. Nat. Biotechnol. 34, 828–837. DOI: 10.1038/nbt.3597.

72. Aron, A. T., Gentry, E. C., McPhail, K. L., Nothias, L.-F., Nothias-Esposito, M., Bouslimani, A., Petras, D., Gauglitz, J. M., Sikora, N., Vargas, F., van der Hooft, J. J. J., Ernst, M., Kang, K. B., Aceves, C. M., Caraballo-Rodríguez, A. M., Koester, I., Weldon, K. C., Bertrand, S., Roullier, C., Sun, K., Tehan, R. M., Boya P, C. A., Christian, M. H., Gutiérrez, M., Ulloa, A. M., Tejeda Mora, J. A., Mojica-Flores, R., Lakey-Beitia, J., Vásquez-Chaves, V., Zhang, Y., Calderón, A. I., Tayler, N., Keyzers, R. A., Tugizimana, F., Ndlovu, N., Aksenov, A. A., Jarmusch, A. K., Schmid, R., Truman, A. W., Bandeira, N., Wang, M. and Dorrestein, P. C. (2020) Reproducible molecular networking of untargeted mass spectrometry data using GNPS. Nat Protoc 15, 1954–1991. DOI: 10.1038/s41596-020-0317-5.

73. Flardh, K. and Buttner, M. J. (2009) *Streptomyces* morphogenetics: dissecting differentiation in a filamentous bacterium. Nat Rev Micro 7, 36–49. DOI: 10.1038/nrmicro1968.

74. Choi, K.-H., Gaynor, J. B., White, K. G., Lopez, C., Bosio, C. M., Karkhoff-Schweizer, R. R. and Schweizer, H. P. (2005) A Tn7-based broad-range bacterial cloning and expression system. Nat. Meth. 2, 443–448. DOI: 10.1038/nmeth765.

75. Rokni-Zadeh, H., Li, W., Sanchez-Rodriguez, A., Sinnaeve, D., Rozenski, J., Martins, J. C. and De Mot, R. (2012) Genetic and functional characterization of cyclic lipopeptide white-line-inducing principle (WLIP) production by rice rhizosphere isolate *Pseudomonas putida* RW10S2. Applied and Environmental Microbiology 78, 4826– 4834. DOI: 10.1128/AEM.00335-12.

76. Siddiqui, I. A., Shaukat, S. S., Sheikh, I. H. and Khan, A. (2006) Role of cyanide production by *Pseudomonas fluorescens* CHA0 in the suppression of root-knot nematode, *Meloidogyne javanica* in tomato. World J. Microbiol. Biotechnol. 22, 641–650. DOI: 10.1007/s11274-005-9084-2.

77. Ugidos, A., Morales, G., Rial, E., Williams, H. D. and Rojo, F. (2008) The coordinate regulation of multiple terminal oxidases by the *Pseudomonas putida* ANR global regulator. Environmental Microbiology 10, 1690–1702. DOI: 10.1111/j.1462-2920.2008.01586.x.

78. Comolli, J. C. and Donohue, T. J. (2002) *Pseudomonas aeruginosa* RoxR, a response regulator related to *Rhodobacter sphaeroides* PrrA, activates expression of the cyanide-insensitive terminal oxidase. Mol. Microbiol. 45, 755–768. DOI: 10.1046/j.1365-2958.2002.03046.x.

79. Feigl, F. and Anger, V. (1966) Replacement of benzidine by copper ethylacetoacetate and tetra base as spot-test reagent for hydrogen cyanide and cyanogen. Analyst 91, 282–284. DOI: 10.1039/AN9669100282.

80. Tschowri, N., Schumacher, M. A., Schlimpert, S., Chinnam, N. B., Findlay, K. C., Brennan, R. G. and Buttner, M. J. (2014) Tetrameric c-di-GMP mediates effective transcription factor dimerization to control *Streptomyces* development. Cell 158, 1136– 1147. DOI: 10.1016/j.cell.2014.07.022.

81. Andrade, M. H. M. L., Niederheitmann, M., de Paula Ribeiro, S. R. R., Oliveira, L. C., Pozza, E. A. and Pinto, C. A. B. P. (2019) Development and validation of a standard area diagram to assess common scab in potato tubers. Eur J Plant Pathol 154, 739–750. DOI: 10.1007/s10658-019-01697-z.

82. Nowicki, M., Foolad, M. R., Nowakowska, M. and Kozik, E. U. (2011) Potato and Tomato Late Blight Caused by *Phytophthora infestans*: An Overview of Pathology and Resistance Breeding. Plant Dis 96, 4–17. DOI: 10.1094/PDIS-05-11-0458.

83. Folders, J., Algra, J., Roelofs, M. S., van Loon, L. C., Tommassen, J. and Bitter, W. (2001) Characterization of *Pseudomonas aeruginosa* chitinase, a gradually secreted protein. Journal of Bacteriology 183, 7044–7052. DOI: 10.1128/JB.183.24.7044-7052.2001.

84. Laarman, A. J., Bardoel, B. W., Ruyken, M., Fernie, J., Milder, F. J., van Strijp, J. A. G. and Rooijakkers, S. H. M. (2012) *Pseudomonas aeruginosa* alkaline protease blocks complement activation via the classical and lectin pathways. J. Immunol. 188, 386–393. DOI: 10.4049/jimmunol.1102162.

85. Kenney, G. E., Dassama, L. M. K., Pandelia, M.-E., Gizzi, A. S., Martinie, R. J., Gao, P., DeHart, C. J., Schachner, L. F., Skinner, O. S., Ro, S. Y., Zhu, X., Sadek, M., Thomas, P. M., Almo, S. C., Bollinger, J. M., Krebs, C., Kelleher, N. L. and Rosenzweig, A. C. (2018) The biosynthesis of methanobactin. Science 359, 1411–1416. DOI: 10.1126/science.aap9437.

86. Ting, C. P., Funk, M. A., Halaby, S. L., Zhang, Z., Gonen, T. and van der Donk, W. A. (2019) Use of a scaffold peptide in the biosynthesis of amino acid-derived natural products. Science 365, 280–284. DOI: 10.1126/science.aau6232.

87. Clark, I. C., Melnyk, R. A., Iavarone, A. T., Novichkov, P. S. and Coates, J. D. (2014) Chlorate reduction in *Shewanella algae* ACDC is a recently acquired metabolism characterized by gene loss, suboptimal regulation and oxidative stress. Mol. Microbiol. 94, 107–125. DOI: 10.1111/mmi.12746.

88. Sarkisova, S. A., Lotlikar, S. R., Guragain, M., Kubat, R., Cloud, J., Franklin, M. J. and Patrauchan, M. A. (2014) A *Pseudomonas aeruginosa* EF-hand protein, EfhP (PA4107), modulates stress responses and virulence at high calcium concentration. PLoS ONE 9, e98985. DOI: 10.1371/journal.pone.0098985.

89. Price, M. N., Wetmore, K. M., Waters, R. J., Callaghan, M., Ray, J., Liu, H., Kuehl, J. V., Melnyk, R. A., Lamson, J. S., Suh, Y., Carlson, H. K., Esquivel, Z., Sadeeshkumar, H., Chakraborty, R., Zane, G. M., Rubin, B. E., Wall, J. D., Visel, A., Bristow, J., Blow, M. J., Arkin, A. P. and Deutschbauer, A. M. (2018) Mutant phenotypes for thousands of bacterial genes of unknown function. Nature 557, 503–509. DOI: 10.1038/s41586-018-0124-0.

90. Mavrodi, O. V., Mavrodi, D. V., Parejko, J. A., Thomashow, L. S. and Weller, D. M. (2012) Irrigation differentially impacts populations of indigenous antibiotic-producing *Pseudomonas* spp. in the rhizosphere of wheat. Applied and Environmental Microbiology 78, 3214–3220. DOI: 10.1128/AEM.07968-11.

91. Mavrodi, D. V., Mavrodi, O. V., Elbourne, L. D. H., Tetu, S., Bonsall, R. F., Parejko, J., Yang, M., Paulsen, I. T., Weller, D. M. and Thomashow, L. S. (2018) Long-Term Irrigation Affects the Dynamics and Activity of the Wheat Rhizosphere Microbiome. Front Plant Sci 9, 345. DOI: 10.3389/fpls.2018.00345.

92. Beskrovnaya, P., Melnyk, R. A., Liu, Z., Liu, Y., Higgins, M. A., Song, Y., Ryan, K. S. and Haney, C. H. (2020) Comparative Genomics Identified a Genetic Locus in Plant-Associated *Pseudomonas* spp. That Is Necessary for Induced Systemic Susceptibility. MBio 11, 356. DOI: 10.1128/mBio.00575-20.

93. Mullins, A. J., Murray, J. A. H., Bull, M. J., Jenner, M., Jones, C., Webster, G., Green, A. E., Neill, D. R., Connor, T. R., Parkhill, J., Challis, G. L. and Mahenthiralingam, E. (2019) Genome mining identifies cepacin as a plant-protective metabolite of the biopesticidal bacterium *Burkholderia ambifaria*. Nature Microbiology 4, 996–1005. DOI: 10.1038/s41564-019-0383-z.

94. Carrión, V. J., Perez-Jaramillo, J., Cordovez, V., Tracanna, V., de Hollander, M., Ruiz-Buck, D., Mendes, L. W., van Ijcken, W. F. J., Gómez Expósito, R., Elsayed, S. S., Mohanraju, P., Arifah, A., van der Oost, J., Paulson, J. N., Mendes, R., van Wezel, G. P., Medema, M. H. and Raaijmakers, J. M. (2019) Pathogen-induced activation of disease-suppressive functions in the endophytic root microbiome. Science 366, 606– 612. DOI: 10.1126/science.aaw9285.

95. Tracanna, V., Ossowicki, A., Petrus, M. L. C., Overduin, S., Terlouw, B. R., Lund, G., Robinson, S. L., Warris, S., Schijlen, E. G. W. M., van Wezel, G. P., Raaijmakers, J. M., Garbeva, P. and Medema, M. H. (2021) Dissecting Disease-Suppressive Rhizosphere Microbiomes by Functional Amplicon Sequencing and 10× Metagenomics. mSystems e0111620. DOI: 10.1128/mSystems.01116-20.

96. Rijavec, T. and Lapanje, A. (2016) Hydrogen Cyanide in the Rhizosphere: Not Suppressing Plant Pathogens, but Rather Regulating Availability of Phosphate. Front Microbiol 7, 1785. DOI: 10.3389/fmicb.2016.01785.

97. Kodani, S., Hudson, M. E., Durrant, M. C., Buttner, M. J., Nodwell, J. R. and Willey, J. M. (2004) The SapB morphogen is a lantibiotic-like peptide derived from the product of the developmental gene ramS in *Streptomyces coelicolor*. Proc. Natl. Acad. Sci. U.S.A. 101, 11448–11453. DOI: 10.1073/pnas.0404220101.

98. Willey, J. M., Willems, A., Kodani, S. and Nodwell, J. R. (2006) Morphogenetic surfactants and their role in the formation of aerial hyphae in *Streptomyces coelicolor*. Mol. Microbiol. 59, 731–742. DOI: 10.1111/j.1365-2958.2005.05018.x.

99. Fukuda, T. T. H., Pereira, C. F., Melo, W. G. P., Menegatti, C., Andrade, P. H. M., Groppo, M., Lacava, P. T., Currie, C. R. and Pupo, M. T. (2021) Insights Into the Ecological Role of *Pseudomonas* spp. in an Ant-plant Symbiosis. Front Microbiol 12, 621274. DOI: 10.3389/fmicb.2021.621274.

100. Helfrich, E. J. N., Vogel, C. M., Ueoka, R., Schäfer, M., Ryffel, F., Müller, D. B., Probst, S., Kreuzer, M., Piel, J. and Vorholt, J. A. (2018) Bipartite interactions, antibiotic production and biosynthetic potential of the *Arabidopsis* leaf microbiome. Nature Microbiology 3, 909–919. DOI: 10.1038/s41564-018-0200-0.

101. Bakker, P. A. H. M., Pieterse, C. M. J. and van Loon, L. C. (2007) Induced Systemic Resistance by Fluorescent *Pseudomonas* spp. Phytopathology 97, 239–243. DOI: 10.1094/PHYTO-97-2-0239.

102. Teixeira, P. J. P. L., Colaianni, N. R., Law, T. F., Conway, J. M., Gilbert, S., Li, H., Salas-González, I., Panda, D., Del Risco, N. M., Finkel, O. M., Castrillo, G., Mieczkowski, P., Jones, C. D. and Dangl, J. L. (2021) Specific modulation of the root immune system by a community of commensal bacteria. Proc. Natl. Acad. Sci. U.S.A. 118. DOI: 10.1073/pnas.2100678118.

103. Xu, L., Naylor, D., Dong, Z., Simmons, T., Pierroz, G., Hixson, K. K., Kim, Y.-M., Zink, E. M., Engbrecht, K. M., Wang, Y., Gao, C., DeGraaf, S., Madera, M. A., Sievert, J. A., Hollingsworth, J., Birdseye, D., Scheller, H. V., Hutmacher, R., Dahlberg, J., Jansson, C., Taylor, J. W., Lemaux, P. G. and Coleman-Derr, D. (2018) Drought delays development of the sorghum root microbiome and enriches for monoderm bacteria. Proc. Natl. Acad. Sci. U.S.A. 115, E4284–E4293. DOI: 10.1073/pnas.1717308115.

104. Santos-Medellín, C., Liechty, Z., Edwards, J., Nguyen, B., Huang, B., Weimer, B. C. and Sundaresan, V. (2021) Prolonged drought imparts lasting compositional changes to the rice root microbiome. Nat. Plants 7, 1065–1077. DOI: 10.1038/s41477-021-00967-1.

105. van der Meij, A., Worsley, S. F., Hutchings, M. I. and van Wezel, G. P. (2017) Chemical ecology of antibiotic production by actinomycetes. FEMS Microbiol Rev 41, 392–416. DOI: 10.1093/femsre/fux005.

106. Miller, J. H. (1972) Experiments in Molecular Genetics. Cold Spring Harbor Laboratory, New York.

107. Kieser, T., Bibb, M. J., Buttner, M. J., Chater, K. F. and Hopwood, D. A. (2000) Practical Streptomyces Genetics. John Innes Foundation, Norwich.

108. King, E. O., Ward, M. K. and Raney, D. E. (1954) Two simple media for the demonstration of pyocyanin and fluorescin. J. Lab. Clin. Med. 44, 301–307.

109. Caporaso, J. G., Lauber, C. L., Walters, W. A., Berg-Lyons, D., Lozupone, C. A., Turnbaugh, P. J., Fierer, N. and Knight, R. (2011) Global patterns of 16S rRNA diversity at a depth of millions of sequences per sample. Proc. Natl. Acad. Sci. U.S.A. 108, 4516– 4522. https://doi.org/10.1073/pnas.1000080107.

110. Castric, K. F. and Castric, P. A. (1983) Method for rapid detection of cyanogenic bacteria. Applied and Environmental Microbiology 45, 701–702.

111. Caten, C. E. and Jinks, J. L. (1968) Spontaneous variability of single isolates of *Phytophthora infestans*. I. Cultural variation. Canadian Journal of Botany 46, 329–348. DOI: 10.1139/b68-055.

112. Zimin, A. V., Marçais, G., Puiu, D., Roberts, M., Salzberg, S. L. and Yorke, J. A. (2013) The MaSuRCA genome assembler. Bioinformatics 29, 2669–2677. DOI: 10.1093/bioinformatics/btt476.

113. Bankevich, A., Nurk, S., Antipov, D., Gurevich, A. A., Dvorkin, M., Kulikov, A. S., Lesin, V. M., Nikolenko, S. I., Pham, S., Prjibelski, A. D., Pyshkin, A. V., Sirotkin, A. V., Vyahhi, N., Tesler, G., Alekseyev, M. A. and Pevzner, P. A. (2012) SPAdes: a new genome assembly algorithm and its applications to single-cell sequencing. J. Comput. Biol. 19, 455–477. DOI: 10.1089/cmb.2012.0021.

114. Seemann, T. (2014) Prokka: rapid prokaryotic genome annotation. Bioinformatics 30, 2068–2069. DOI: 10.1093/bioinformatics/btu153.

115. Hyatt, D., Chen, G.-L., Locascio, P. F., Land, M. L., Larimer, F. W. and Hauser, L. J. (2010) Prodigal: prokaryotic gene recognition and translation initiation site identification. BMC Bioinformatics 11, 119. DOI: 10.1186/1471-2105-11-119.

116. Parks, D. H., Imelfort, M., Skennerton, C. T., Hugenholtz, P. and Tyson, G. W. (2015) CheckM: assessing the quality of microbial genomes recovered from isolates, single cells, and metagenomes. Genome Res. 25, 1043–1055. DOI: 10.1101/gr.186072.114.

117. Edgar, R. C. (2004) MUSCLE: multiple sequence alignment with high accuracy and high throughput. Nucleic Acids Research 32, 1792–1797. DOI: 10.1093/nar/gkh340.

118. Stamatakis, A. (2014) RAxML version 8: a tool for phylogenetic analysis and post-analysis of large phylogenies. Bioinformatics 30, 1312–1313. DOI: 10.1093/bioinformatics/btu033.

119. Miller, M. A., Schwartz, T., Pickett, B. E., He, S., Klem, E. B., Scheuermann, R. H., Passarotti, M., Kaufman, S. and O’Leary, M. A. (2015) A RESTful API for Access to Phylogenetic Tools via the CIPRES Science Gateway. Evol. Bioinform. Online 11, 43– 48. DOI: 10.4137/EBO.S21501.

120. Letunic, I. and Bork, P. (2021) Interactive Tree Of Life (iTOL) v5: an online tool for phylogenetic tree display and annotation. Nucleic Acids Res. 49, W293–W296. https://doi.org/10.1093/nar/gkab301.

121. Campilongo, R., Fung, R. K. Y., Little, R. H., Grenga, L., Trampari, E., Pepe, S., Chandra, G., Stevenson, C. E. M., Roncarati, D. and Malone, J. G. (2017) One ligand, two regulators and three binding sites: How KDPG controls primary carbon metabolism in *Pseudomonas*. PLoS Genet 13, e1006839. DOI: 10.1371/journal.pgen.1006839.

122. Scott, T. A., Heine, D., Qin, Z. and Wilkinson, B. (2017) An L-threonine transaldolase is required for L-threo-β-hydroxy-α-amino acid assembly during obafluorin biosynthesis. Nat. Commun. 8, 15935. DOI: 10.1038/ncomms15935.

123. Shannon, P., Markiel, A., Ozier, O., Baliga, N. S., Wang, J. T., Ramage, D., Amin, N., Schwikowski, B. and Ideker, T. (2003) Cytoscape: a software environment for integrated models of biomolecular interaction networks. Genome Res. 13, 2498–2504. https://doi.org/10.1101/gr.1239303.

124. Sarwar, A., Latif, Z., Zhang, S., Zhu, J., Zechel, D. L. and Bechthold, A. (2018) Biological Control of Potato Common Scab With Rare Isatropolone C Compound Produced by Plant Growth Promoting *Streptomyces* A1RT. Front. Microbiol. 9, 41. DOI: 10.3389/fmicb.2018.01126.

125. Lin, C., Tsai, C.-H., Chen, P.-Y., Wu, C.-Y., Chang, Y.-L., Yang, Y.-L. and Chen, Y.-L. (2018) Biological control of potato common scab by *Bacillus amyloliquefaciens* Ba01. PLoS ONE 13, e0196520. DOI: 10.1371/journal.pone.0196520.

