## Supplementary Information for "Pan-genome analysis identifies intersecting roles for *Pseudomonas* specialized metabolites in potato pathogen inhibition"

### SUPPLEMENTARY METHODS

#### Identification and scoring of biosynthetic gene clusters and accessory genome loci

All genome sequences were subjected to biosynthetic gene cluster (BGC) analysis using antiSMASH 5.0 (1). Any "similar known cluster" and/or MIBiG BGC-ID annotations (2) from antiSMASH were assessed for their accuracy via a manual comparison with the putative matching BGC. This was particularly important for BGCs that are split across distinct genomic loci, such as pyoverdine and viscosin (3). When a characterised homologous BGC could not be identified, the BGCs were named and numbered based on their biosynthetic class (e.g. NRPS 1, NRPS 2, NRPS 3). To ensure that no BGCs had been missed by antiSMASH analysis, all genomes were searched against a library of known *Pseudomonas* BGCs using MultiGeneBlast (4) (settings: minimal sequence coverage of BLAST hits = 25%; minimal identity of BLAST hits = 25%; maximal distance between genes in locus = 20 kb). MultiGeneBlast was also used to validate the antiSMASH annotations. Brief details of each BGC are summarised below and were scored as 0 (no BGC), 1 (partial BGC) or 2 (full BGC). Scoring criteria vary between BGC type in relation to the requirements for a functional BGC in each case (details below). Where conserved domains were not evident in an antiSMASH output, their identity and specificity (for NRPS/PKS domains) were further assessed using NCBI CD-Search (5) and NRPSpredictor2 (6). When a single protein defined a genotype (such as secreted proteases), BLAST analysis (7) was carried out to identify homologues. Results are summarised in Supporting Dataset 1.

*Key to domain nomenclature for polyketide synthases (PKSs) and non-ribosomal peptide synthetases (NRPSs):* A = adenylation; T = thiolation; C = condensation; TE = thioesterase; AT = acyltransferase; KS = keto synthase; KR = ketoreductase; DH = dehydratase; ER = enoylreductase; MT = methyltransferase. / = denotes the boundary between PKS or NRPS proteins.

**NRPSs**

Note that some NRPS BGCs are described in the siderophore section, such as pyoverdine.

**Cyclic and linear lipopeptides:** These were subjected to detailed analysis as reported elsewhere in Materials and Methods.

**Safracin:** A well-characterized antitumor compound produced by an NRPS (8). The full ten gene BGC is present in the Ps903 genome.

**mgo/pvf operon:** A conserved cluster that includes a single module NRPS protein with A-T-reductase domain organization and predicted leucine specificity. Initially identified as being involved in mangotoxin biosynthesis, the *mgo* operon is distinct from the *mbo* operon, which is actually responsible for mangotoxin biosynthesis (9). The *mgo* operon is homologous to the *Pseudomonas entomophila* virulence factor (*pvf*) gene cluster, which is a regulator of virulence factors (10, 11). This indicates that the product of this BGC functions as a regulator of other BGCs in *Pseudomonas* species.

**NRPS 1:** Trimodular NRPS (A-T-C-A-T-C-A-T-TE) with Thr-Thr-? predicted specificity.

**NRPS 2:** Three co-transcribed proteins featuring six NRPS modules plus a single PKS lacking an AT (A-T-C-A-T-C-A-T / C-A-T-TE / A-T-C-A-T-KS-KR-T-TE; predicted specificity = Arg-Pro-Cys / ?-TE / Val-Pro-?-TE). Two TE domains indicate that this could produce two metabolites. Wider cluster includes an MbtH-like protein and a dioxygenase.

**NRPS 3:** Single protein containing a single NRPS module (A-T-TE) with no known amino acid specificity. Adjacent to a glutathione S-transferase, but this is conserved in a number of strains that do not encode an adjacent NRPS so is unlikely to feature as part of this pathway.

**NRPS 4:** One protein containing a single atypical NRPS module: T-A-AT with predicted phenylglycine or threonine specificity.

**NRPS 5:** One protein containing a single NRPS module: A-T-TE with predicted threonine specificity.

**NRPS 6:** One protein containing a single NRPS module: A-T-AT with predicted glutamate specificity. Adjacent to additional putative biosynthetic genes (including genes that encode isochorismitases, peptidases and glycosyltransferases), but these differ between strains.

**NRPS 7:** One protein containing a putative NRPS module: sulfotransferase-A-T-reductase with predicted phenylglycine specificity. Homologous gene cluster analysis in antiSMASH indicates a wider gene cluster: methyltransferase, dioxygenase, dehydrogenase, epimerase, oxygenase, transcriptional regulator, transporter, NRPS, dehydrogenase, kinase.

**NRPS 8:** Single module PKS: KS-T-TE. Encoded alongside a 4'-pantetheinephosphotransferase and an ABC transporter. These three genes are homologous to a contiguous set of genes (PSF113_3664 to PSF113_3666) within a larger lankacidin-like PKS BGC in *Pseudomonas fluorescens* F113.

**NRPS 9:** One protein containing a single NRPS module: A-T-reductase, with predicted isoleucine specificity.

**NRPS 10:** One protein containing a single NRPS module fused to a P450: A-T-TE-P450. The specificity of the A domain could not be predicted.

**NRPS-PKS:** BGC with an unknown product identified in Ps843. NRPS-PKS organization: A-T / KS-AT-KR-C / TE. This reflects the order of genes but may not reflect functional order of the synthetase. antiSMASH analysis indicates a larger set of related genes associated with this NRPS-PKS, with multiple homologous gene clusters found in *Pseudomonas aeriginosa* strains.

**PKSs**

**Aryl polyenes:** Widespread type II PKS gene clusters detected by antiSMASH, as defined by MIBiG entry BGC0000837 (APE Vf from *Aliivibrio fischeri* ES114) (12). Cluster defined as aryl polyene if set of PKS genes are found alongside homologues of other characteristic aryl polyene biosynthetic genes, including ammonia lyase and acyltransferase.

**PKS 1:** BGC with an unknown product identified in Ps664. Putative type I PKS organization: A / T / KS-AT-KR-T-KS-AT-KR-T / KS-AfsA domain / TE. The AfsA domain has homology to the AfsA family of proteins required for A-factor biosynthesis (13).

**PKS 2:** BGC with an unknown product identified in Ps691. Putative type I PKS organization: KS-AT-DH-ER-KR-T. This single PKS module is associated with polysaccharide biosynthesis proteins, which is conserved across homologous clusters identified from antiSMASH analysis.

**PKS 3:** BGC with an unknown product identified in multiple strains. Putative type III PKS gene cluster encoding the following conserved proteins: type III PKS (NCBI conserved domain cd00831), oxidoreductase, SnoaL-like protein, methyltransferase, methyltransferase.

**SIDEROPHORES**

**Pyoverdine:** Characteristic biosynthetic genes identified by antiSMASH. Analysis complicated as genes are usually distributed between distinct genomic loci (3).

**Achromobactin:** Siderophore produced by *Pseudomonas syringae* pv. syringae B728a (14). Gene cluster defined as achromobactin when homologues of all biosynthetic genes (Psyr2582-Psyr2589) are present in a single BGC.

**Pseudomonine:** Siderophore produced by *Pseudomonas fluorescens* WCS374 (15). Gene cluster defined as pseudomonine when homologues of all biosynthetic genes (as defined in MIBiG entry BGC0000410) are present in a single BGC.

**Ornicorrugatin-like:** Lipopeptide siderophore produced by *Pseudomonas fluorescens* AF76 (16), whose gene cluster is described in *P. fluorescens* SBW25 (17). Gene cluster defined as ornicorrugatin-like when homologues of all SBW25 genes are present in a single BGC.

**Pyochelin-like 1:** Gene cluster is similar to the gene cluster to pyochelin (18) but contains an extra NRPS module predicted to incorporate cysteine. NRPS domain organization: A / C-A-T-C-A-T / C-A-MT-T-TE; predicted specificity = dihydroxybenzoic acid-Cys-Cys-Cys. This would be the right organization for an ulbactin F-like molecule (19).

**Pyochelin-like 2:** Similar NRPS organization to “pyochelin-like 1” gene cluster, but different set of associated genes. NRPS domain organization: A / C-A-T-C-A / T / C-A-MT-T-TE; predicted specificity = dihydroxybenzoic acid-Cys-Cys-Cys.

**Pyochelin-like 3:** Canonical pyochelin gene cluster with identical NRPS module organization to the characterized PchDEF system (18): A / T-C-A-T / C-A-MT-T-E; predicted specificity = dihydroxybenzoic acid-Cys-Cys.

**Quinolobactin:** Siderophore produced by *Pseudomonas fluorescens* ATCC 17400 (20). Gene cluster defined as quinolobactin when homologues of all biosynthetic genes (as defined in MIBiG entry BGC0000925) are present in a single BGC.

**Putative siderophore 1 and 2:** Gene clusters encoding pathways predicted to biosynthesize siderophores (21). Two distinct gene clusters were identified, so are defined as “Putative siderophore 1” (encoding: diaminopimelate decarboxylase, PLP-dependent enzyme, dehydrogenase, IucA/IucC-like siderophore biosynthesis protein, major facilitator transporter, aminotransferase) and “Putative siderophore 2” (encoding: argininosuccinate lyase-like protein, IucA/IucC-like siderophore biosynthesis protein, major facilitator transporter, IucA/IucC-like siderophore biosynthesis protein, diaminopimelate decarboxylase). A few strains encoded IucA/iucC-like proteins, but no other siderophore biosynthesis proteins were encoded alongside these proteins, so were not annotated as siderophore BGCs.

### RIBOSOMALLY SYNTHESIZED AND POST-TRANSLATIONALLY MODIFIED PEPTIDES (RiPPs)

**Microcin B17-like:** BGCs with homology to the microcin B17 gene cluster from *E. coli* (22, 23), encoding the following proteins: McbA-like precursor peptide, McbB-like cyclodehydratase component

(TIGR04424), McbC-like flavin-dependent dehydrogenase, McbD-like YcaO domain protein, McbE-like immunity protein, McbF-like immunity protein. Unlike the microcin B17-like gene clusters identified in *Pseudomonas syringae* (24), no McbG homologues were encoded in any of the strains in this current study.

**Lanthipeptide:** Putative precursor peptide encoded alongside a lanthipeptide synthetase (LanM family, TIGR03897).

**YcaO cluster:** Putative BGC encoding YcaO and TfuA domain proteins, which are characteristic of RiPP biosynthesis (25). BGC organization: putative precursor peptide, YcaO domain protein, TfuA domain protein, unknown domain protein, ubiquitin-like domain protein, E1/ThiF-like domain protein, major facilitator superfamily transporter, methyltransferase.

**"Pep" BGCs:**
All "Pep" BGCs encode short peptides alongside DUF692 domain proteins, which have been shown to be essential for the biosynthesis of the RiPP methanobactin (26). Below are lists of proteins encoded in each putative BGC (Figure S9):

**Pep1:** Short peptide (DUF2282), DUF692 protein, DUF2063 protein, DoxX domain protein (pfam07681).
**Pep2:** Short peptide (DUF2282), DUF692 protein, DUF2063 protein, methyltransferase, cardiolipin synthase.
**Pep3:** Short peptide (COG3767), DUF692 protein, DUF2063 protein, DMT family transporter, LysR family transcriptional regulator.
**Pep4:** Short peptide (DUF2282), DoxX domain protein, hydrolase, short peptide (no conserved domain), DUF692 protein, DUF2063 protein, hydrolase.
**Pep5:** short peptide (no conserved domain), DUF692 protein, DUF2063 protein, DoxX domain protein.

#### TERPENES

**Carotenoid:** BGC containing homologues of all carotenoid genes defined in MIBiG entry BGC0000642 from *Enterobacteriaceae* bacterium DC413 (27).

**Unknown terpene 1:** Putative BGC identified in Ps655. Terpene synthase/cyclase (NCBI conserved domain cd00687) encoded alongside a polyprenyl synthetase (pfam00348).

**Unknown terpene 2:** Putative BGC identified in Ps706. Fused terpene synthase/P450, methyltransferase, isopentenyl diphosphate isomerase.

#### OTHERS

**HCN:** BGC containing homologues of *hcnABC*, which together encode the HCN synthase complex. The *hcn* operon from *Pseudomonas aeruginosa* PAO1 (PA2193 - PA2195, Figure S9) (28) was used with MultiGeneBlast.

**Dimethyl sulfide (DMS):** MegL (methionine gamma-lyase) and MddA (methyltransferase) convert methionine to DMS in *Pseudomonas deceptionensis* (29). A DMS BGC is defined when a strain encodes homologues of both MddA and MegL with over 60% identity. A score of 1 was defined when it only encoded a homologue of MddA with over 60% identity but not MegL.

**Tabtoxin-like:** *Pseudomonas* Beta-lactam whose BGC is defined by MIBiG entry BGC0000846 (30). Identified by antiSMASH and confirmed by MultiGeneBlast.

**Indole-3-acetic acid 1 (IAA 1):** The IAA BGC encodes homologues of IaaM (tryptophan 2-monooxygenase) and IaaH (indoleacetamide hydrolase) from *Pseudomonas savastanoi* (for the indole-3-acetamide pathway to IAA) (31). Homologues of these were identified using MultiGeneBlast.

**Indole-3-acetic acid 2 (IAA 2):** Defined as encoding a protein with high homology to IdaA (PSPTO_0092) from *Pseudomonas syringae* pv. DC3000 (32). All strains encoded proteins with >88% identity to AldA. MultiGeneBlast analysis showed that these genes are all present in exactly the same genetic context as in *P. syringae* pv. DC3000.

**Homoserine lactone 1:** Defined as encoding a protein homologous to an acyl-homoserine-lactone synthase (pfam00765) from Ps887 (identified by antiSMASH analysis).

**Homoserine lactone 2:** Defined as encoding a protein homologous to the acyl-homoserine-lactone synthase HdtS from *Pseudomonas fluorescens* F113 (33).

**A-factor-like:** Identified by antiSMASH as an A-factor-like BGC in Ps664. Clustered genes encode an AfsA-like protein (13), a hydrolase, a P450 and a major facilitator superfamily transporter. This lacks the reductase encoded in a classical A-factor gene cluster (13).

**Aminoglycoside:** Identified by antiSMASH as an aminoglycoside-like BGC in Ps639 (Figure S6). The BGC encodes the following enzymes that could assemble an aminoglycoside (34) (putative biosynthetic roles are indicated): a 2-deoxy-scyllo-inosose synthase (47% identity to paromomycin homologue, ParC), a phosphoribosyltransferase (33% identity to neomycin homologue, NeoL), a neamine aminotransferase (36% identity to neomycin homologue, NeoN), a 2'-N-acetylparomamine deacetylase (37% identity to neomycin homologue, NeoD), a L-glutamine:scyllo-inosose aminotransferase (56% identity to paromomycin homologue, ParS), a 6'-hydroxyparomomycin dehydrogenase (51% identity to paromomycin homologue, ParQ), a 2-deoxystreptamine N-acetyl-glucosaminyltransferase (44% identity to neomycin homologue, NeoM), and a 2OG-Fe(II) oxygenase that is not homologous to known aminoglycoside biosynthetic enzymes.

**Coronamic acid:** This unusual amino acid (1-amino-2-ethylcyclopropane carboxylic acid) forms part of the *Pseudomonas savastanoi* phytotoxin coronatine. BGC identified by antiSMASH in Ps834 contains homologues of all coronamic acid biosynthetic genes, as defined by Couch *et al.* (35) (MIBiG entry BGC0000328). The Ps834 BGC is not associated with polyketide or ligase genes required for full coronatine biosynthesis (36).

**Fosfomycin-like:** The fosfomycin BGC from *Pseudomonas syringae* PB-5123 (37) was used with

MultiGeneBlast to search for similar BGCs. A significant number of homologous proteins are clustered in two strains, including multiple proposed biosynthetic proteins: trans-homoaconitate synthase (Psf2 homologue), epoxidase (Psf4 homologue), 6-phosphogluconate dehydrogenase (Psf3 homologue), fumarylacetoacetate hydrolase, a hypothetical protein, phosphoenolpyruvate phosphomutase (Psf1 homologue) (Figure S6).

**N-acetylglutaminylglutamine amide (NAGGN):** Dipeptide BGC identified by antiSMASH. Representative BGC present in *P. aeruginosa* PAO1 (38) (genes PA3459 and PA3460).

**Beta-lactones:** Identified by antiSMASH and defined by the presence of an AMP-dependent synthetase/ligase and a 2-isopropylmalate synthase (39). Three distinct BGC types were identified, where characteristic examples are present in Ps619 (Beta-lactone 1), Ps639 (Beta-lactone 2) and Ps659 (Beta-lactone 3).

**Ectoine-like 1:** BGC identified by antiSMASH in Ps663, although this is different to the well-characterised *ectABC* cluster identified in *Pseudomonas stutzeri* (40). The putative Ps663 cluster encodes: an AMP-dependent ligase, a hypothetical protein, a dehydrogenase, an ectoine synthase and a major facilitator superfamily transporter.

**Ectoine-like 2:** BGC identified by antiSMASH in Ps664 and differs from “ectoine-like 1” BGC. Features similarities with characterized ectoine gene clusters (e.g. MIBiG entry BGC0000859), including homologues of the ectoine synthase and transaminase genes. The Ps664 BGC encodes: ectoine synthase, transaminase, a dioxygenase and a putative transporter.

### PYRROLOQUINOLINE QUINONE (PQQ)

To search for PQQ BGCs, the *Pseudomonas fluorescens* Pf0-1 PQQ BGC (41) was used with MultiGeneBlast. Specifically, proteins PqqF, PqqA, PqqB, PqqC, PqqD, PqqE, PqqM were used, which are encoded by contiguous genes in *P. fluorescens* Pf0-1. This identified a series a distinct variants of PQQ BGCs (Figure S7):

**PQQ1:** BGC encoding all PQQ proteins as defined above.
**PQQ2:** BGC encoding PqqF, PqqA, PqqB, PqqC, PqqD, PqqE, but not PqqM. A putative amidase is encoded alongside PqqF, which could potentially functionally replace PqqM.
**PQQ3:** BGC encoding PqqA, PqqB, PqqC, PqqD, PqqE, but not PqqF or PqqM. BGC is only found in strains that also encode a PQQ1-type BGC.
**PQQ4:** BGC encoding PqqA, PqqC, PqqD, PqqE, but not PqqB, PqqF or PqqM. BGC is only found in strains that also encode a PQQ1-type BGC.

### GENE CLUSTERS NOT FOUND IN THIS STRAIN COLLECTION

In addition to the specific BGCs described above that are found in one or more strains in this study, a number of *Pseudomonas* BGCs were not present in any strains, as described below:

**Bicyclomycin:** *Psuedomonas aeruginosa* SCV20265 BGC (42) was used in MultiGeneBlast.

**Cyclodipeptides:** tRNA-dependent cyclodipeptide synthases from *Pseudomonas protegens* (NCBI accession OKK65715.1) and *P. aeruginosa* (WP_003158562.1) were used in BLAST analyses.

**Pyreudiones:** No strains encode the standalone C-A-T-TE NRPS that defines the *Pseudomonas fluorescens* HKI0770 BGC (43).

**Kalimantacin:** No strains encode the large hybrid PKS-NRPS that makes this molecule (44).

**Brabantamide:** BraA to BraE proteins from *Pseudomonas* sp. SH-C52 (45) was used with MultiGeneBlast.

**Pseudopyronines:** The characterized PpyS protein from *Pseudomonas putida* BW11M1 (46) was used in BLAST analyses.

**2,4-diacylphloroglucinol (DAPG):** The biosynthetic genes (*phlABCD*) from *Pseudomonas fluorescens* F113 (47) was used with MultiGeneBlast.

**Phenazines:** The BGC from *Pseudomonas fluorescens* 2-79 (48) (NCBI accession L48616.1) was used with MultiGeneBlast.

**Pyrrolnitrin:** The BGC (*prnABCD*) from *Pseudomonas aurantiaca* (previously named *P. fluorescens*) BL915 (49) was used with MultiGeneBlast.

**2,5-Dialkylresorcinols:** The *darABC* genes from *Pseudomonas chlororaphis* PCL1606 (50) (NCBI accession JQ663992.1) were used with MultiGeneBlast.

**Toxoflavin:** The toxoflavin BGC from *Pseudomonas protegens* Pf-5 (PFL_1028 to PFL_1037) (51) was used with MultiGeneBlast.

**L-2-Amino-4-methoxy-trans-3-butenoic acid (AMB):** The *ambABCDE* gene cluster from *P. aeruginosa* PAO1 (PA2302-PA2306) (52) was used with MultiGeneBlast.

### ACCESSORY GENOME LOCI

#### EXOPOLYSACCHARIDES

**Psl operon:** The *psl* gene cluster (PA2231 to PA2245) from *P. aeruginosa* PAO1 (53) was used with MultiGeneBlast. Score = 2 for all genes present; score = 1 for missing 1-3 genes in the cluster.

**Wss operon:** The *wss* gene cluster (*wssA-J*) from *P. fluorescens* SBW25 (54) was used with MultiGeneBlast. Score = 2 for all genes present; score = 1 for only homologues of *wssA-E* clustered.

**Pel operon:** The *pel* gene cluster (*pelA-G*) from *P. aeruginosa* PA14 (55) was used with MultiGeneBlast. Score = 2 for all genes present.

**Pga operon:** The *pga* gene cluster (*pgaA-D*) from *P. fluorescens* SBW25 (PFLU0143 -PFLU0146) (56) was used with MultiGeneBlast. Score = 2 for all genes present.

**Alginate:** The alginate gene cluster from *P. fluorescens* SBW25 (PFLU0979 - PFLU0990) (57) was used with MultiGeneBlast. Score = 2 for all genes present.

**STRESS RESPONSE POLYSACCHARIDES**

**Alpha glucan biosynthesis:** Biosynthetic proteins from *P. syringae* pv. DC3000 (PSPTO_2760-62, PSPTO_3125-30, PSPTO_5165) (58) were used with BLAST and MultiGeneBlast. Score = 2 for homologues of all genes present; score = 1 when 1-2 genes were absent from the PSPTO_3125-30 cluster.

**Trehalose degradation:** PSPTO_2952 protein from *P. syringae* pv. DC3000 (NCBI accession NP_792749.1) was used with BLAST. Score = 2 for homologues >70% identity.

**LIPOPOLYSACCHARIDES**

**Fuzzy spreader:** The *fuzVWXYZ* operon from *P. fluorescens* SBW25 (PFLU0475 - PFLU0479) (59) was used with MultiGeneBlast. Score = 2 for all genes present in contiguous cluster. Score = 1 for all genes in a cluster but intercalated with additional genes.

**PROTEINACEOUS ADHESINS**

**LapA adhesin:** PFL_0133 protein from *P. protegens* Pf-5 (NCBI accession AAY95545.1) (60) was used with BLAST. Score = 2 for homologues >60% identity.

**BapABCD adhesin:** The *bapABCD* gene cluster from *P. aeruginosa* PAO1 (PA1874-PA1877) (61) was used with MultiGeneBlast. Score = 2 for all genes present.

**Curli fimbriae:** The curli fimbriae gene cluster of *P. fluorescens* Pf0-1 (Pfl01_1982 - Pfl01_1993) (62) was used with MultiGeneBlast. Score = 2 for all genes present.

**PLANT-BACTERIAL COMMUNICATION**

**Auxin (IAA) catabolism:** The IAA catabolism gene cluster (*iacA-I*, NCBI accession EU360594.1) from *P. putida* strain 1290 (63) was used with MultiGeneBlast. Clusters containing all catabolic genes but lacking a homologue of the regulatory gene *iacR* were scored as 2; one cluster lacked *iacF* so was scored as 1.

**Phenyl acetic acid (PAA) catabolism:** The PAA catabolism gene cluster from from *P. protegens* Pf-5 (PFL_3128 - PFL_3140) (64) was used with MultiGeneBlast. Strains containing homologues of all genes were scored as 2. Some strains encoded the proteins across two distinct genomic loci (e.g. Ps673) but all proteins had high homology to *P. protegens* Pf-5 proteins (>70% identity) so were still scored as 2.

**1-Aminocyclopropane-1-carboxylate (ACC) deaminase:** AcdS from *P. fluorescens* F113 (NCBI accession AEV63500.1) (65) was used with BLAST. Score = 2 for homologues >70% identity.

**3-hydroxybutanone (acetoin) catabolism 1:** The acetoin catabolism gene cluster (*acoA-C*, *acoX*, *adh*) from *P. protegens* Pf-5 (PFL_2168 - PFL_2172) (66) was used with MultiGeneBlast. Score = 2 for all genes present in cluster. Score = 1 for clusters that lacked a homologue of AcoX.

**Acetoin catabolism 2:** Acetoin reductase from *P. fluorescens* A506 (NCBI accession AFJ57022.1) was used with BLAST. Score = 2 for homologues >70% identity.

**SECRETION SYSTEMS**

**Type II secretion system (T2SS):** The T2SS gene cluster (*gspCDEFGHIJKLM*) of *P. fluorescens* SBW25 (PFLU2415 - PFLU2425) (67) was used with MultiGeneBlast. Some genomes contain all genes within the same locus (score = 2), whereas some seem to only contain DEFGH and sometimes one pseudolipin, so are scored 1. Some (e.g. Ps664 and Ps720) have two authentic systems and others have a complete SBW25-like system as well as a smaller DEFGH-like clusters.

**Type III secretion system (T3SS):** The T3SS gene cluster of *P. fluorescens* SBW25 (PFLU0708, PFLU0710 - PFLU0727) was used with MultiGeneBlast. Score = 2 when homologues of all genes were found in a gene cluster. A score of 2 was also provided if homologues of the effector (PFLU0708) were not encoded in the gene cluster. Some gene clusters feature additional genes that could reflect additional components of the secretion system absent from SBW25. For example, the gene cluster in Ps664 included a Type III secretion ATP synthase (HrcN), a hypothetical protein and type III secretion protein (HrcV).

**Type VI secretion system (T6SS):** The T6SS gene cluster of *P. fluorescens* A506 (PflA506_2406 - PflA506_2421) was used with MultiGeneBlast. Score = 2 when homologues of all genes were found in a gene cluster. Some gene clusters feature additional genes that could reflect additional components of the secretion system absent from A506.

**EXOENZYMES**

**Pectin lyase 1:** Protein PFLU2293 from *P. fluorescens* SBW25 was used in BLAST analysis. Clear distinction between high identity (>90%, Score = 2) and low identity (<30%, score = 0) proteins.

**Pectin lyase 2:** Protein PFLU2269 from *P. fluorescens* SBW25 was used in BLAST analysis. Score = 2 with >80% identity.

**Pectate lyase:** Protein PFLU3229 from *P. fluorescens* SBW25 was used in BLAST analysis. Score = 2 with >80% identity. Strains such as Ps706 have high homology across to have “split” proteins and were scored as 1.

**Chitinase ChiC:** Protein PFL_2091 from *P. protegens* Pf-5 was used in BLAST analysis. Score = 2 with >50% identity. This cut-off retained proteins annotated as “Chitinase D”.

**Chitinase class 1:** Protein PSF113_1189 from *P. fluorescens* F113 was used in BLAST analysis. Score = 2 with >60% identity. This cut-off retained proteins annotated as "Chitinase class I".

**Extracellular alkaline metalloprotease AprA:** Protein PFL_3210 from *P. protegens* Pf-5 was used in BLAST analysis. Score = 2 with >80% identity; score = 1 with >65% identity.

**LipA lipase:** Protein PFLU0569 from *P. fluorescens* SBW25 was used in BLAST analysis. Score = 2 with >75% identity.

**LipB lipase:** Protein PFLU3141 from *P. fluorescens* SBW25 was used in BLAST analysis. Score = 2 with >80% identity; score = 1 with >65% identity.

### TOXINS

**Insecticidal toxin complex (Tc) gene clusters:** To search for putative insecticidal toxin complex (Tc) gene clusters, the examples reported by Rangel *et al*. (68) were used with MultiGeneBlast and/or BLAST analysis. Specifically, the following proteins were used:
Type I = *P. chlororapsis* 30–84 (Pchl3084_2947 and Pchl3084_2950)
Type II = *P. fluorescens* Q2-87 (PflQ2_0667–0670)
Type III = *P. fluorescens* Q8r1-96 (PflQ8_4696, PflQ8_4570–4571 and PflQ8_4580–4581)
Type IV = *P. fluorescens* Pf0-1 (Pfl01_0947–0948 and Pfl01_4453–4456)
Type V = *P. fluorescens* A506 (PflA506_3065–3068)
Type VI = *P. synxantha* BG33R (PseBG33_3799–3804)
Type II/V/VI Tc proteins provided comparable hits (due to sequence homology between components of these toxin systems) whose genes were arranged differently to the characterised examples. These were therefore grouped as a single genotype and scored as 1.

**HicAB toxin-antitoxin:** The HicAB proteins from *P. aeruginosa* PA1 *(*PA1S_06925 and PA1S_06920) (69) was used with MultiGeneBlast.

### ACCESSORY GENOME LOCI NOT FOUND IN THIS STRAIN COLLECTION

**Cytokinin:** The cytokinin isopentenyl transferase Ptz from *P. savastanoi* (NCBI accession P06619.1) (70) was used in a BLAST analysis.

**CdrA adhesin:** Protein PA4625 from *P. aeruginosa* PAO1 (71) was used in a BLAST analysis.

**N-acyl homoserine lactonase:** Protein BW979_RS17690 from *Pseudomonas* sp. A214 used in a BLAST analysis. This provided some hits, but all were <25% identity so were scored as 0.

**Fit toxin:** The Fit insect toxin cluster of *P. protegens* Pf-5 (PFL_2980 to PFL_2987) (72) was used with MultiGeneBlast.

**Insecticidal protein IPD072Aa:** Protein IPD072Aa from *P. chlororaphis* (NCBI accession KT795291.1) (73) was used in a BLAST analysis.

### SUPPLEMENTARY TABLES

**Table S1**. Bacterial strains used in this study.

| Strain | Description | Reference |
|---|---|---|
| SBW25 | *Pseudomonas fluorescens* SBW25 | (74) |
| LMG 2338 | *Pseudomonas* sp. LMG 2338 (NCPPB 387) | (75) |
| *E. coli* DH5α (Subcloning Efficiency™ DH5α™) | *E. coli endA*1, *hsdR*17(rK-mK+), *supE*44, *recA*1, *gyrA* (Nal$^r$), *relA*1, Δ(*lacIZYA-argF*)U169, *deoR*, Φ80*dlacΔ(lacZ)M15* | Thermo Fisher Scientific |
| Ps616 - Ps734 | 120 environmental *Pseudomonas* strains collected from RG Abrey Farms in Feb 2015 | This study |
| Ps831 - Ps950 | 120 environmental *Pseudomonas* strains collected from RG Abrey Farms in May 2015 | This study |
| IR1-1 - NS6-8 (see Supporting Dataset 1) | 192 environmental *Pseudomonas* strains collected from RG Abrey Farms in June 2017 | This study |
| Ps682Δvisc | *Pseudomonas* sp. Ps682 with deletion of a NRPS gene in the viscosin-like BGC | This study |
| Ps682::lux | *Pseudomonas* sp. Ps682 expressing *lux* operon | This study |
| Ps682Δvisc::lux | *Pseudomonas* sp. Ps682 Δviscosin expressing *lux* operon | This study |
| Ps619Δten | *Pseudomonas* sp. Ps619 with deletion of a NRPS gene in the tensin-like BGC | This study |
| Ps619Δhcn | *Pseudomonas* sp. Ps619 with deletion of the HCN BGC | This study |
| Ps619ΔtenΔhcn | *Pseudomonas* sp. Ps619 with deletions in tensin and HCN BGCs | This study |
| Ps619::lux | *Pseudomonas* sp. Ps619 expressing *lux* operon | This study |
| Ps619Δten::lux | *Pseudomonas* sp. Ps619Δtensin expressing *lux* operon | This study |
| Ps619Δhcn::lux | *Pseudomonas* sp. Ps619ΔHCN expressing *lux* operon | This study |
| Ps619ΔtenΔhcn::lux | *Pseudomonas* sp. Ps619ΔtensinΔHCN expressing *lux* operon | This study |

**Table S2**. Primers used in this study.

| No. | Name | Sequence |
|---|---|---|
| 1 | Viscosin up NdeI | GGAATTCCATATGTCACAGGACGATCTCCATTCGTTC |
| 2 | Viscosin up XbaI | GCTCTAGACCGTGGCCTGCGTATCGAACTG |
| 3 | Viscosin dn XbaI | GCTCTAGACTTGCAGCGGTTCGTCGAG |
| 4 | Viscosin dn BamHI | CGGGATCCGCAGTTCAGCGAATTATTGGCAG |
| 5 | Viscosin ext | GACTTACCAGGCAGGCAGGTTAC |
| 6 | Tensin up Fwd | CGCATATGGGCCGTCATGCCACCACG |
| 7 | Tensin up Rev | CGCCTAGGGTGGCGTCGTTGACCCTG |
| 8 | Tensin dwn Fwd | CGCCTAGGGCAGGATGTCGTGGCGATC |
| 9 | Tensin dwn Rev | CGTCTAGATCAAGTGAATGTGCTCGAACTATTGGCA |
| 10 | Tensin Int Screen Fwd | CCTGACGCCAGACCACTTG |
| 11 | TensinExtr_f | CGTTTGCCCTTCTTGAGTGA |
| 12 | Tens619InterRev | TTGGGAGAACAAGGTCGGTA |
| 13 | Pf_HCN_Up_Fwd | CGGAGCTCCCCCAATCACAGCAACAATAA |
| 14 | PfHCN_UP_Rev | CGTCTAGAGCCATCGAGTACCACCAAA |
| 15 | PfHCN_down_Fwd | CGTCTAGATGAAAGGCCAGATTCTACT |
| 16 | PfHCN_Down_R_Rev | CGCATATGTGTCAGTCAACCTATTCGT |
| 17 | SacB Fwd | GTTGATTGTTTGTCTGCGT |
| 18 | SacB Rev | TTTAGTTCTTTAGGCCCGT |
| 19 | PTS1f | CGGCAGGTATATGTGATGG |
| 20 | PTS1r | GTGAGAAATCACCATGAGTG |

**Table S3**. Plasmids used in this study.

| **Plasmids** | **Description** | **Reference** |
|---|---|---|
| pTS1 | pME3087 derivative containing a *sacB* counter-selection marker | (76) |
| pTS1-Δviscosin | Construct for viscosin deletion, fragments amplified with primers 1- 2 and 3-4, | This study |
| pTS1-Δtensin | Construct for tensin deletion, fragments amplified with primers 6-7 and 8-9, | This study |
| pTS1-Δ619HCN | Construct for HCN cluster deletion, fragments amplified with primers 13-14 and 15-16 | This study |
| pTNS2 | Tn7 transposase expression plasmid | (77) |
| pUC18-mini-Tn7T-Gm-lux | mini-Tn7 *luxCDABE* transcriptional fusion vector | (77) |

**Table S4**. NMR data for the isolated viscosin I (600 MHz, DMF-d7, 298 K). Atom numbering and spectra shown in Figures S19-S29.

| Number | δ$^{1}$H | δ$^{13}$C |
|---|---|---|
| **HDA**[a] | | |
| 1' | | 173.17 |
| 2' | H2 2.45 | 44.39 |
| 3' | H 4.01 | 68.10 |
| 4' | H2 1.47 | 38.13 |
| 5' | Ha 1.46 | 25.92 |
| 5' | Hb 1.35 | |
| 6' | H2 1.28 | 29.44 |
| 7' | H2 1.28 | 29.44 |
| 8' | H2 1.27 | 31.94 |
| 9' | H2 1.28 | 22.67 |
| 10' | H3 0.87 | 13.85 |
| **Leu1** | | |
| 1NH | H nd[b] | |
| 1a | H 4.16 | 52.34 |
| 1b | H' 1.46 | 39.60 |
| 1c | H 1.73 | 24.84 |
| 1d | H3 0.93 | 23.09 |
| 1e | H3 0.91 | 21.41 |
| 1CO | | 173.59 |
| **Glu2** | | |
| 2NH | H nd[b] | |
| 2a | H 4.16 | 56.87 |
| 2b | H2 2.02 | 27.44 |
| 2c | H2 2.24 | 34.82 |
| 2d | | 177.25 |
| 2CO | | 173.09 |
| **Thr3** | | |
| 3NH | H 8.76 | |
| 3a | H 4.38 | 59.73 |
| 3b | H 5.41 | 70.28 |
| 3c | H3 1.34 | 17.37 |
| 3CO | | 173.08 |
| **Val4** | | |
| 4NH | H 8.25 | |
| 4a | H 3.68 | 63.92 |
| 4b | H 2.32 | 29.36 |
| 4c | H3 1.02 | 20.14 |
| 4d | H3 0.95 | 19.15 |
| 4CO | | 173.09 |

| Number | δ$^{1}$H | δ$^{13}$C |
|---|---|---|
| **Leu5** | | |
| 5NH | H 7.08 | |
| 5a | H 4.24 | 53.39 |
| 5b | H 1.88 | 41.34 |
| 5b | H 1.65 | |
| 5c | H 1.75 | 24.72 |
| 5d | H3 0.86 | 20.82 |
| 5e | H3 0.99 | 22.90 |
| 5CO | | 173.08 |
| **Ser6** | | |
| 6NH | H nd[b] | |
| 6a | H 4.26 | 56.86 |
| 6b | H' 4.07 | 63.10 |
| 6b | H" 3.86 | |
| 6CO | | 171.38 |
| **Leu7** | | |
| 7NH | H nd[b] | |
| 7a | H 4.24 | 54.61 |
| 7b | H' 1.47 | 40.10 |
| 7b | H" 1.82 | |
| 7c | H 1.75 | 24.84 |
| 7d | H3 0.86 | 20.82 |
| 7e | H3 0.88 | 22.90 |
| 7CO | | 173.09 |
| **Ser8** | | |
| 8NH | H 8.03 | |
| 8a | H 4.43 | 56.27 |
| 8b | H' 3.93 | 61.92 |
| 8b | H" 3.74 | |
| 8CO | | 171.37 |
| **Ile9** | | |
| 9NH | H 6.98 | |
| 9a | H 4.54 | 56.46 |
| 9b | H 1.95 | 36.55 |
| 9c | H' 1.29 | 24.48 |
| 9c | H" 1.05 | |
| 9d | H3 0.89 | 11.58 |
| 9e | H3 0.86 | 15.52 |
| 9CO | | 169.78 |

[a.] HDA = 3-hydroxydecanoic acid
[b.] nd = not determined

### SUPPLEMENTARY FIGURES

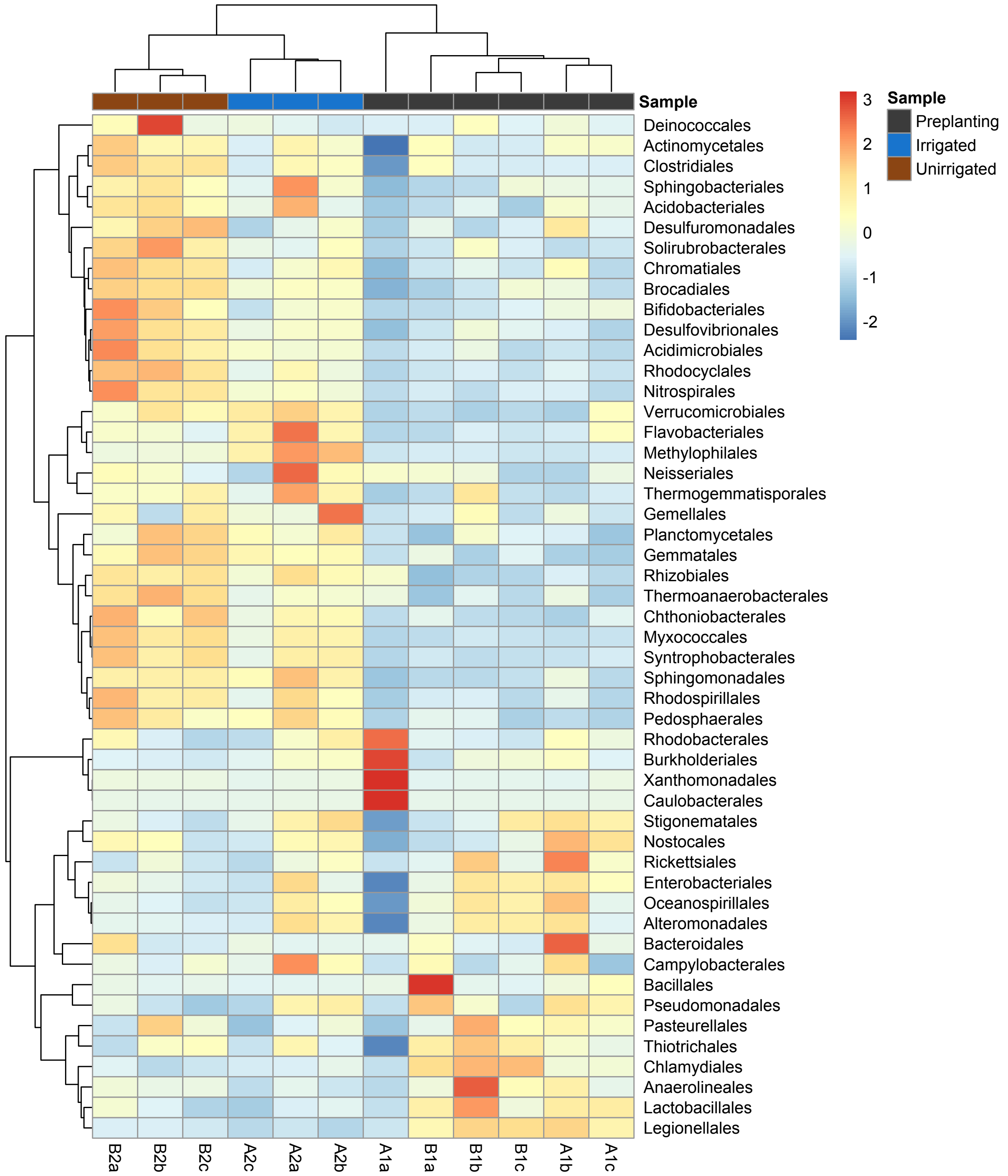


**Figure S1** Heatmap of the 50 most abundant bacterial orders across all samples (unclassified reads were removed from this analysis). The heatmap is scaled by z-score for each row, and samples (columns) and bacterial orders (rows) are clustered by Pearson correlation. The heatmap was generated using Pheatmap (https://CRAN.R-project.org/package=pheatmap) in R 3.5.1.

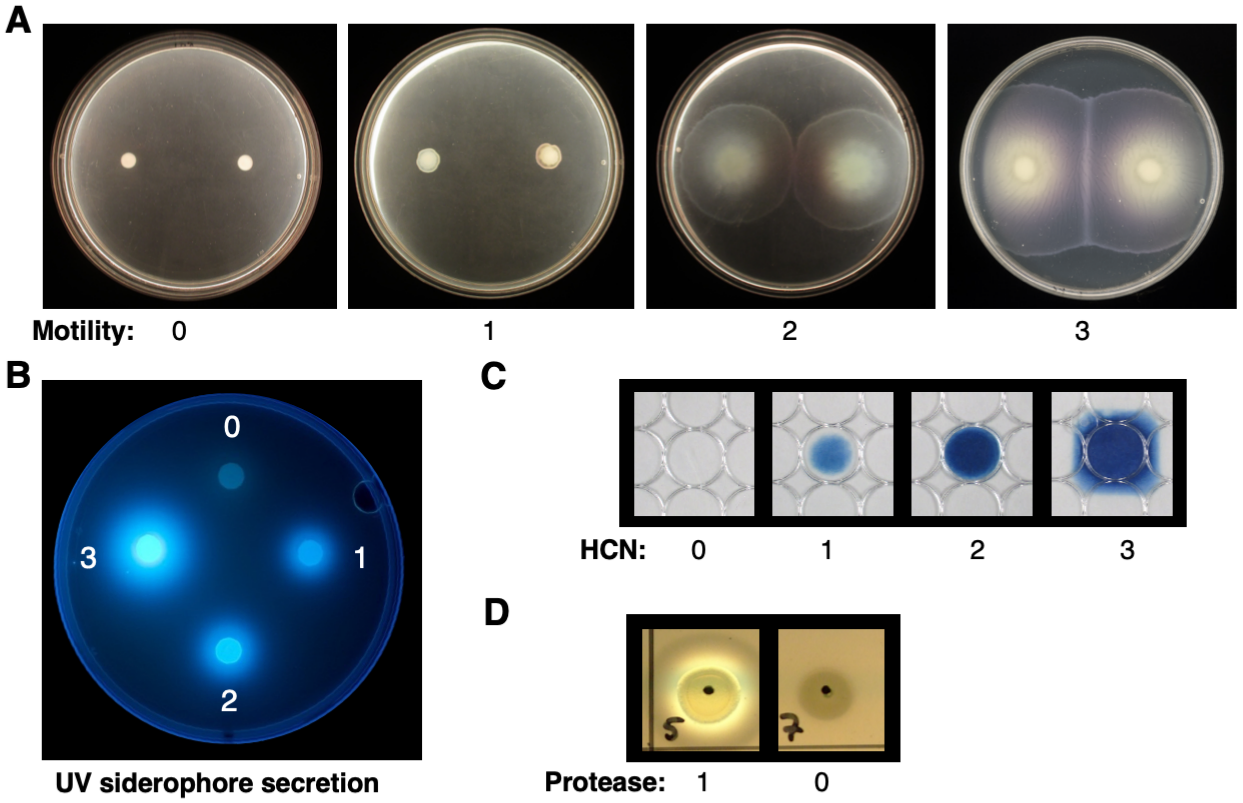


**Figure S2** Representative images of *Pseudomonas* phenotypes. (A) Motility; (B) UV siderophore secretion; (C) HCN production; (D) protease production.

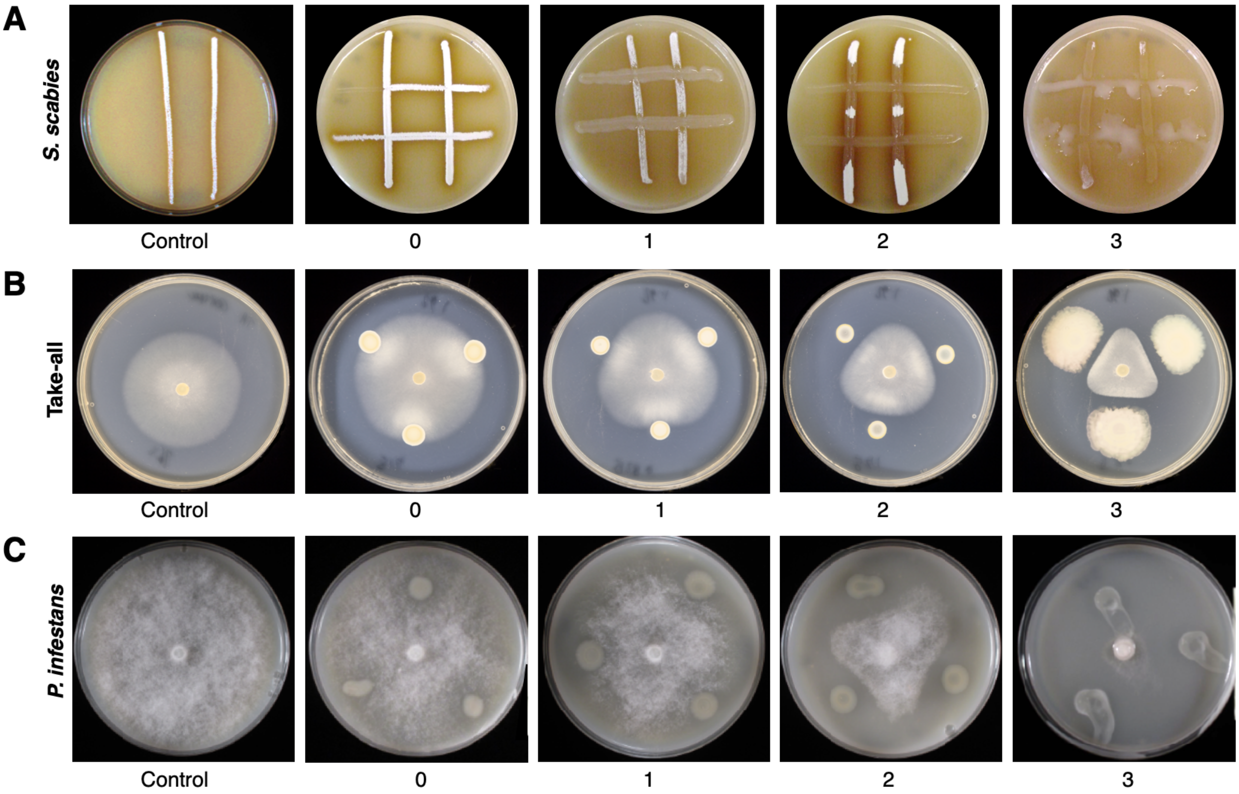


**Figure S3** Representative images of on-plate suppressive activity of *Pseudomonas* isolates. (A) Versus *S. scabies*; (B) Versus *Gaeumannomyces graminis pv. tritici* (Take-all); (C) Versus *P. infestans*.

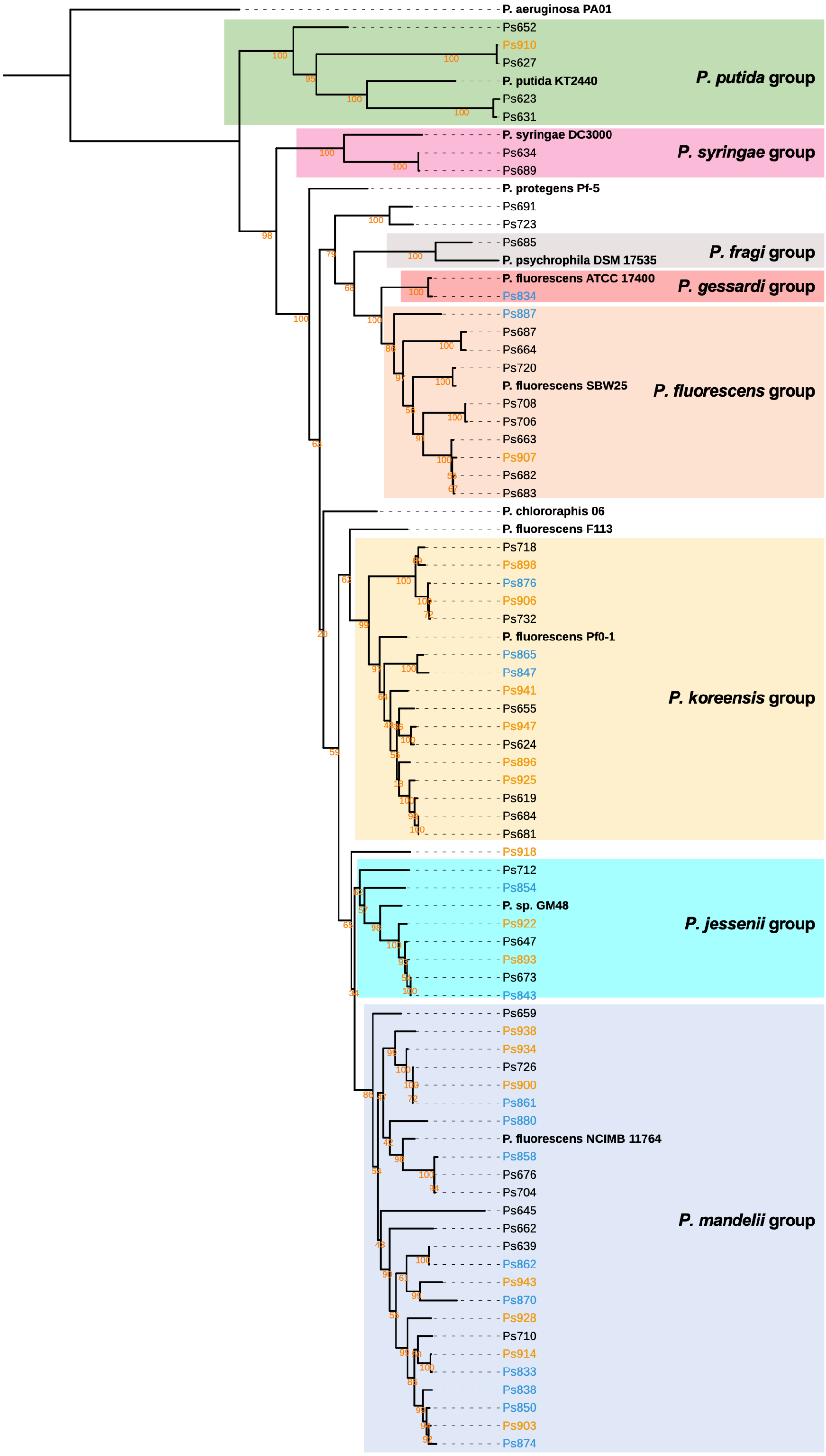


**Figure S4** Maximum likelihood tree of full-length *gyrB* nucleotide sequences of *Pseudomonas* strains with representatives of the eight *Pseudomonas* phylogenomic groups defined by Garrido-Sanz *et al.* (78). Black names represent pre-planting isolates, blue names represent isolates from irrigated soil and orange names indicate isolates from unirrigated soil. Bootstrap values are shown for each branch in orange text. Sequences were aligned with MUSCLE (79) and the tree was inferred using RAxML (80). The tree was visualised using iTOL (81). An interactive version of the tree annotated with features shown in Figure 2 is available at: https://itol.embl.de/tree/9024318186586716333395 79

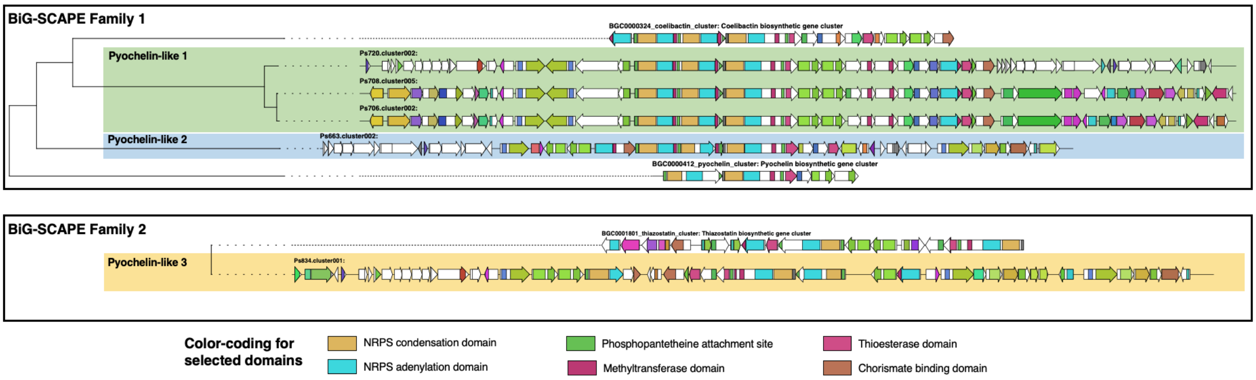


**Figure S5** Comparison of characterized pyochelin-like BGCs versus pyochelin-like BGCs identified in this work. BGCs were retrieved from MIBiG (https://mibig.secondarymetabolites.org) (2) for pyochelin (BGC0000412), coelibactin (BGC0000324) and thiazostatin (BGC0001801). BGC homology was assessed using BiG-SCAPE (https://git.wageningenur.nl/medema-group/BiG-SCAPE/) (82) in glocal mode with a cut-off of 0.75. Two figure shows two distinct families.

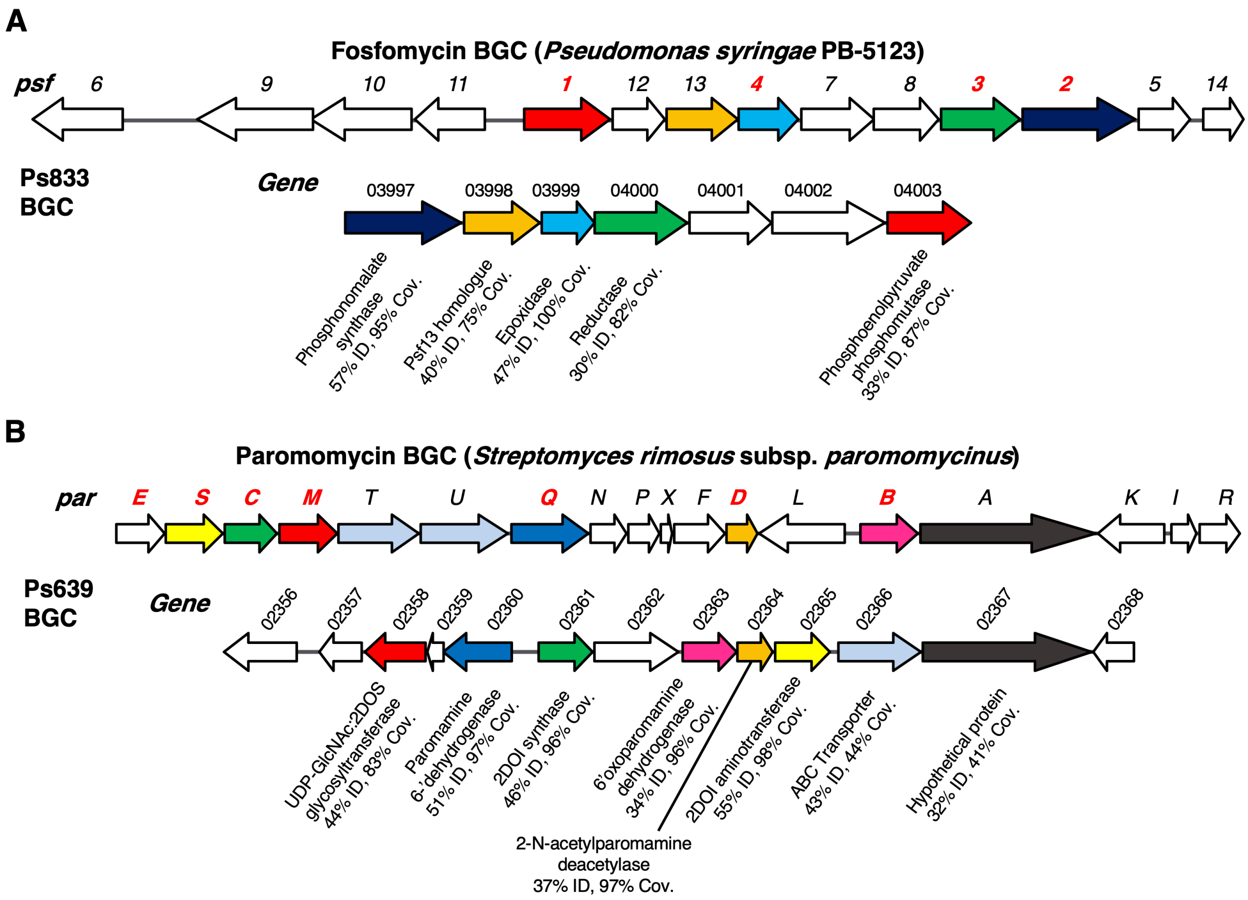


**Figure S6** Examples of BGCs not previously characterized in *P. fluorescens*. (A) Fosfomycin-like BGC in strain Ps833. Comparison to the *psf* BGC in *P. syringae* PB-5123 is shown, where color-coding represents homologous genes (identity and coverage values relate to encoded proteins). Gene numbers colored in red have been experimentally characterized and encode enzymes that catalyze key biosynthetic steps (37, 83). (B) Aminoglycoside-like BGC in strain Ps639. Comparison to the *par* BGC in *S. rimosus* is shown, where color-coding represents homologous genes (identity and coverage values relate to encoded proteins). Gene numbers colored in red are predicted to be required for the biosynthesis of the minimal aminoglycoside, neamine (34).

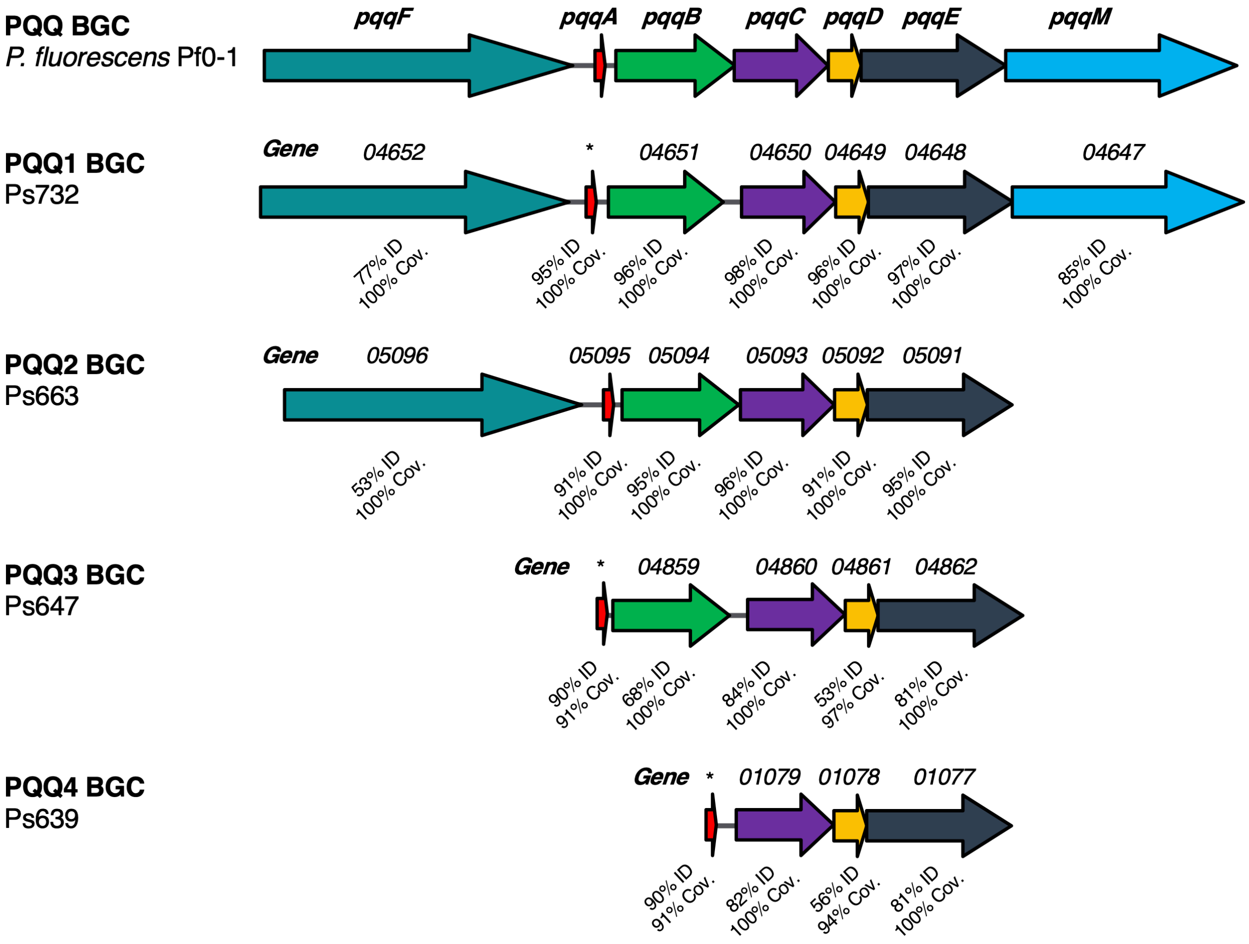


**Figure S7** Different PQQ BGCs identified in this study. Comparison to the *pqq* BGC in *P. fluorescens* Pf0-1 is shown, where color-coding represents homologous genes and % identity/coverage indicate how similar the encoded proteins are to the Pf0-1 PQQ proteins. * = gene not annotated but *pqqA* homologue identified by tblastn analysis.

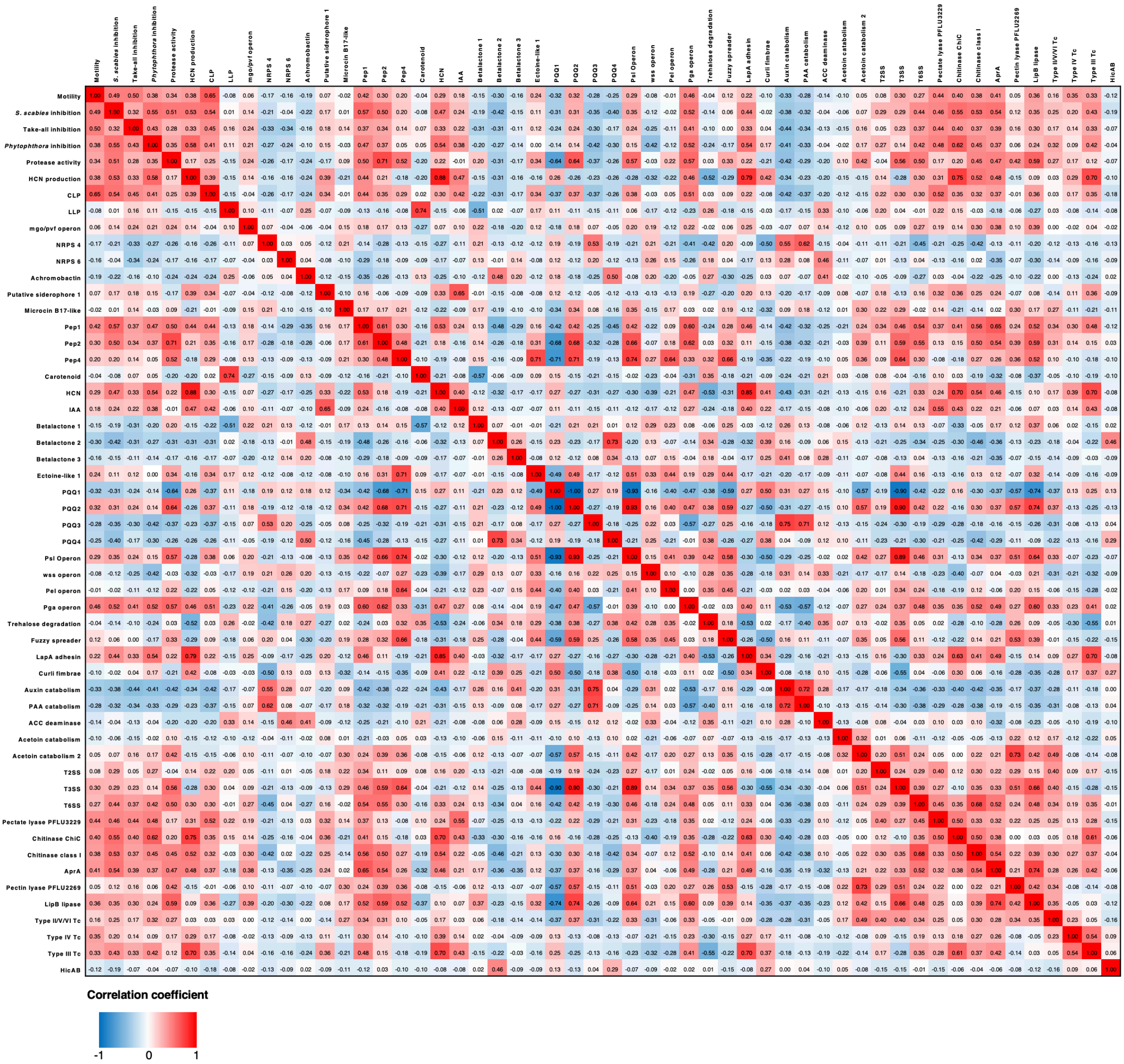


**Figure S8** Heatmap showing Pearson correlation coefficients of phenotypes and genotypes across the sequenced *Pseudomonas* isolates. For clarity, any phenotype or genotype that had fewer than 5 or greater than 65 representatives (total number of strains = 69) was removed from this analysis.

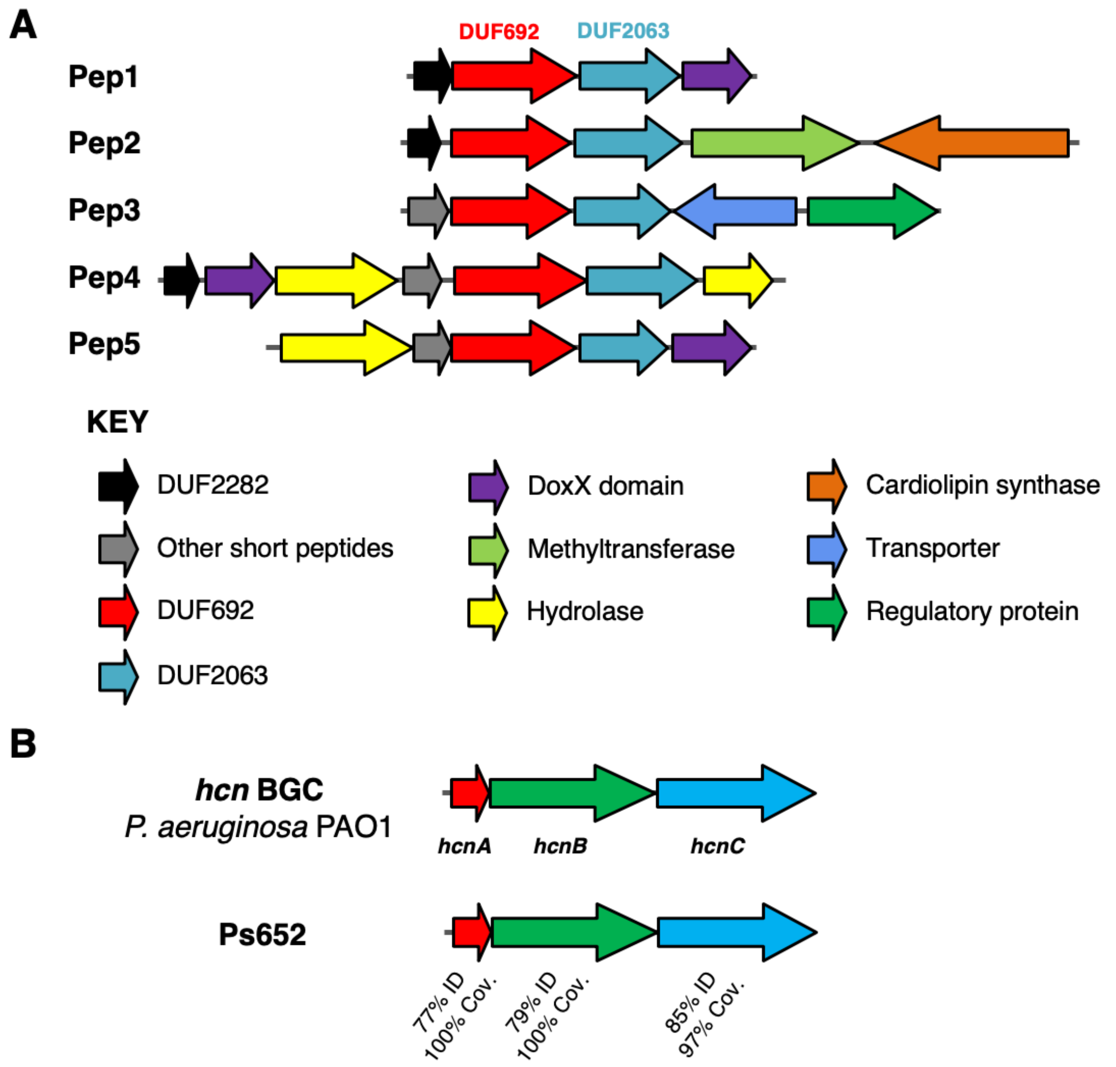


**Figure S9** BGCs that correlate with *S. scabies* inhibition. (A) Putative Pep BGCs. (B) HCN BGC compared to the characterized *P. aeruginosa* PAO1 BGC (28).

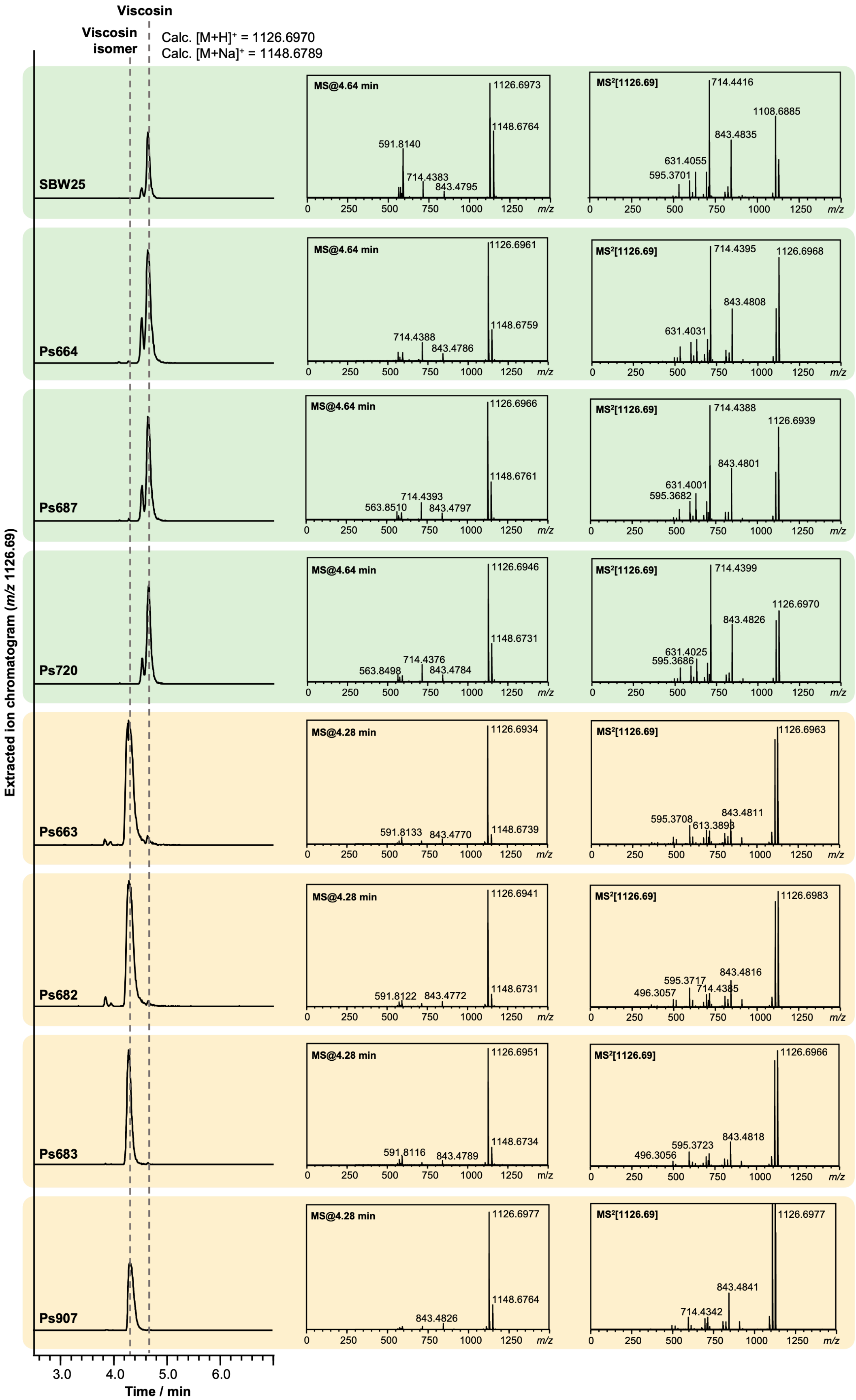


**Figure S10** LC-MS spectra showing that some strains produce viscosin (identical retention time and MS/MS fragmentation to viscosin produced by *P. fluorescens* SBW25; green boxes) and some produce an isomer with a distinct retention time and MS/MS fragmentation pattern (yellow boxes).

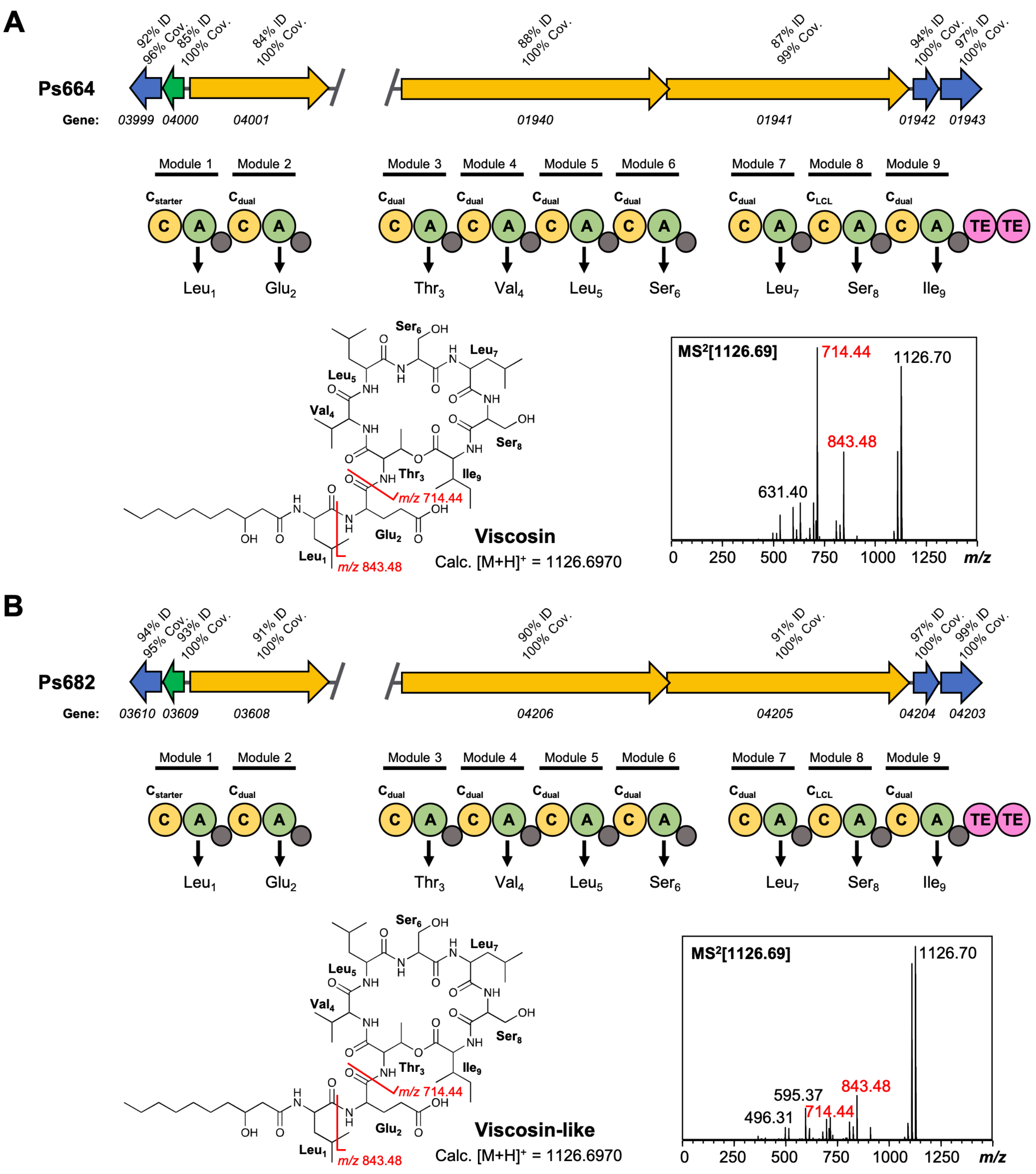


**Figure S11** Comparison of viscosin (A) and viscosin-like (B) BGCs. Genes encoding regulatory proteins are green, transporter genes are blue, and NRPS genes are yellow. The break in the BGC indicates that the BGC is found in two distinct genomic loci. Homology to the *Pseudomonas fluorescens* SBW25 viscosin BGC is shown in both panels A and B. The NRPS organization is displayed, where C = condensation domain, A = adenylation domain, TE = thioesterase, and small grey circles are peptidyl carrier protein domains. Amino acids incorporated by each module are displayed, along with predicted condensation domain specificity. MS/MS data showing characteristic viscosin fragmentation is also shown.

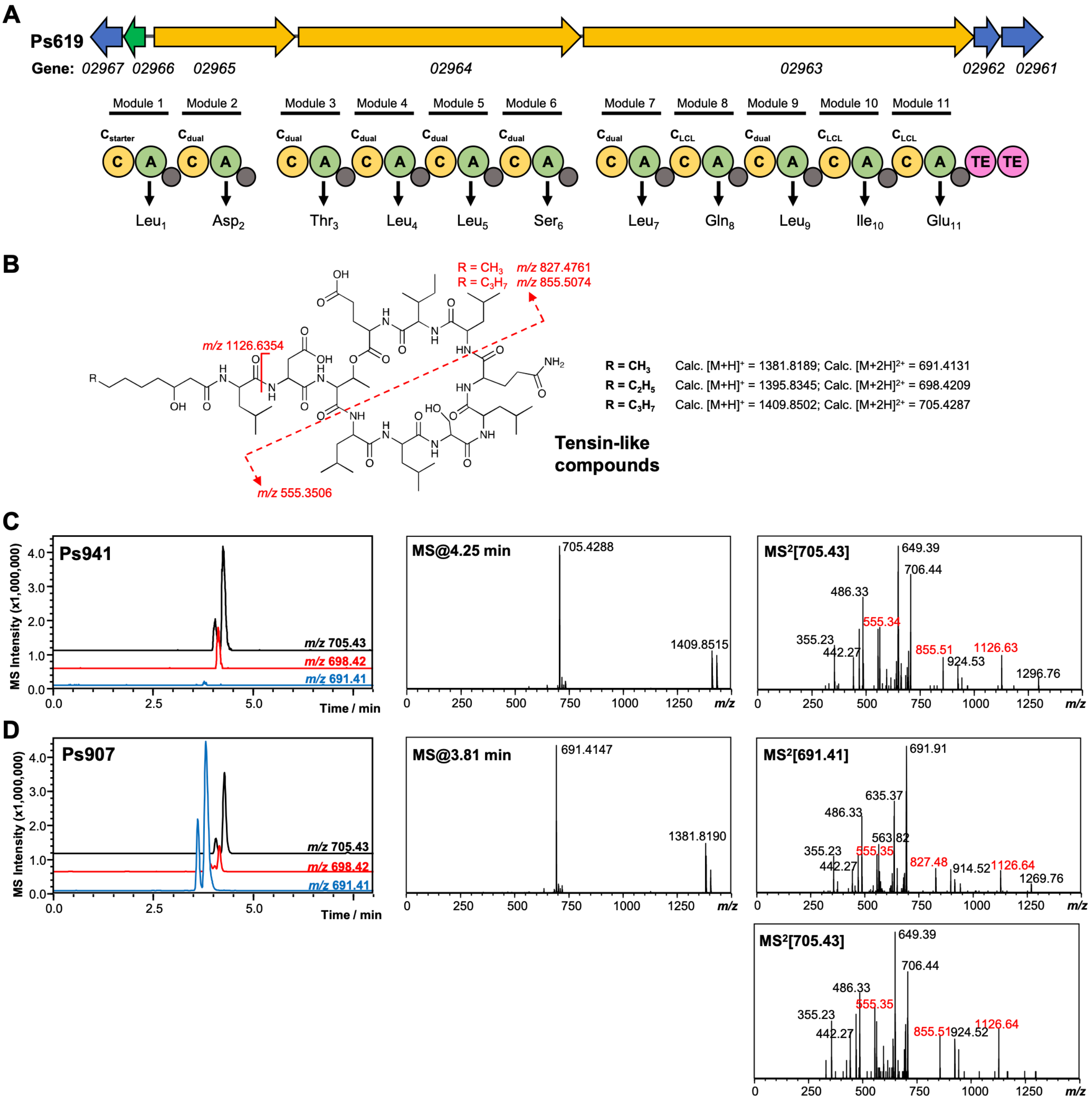


**Figure S12** Genetic and LC-MS analysis of tensin-like compounds. (A) Organisation of the CLP NRPS BGC from Ps619. Gene and NRPS domain colours are the equivalent to Figure 5 of the main paper. Amino acids predicted to be incorporated by each module are displayed, along with predicted condensation domain specificity. (B) Predicted structure, masses and key MS/MS fragments for tensin-like compounds. (C) Representative LC-MS spectra for another producer of a tensin-like molecule, where the major product $[M+2H]^{2+}$ = 705.43. Extracted ion chromatograms are shown for *m/z* values relating to different acyl chain lengths. Characteristic MS/MS fragments are highlighted in red. (D) LC-MS spectra of tensin-like compounds produced by Ps907, where the major product $[M+2H]^{2+}$ = 691.41. The presence of characteristic MS/MS fragments, as well as a smaller amount of $[M+2H]^{2+}$ = 705.43, is consistent with a shorter acyl chain (see panel B).

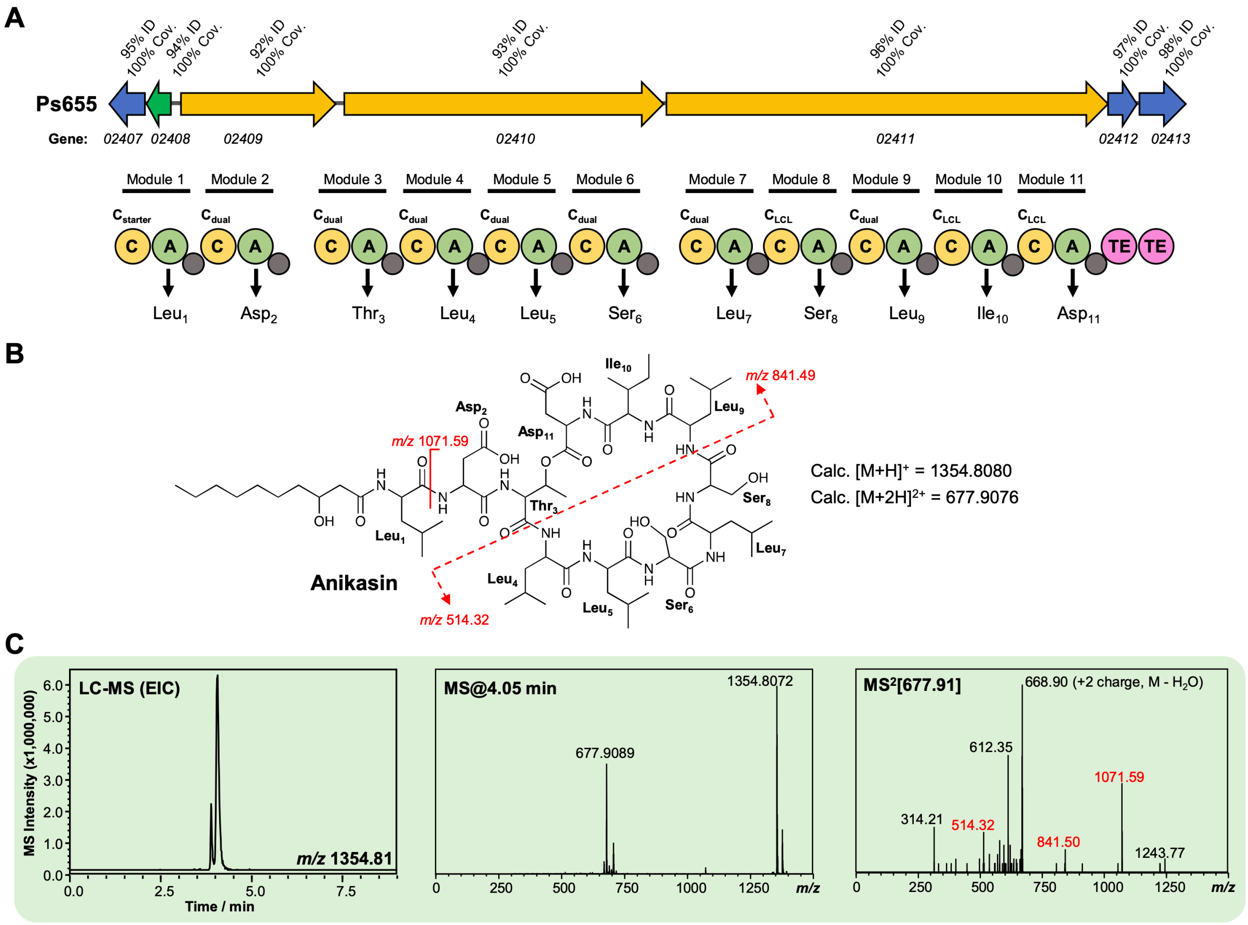


**Figure S13** Characterization of an anikasin-like lipopeptide from strain Ps655. (A) Gene cluster listing the homology to the *Pseudomonas fluorescens* HKI0770 anikasin BGC (MiBIG BGC0001509). The NRPS organization is displayed (color-coding and nomenclature as in Figure S11) and amino acids incorporated by each module are displayed, along with predicted condensation domain specificity. (B) Structure of anikasin showing calculated masses for observed MS/MS fragmentation. (C) LC-MS data showing production of an anikasin-like molecule by Ps655 (EIC = extracted ion chromatogram). Red MS/MS fragments relate to fragments shown in panel B and are equivalent to the tensin fragments shown in Figure S12. A variety of isomers of anikasin have been reported (84), such as arthrofactin and pholipeptin.

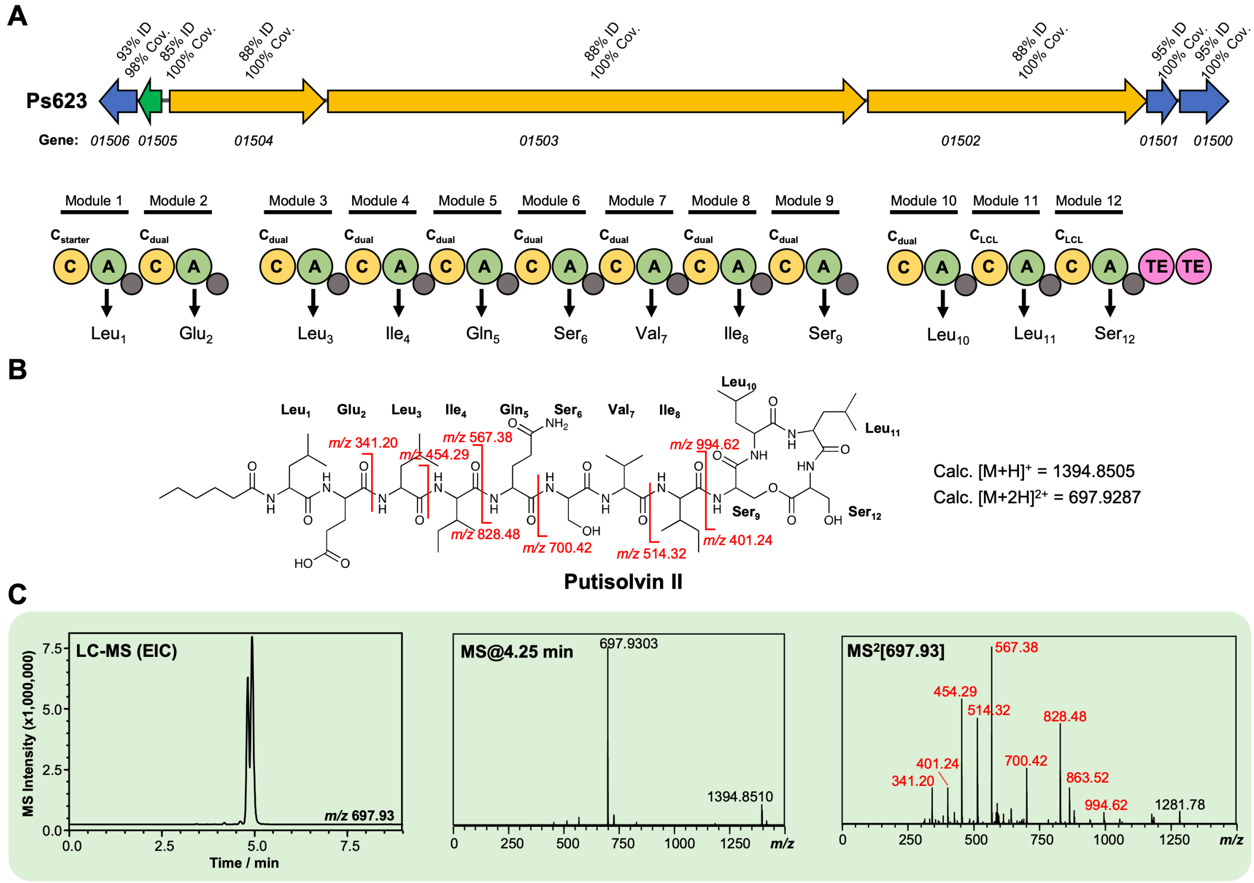


**Figure S14** Characterization of a putisolvin II-like lipopeptide from strain Ps623. (A) Gene cluster listing the homology to the *Pseudomonas putida* PCL1445 putisolvin BGC (MiBIG BGC0000411). The NRPS organization is displayed (color-coding and nomenclature as in Figure S11) and amino acids incorporated by each module are displayed, along with predicted condensation domain specificity. (B) Structure of putisolvin II showing calculated masses for observed MS/MS fragmentation. (C) LC-MS data showing production of a putisolvin II-like molecule by Ps623 (EIC = extracted ion chromatogram). Red MS/MS fragments relate to fragments shown in panel B.

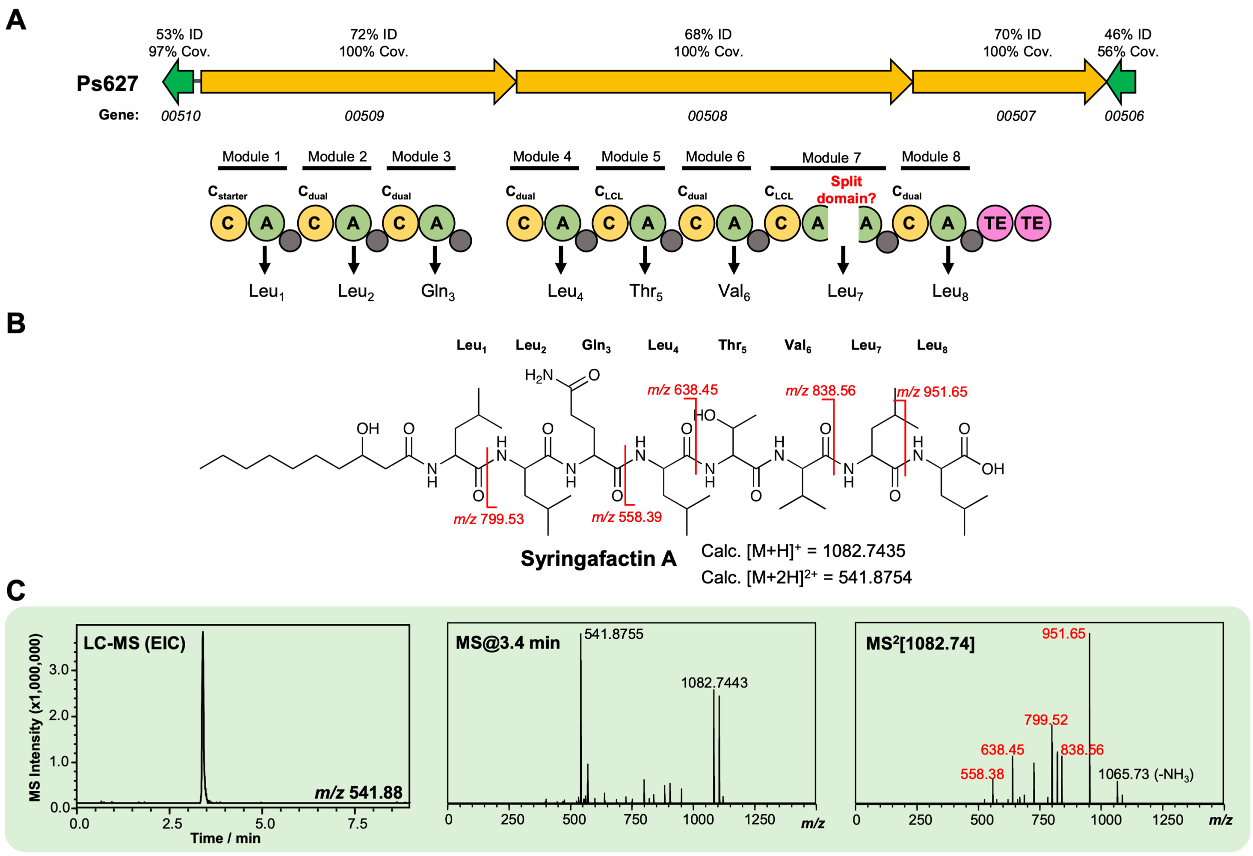


**Figure S15** Characterization of a syringafactin-like lipopeptide from strain Ps627. (A) Gene cluster listing the homology to the *P. syringae* DC3000 syringafactin BGC (MiBIG BGC0000435). The NRPS organization is displayed (color-coding and nomenclature as in Figure S11) with an unexpected “split” A domain. Further experiments are needed to determine whether this is real or a sequencing artefact. Amino acids incorporated by each module are displayed, along with predicted condensation domain specificity. (B) Structure of syringafactin A showing calculated masses for observed MS/MS fragmentation. (C) LC-MS data showing production of a syringafactin-like molecule by Ps627 (EIC = extracted ion chromatogram). Red MS/MS fragments relate to calculated fragments shown in panel B.

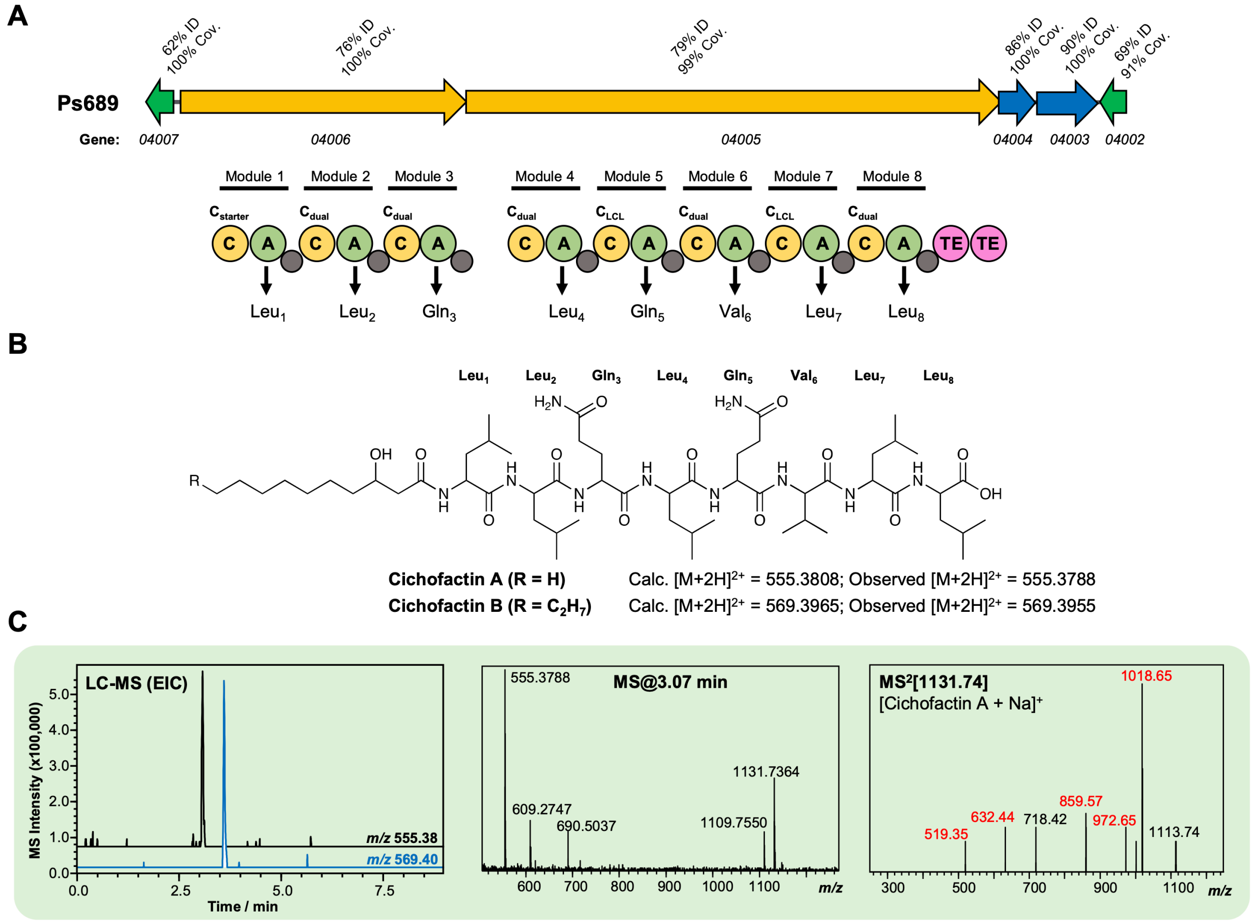


**Figure S16** Characterization of a cichofactin-like lipopeptide from strain Ps689. (A) Gene cluster listing the homology to the *Pseudomonas cichorii* cichofactin BGC (MiBIG BGC0000323). The NRPS organization is displayed (color-coding and nomenclature as in Figure S11). Amino acids incorporated by each module are displayed, along with predicted condensation domain specificity. (B) Structure of cichofactins showing calculated and observed $[M+2H]^{2+}$ values. (C) LC-MS data showing production of a cichofactin-like molecules by Ps689 (EIC = extracted ion chromatogram). Red MS/MS fragments relate to fragments reported for [cichofactin A + Na]$^{+}$ in Pauwelyn *et al*. (85).

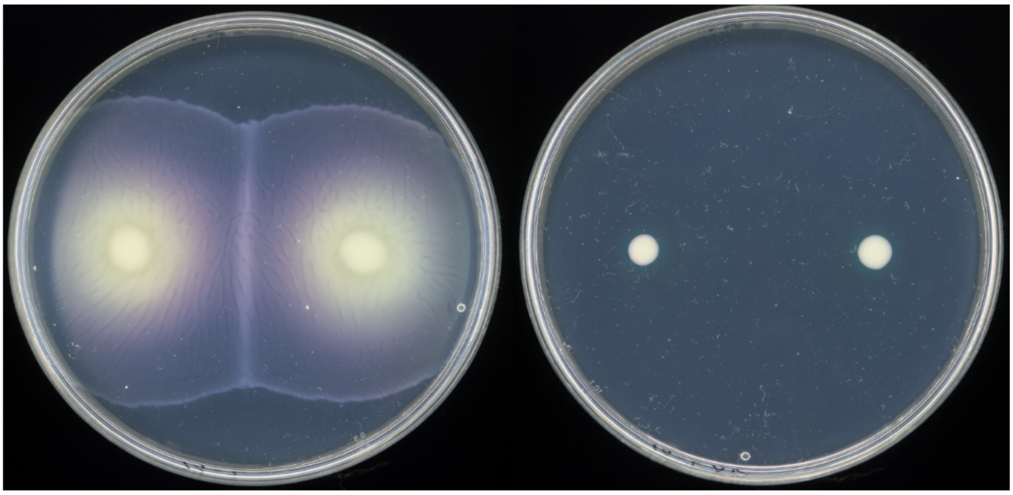


**Figure S17** Swarming motility of wild type Ps682 and Ps682 Δ*visc*.

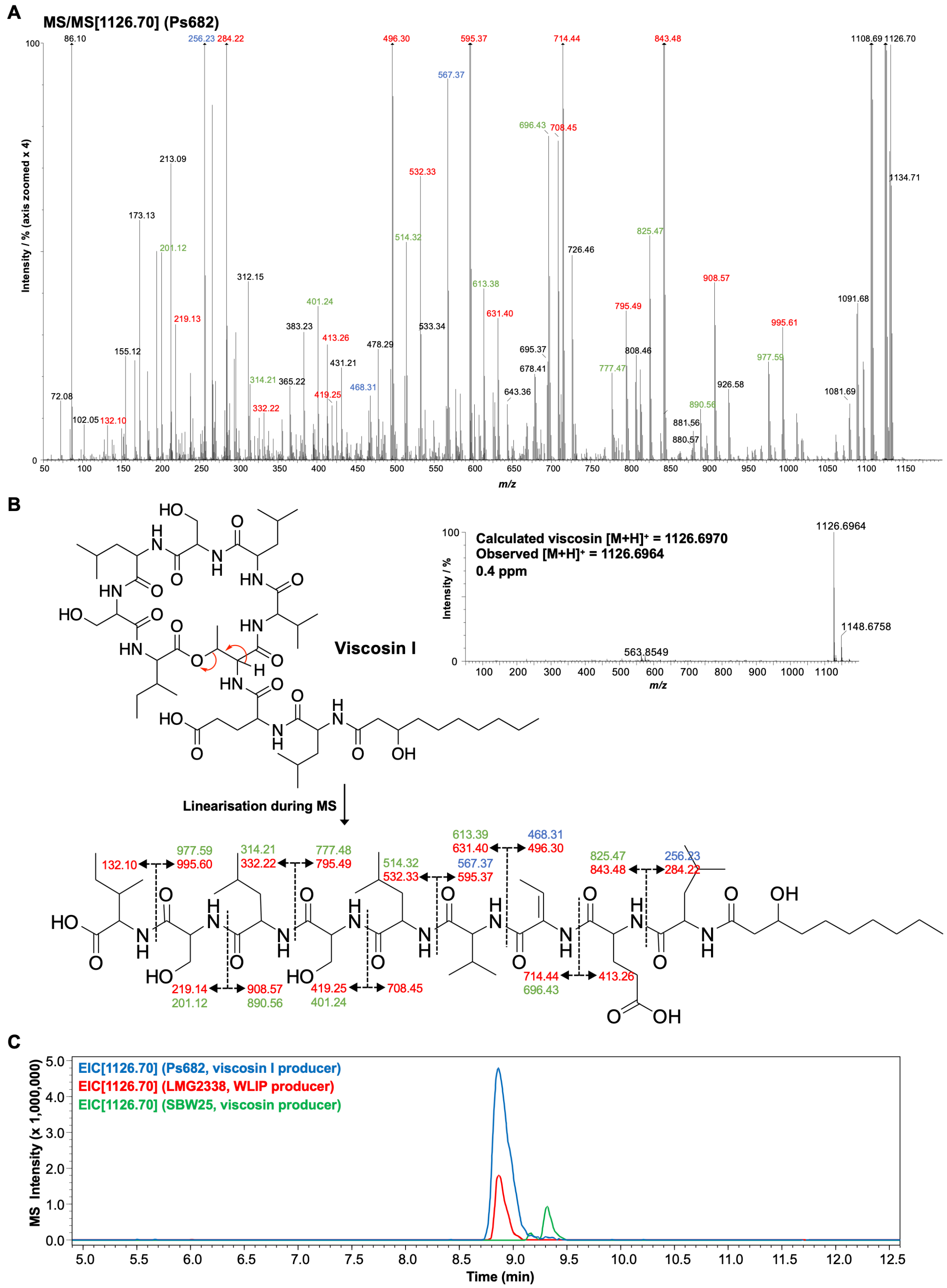


**Figure S18** Detailed MS analysis of viscosin-like molecule produced by Ps682. (A) MS/MS spectrum with annotated peaks colour-coded (intensity zoomed x4 for clarity). Red labels = b and y fragments; green labels = loss of water from b/y fragments; blue labels = a fragments. (B) Viscosin I accurate mass spectrum plus MS/MS fragment annotations. (C) LC-MS comparison of Ps682 (viscosin I) versus *Pseudomonas* producers of WLIP and viscosin. EIC = extracted ion chromatogram.

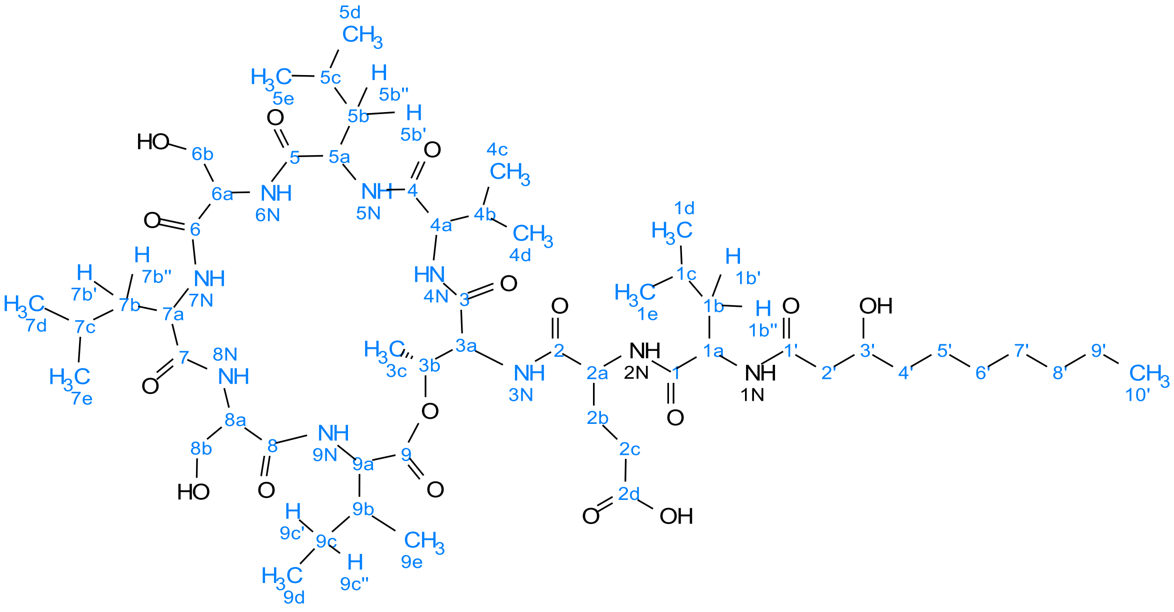


**Figure S19** Atom numbering for viscosin I NMR annotation.

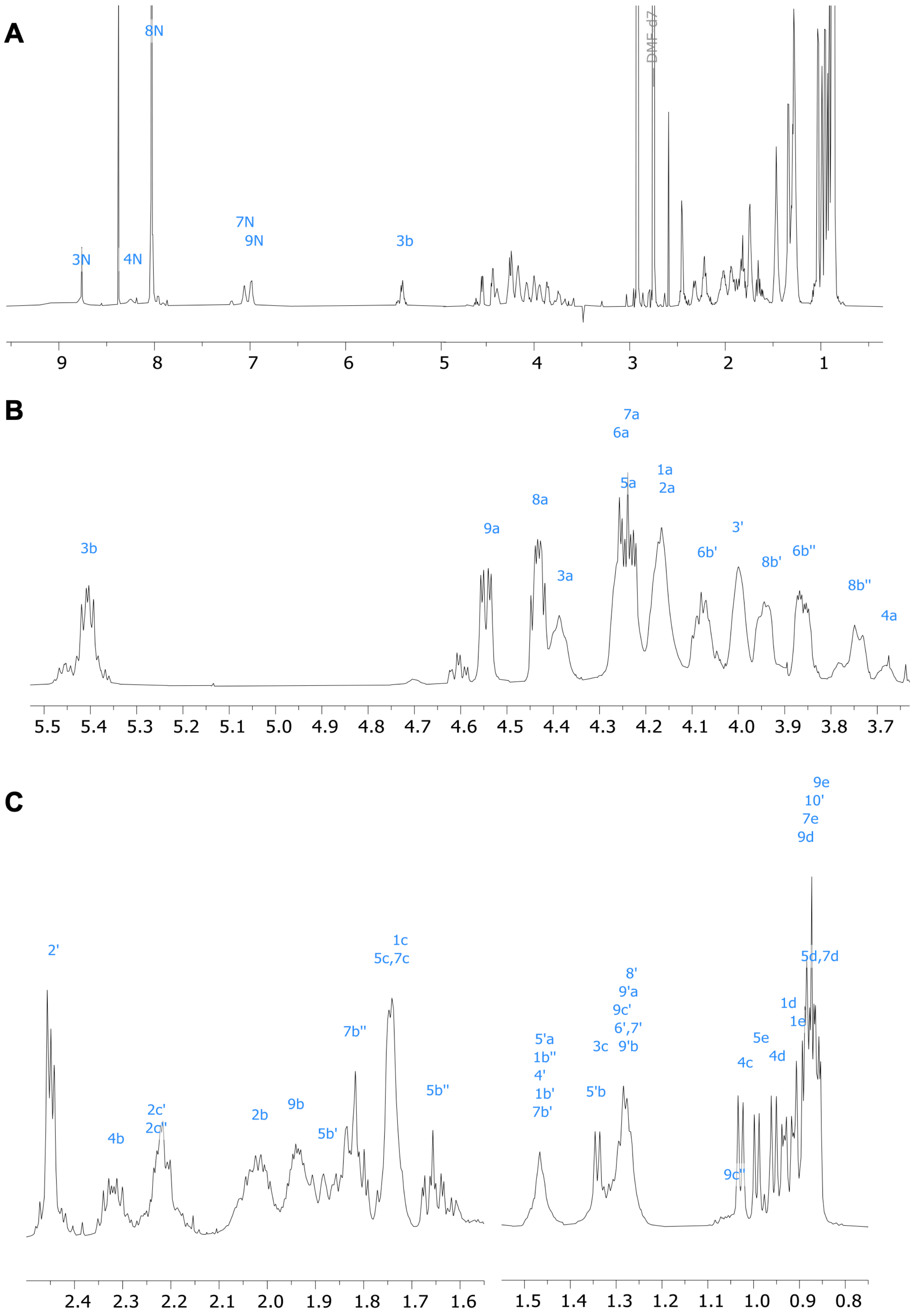


**Figure S20** [1]H NMR spectrum of viscosin I (600 MHz, DMF-d7, 298K). (A) Full spectrum, (B) and (C) show expanded regions of the spectrum.

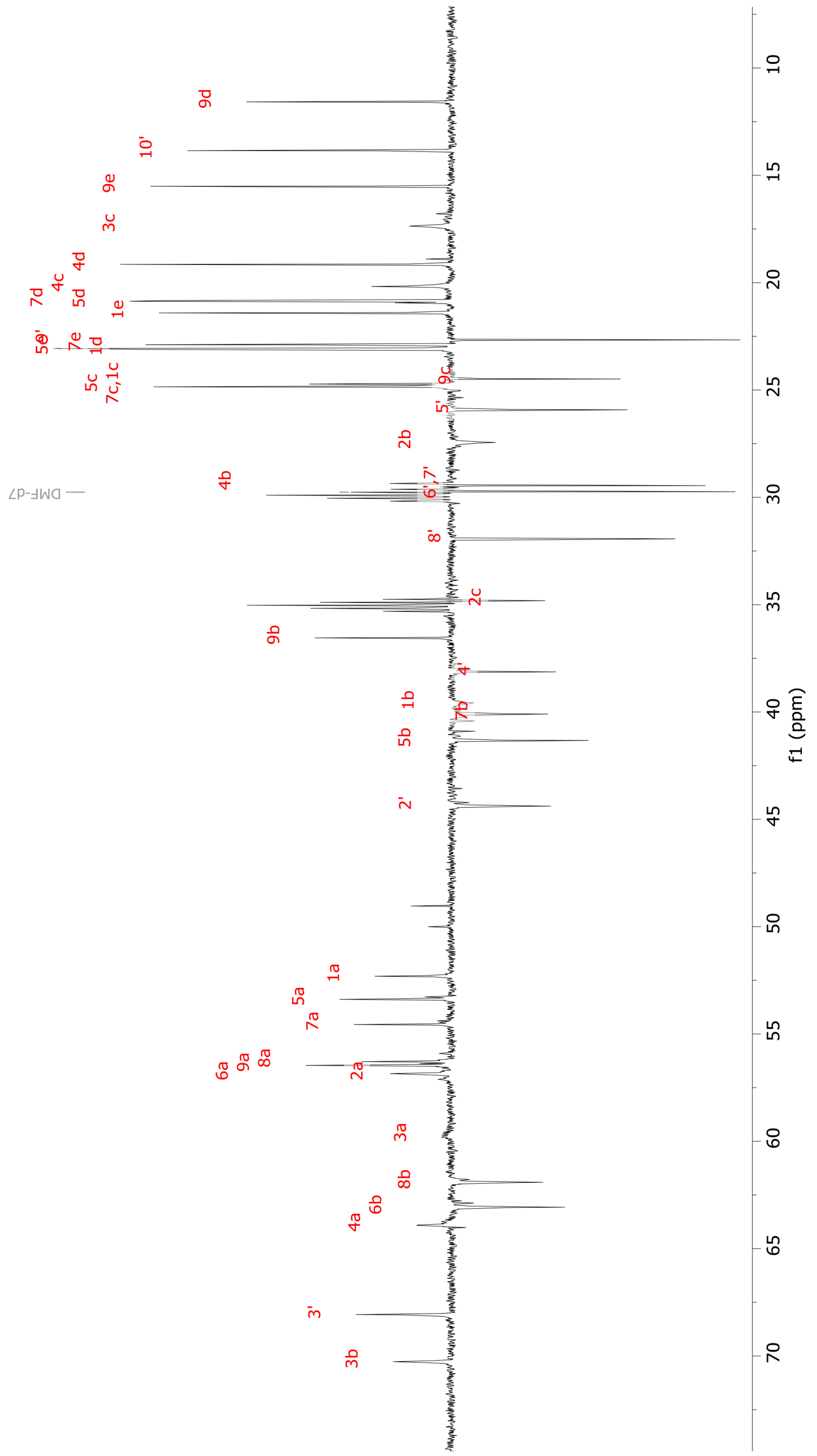


**Figure S21** $^{13}$C NMR spectrum (DEPT135) of viscosin I (600 MHz, DMF-d7, 298K).

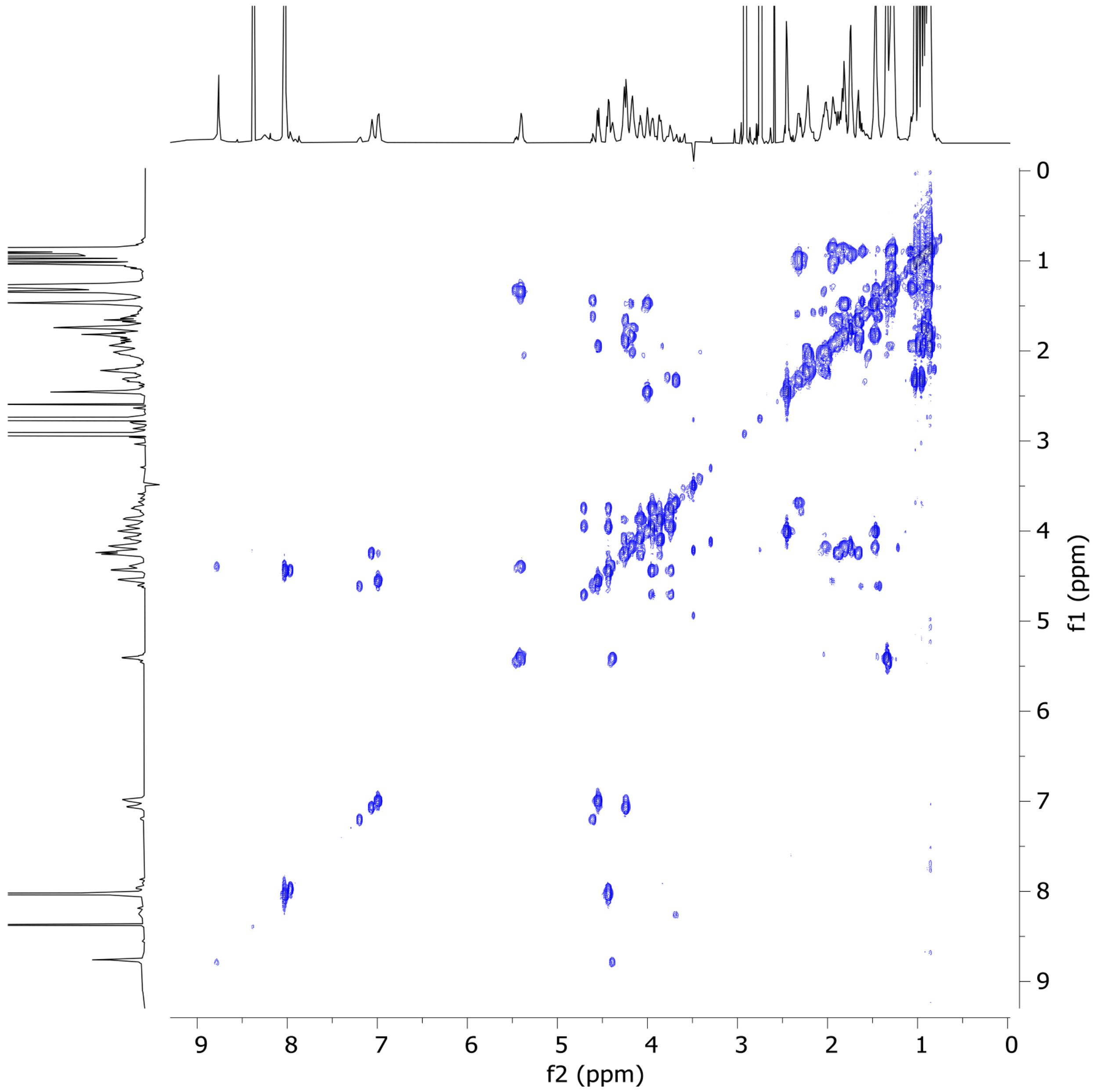


**Figure S22** 2D COSY NMR spectrum of viscosin I (600 MHz, DMF-d7, 298K).

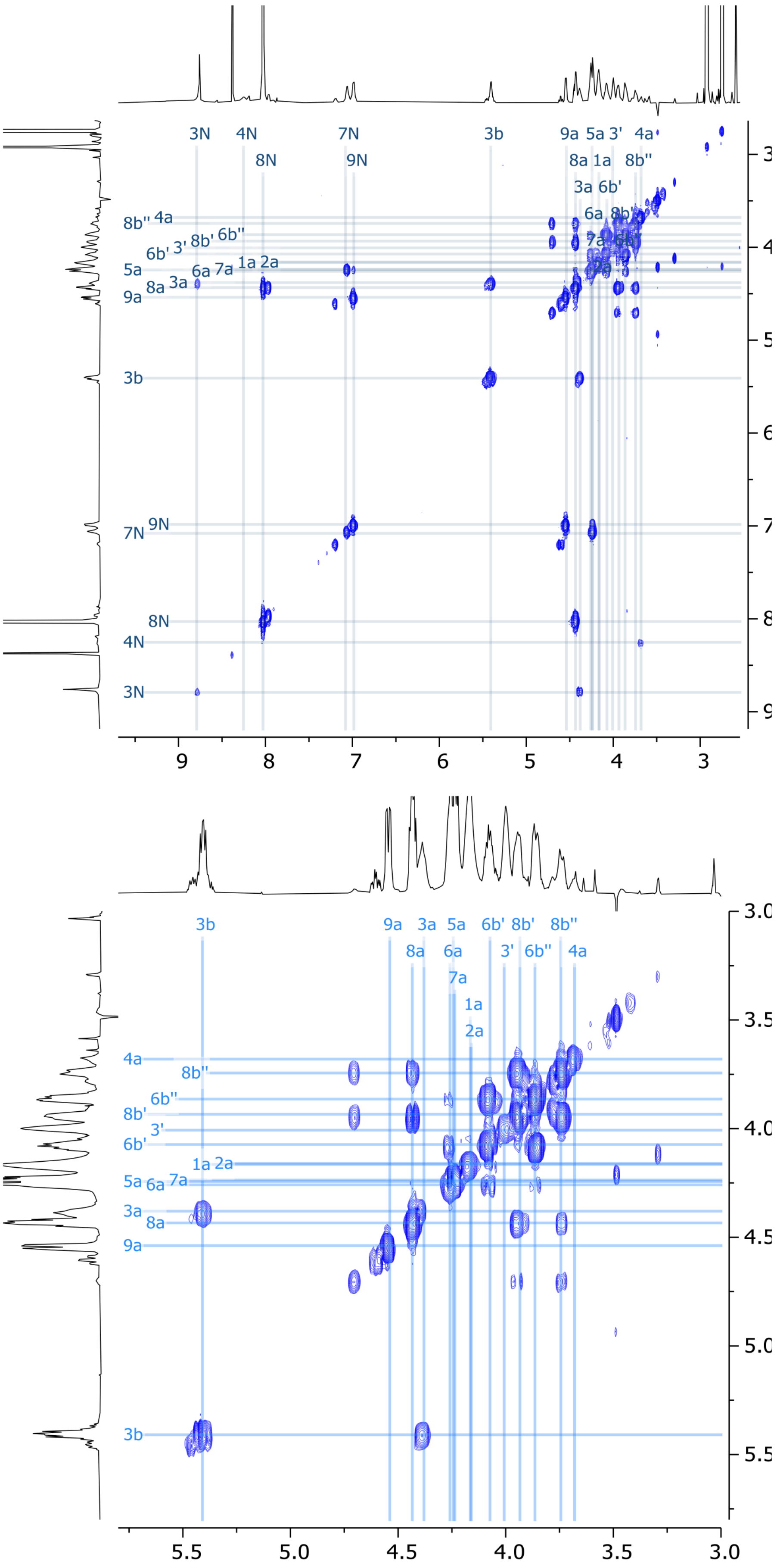


**Figure S23** Expanded regions of 2D COSY NMR spectrum of viscosin I (600 MHz, DMF-d7, 298K).

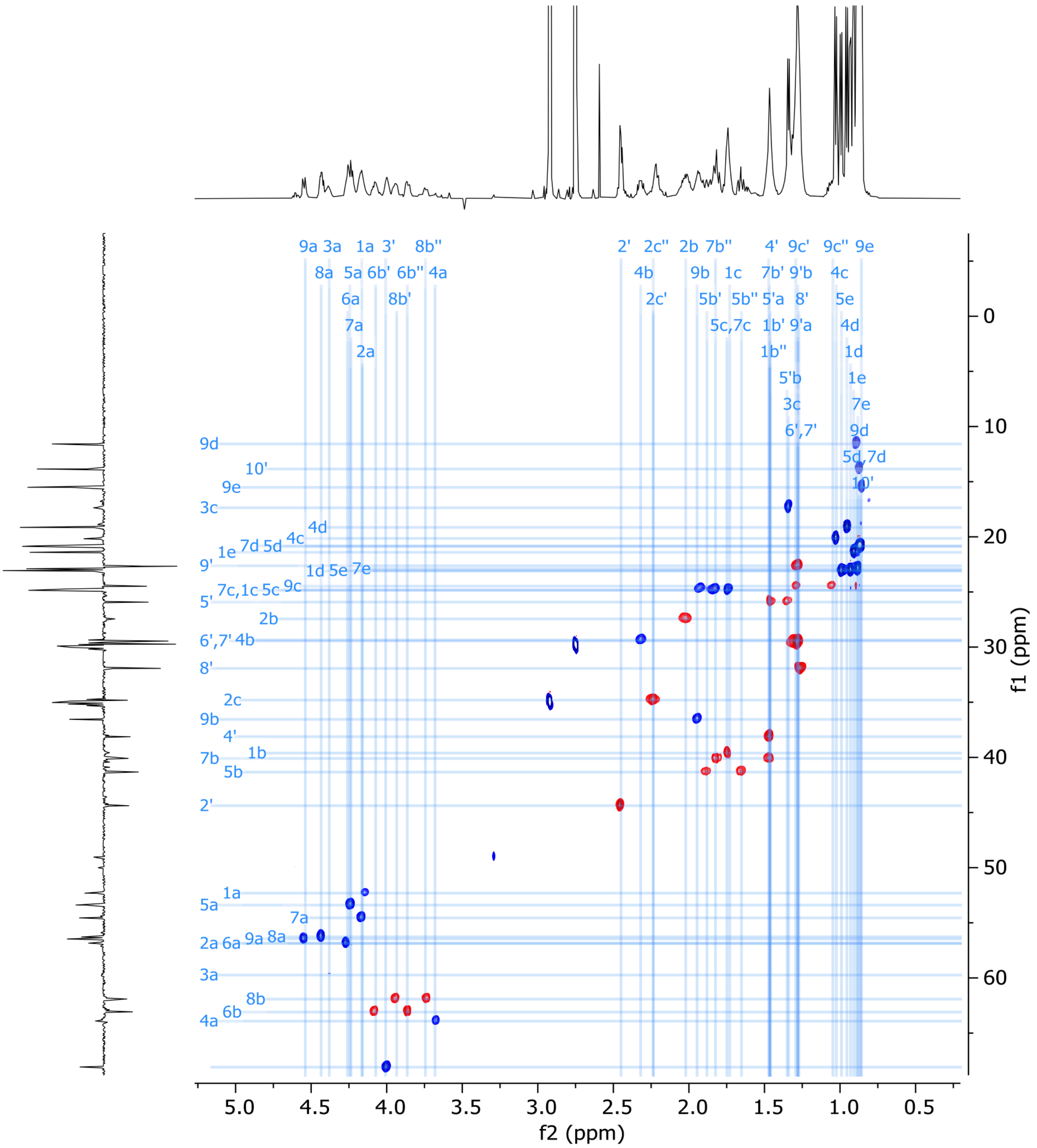


**Figure S24** 2D HSQC-EDITED NMR spectrum of viscosin I (600 MHz, DMF-d7, 298K).

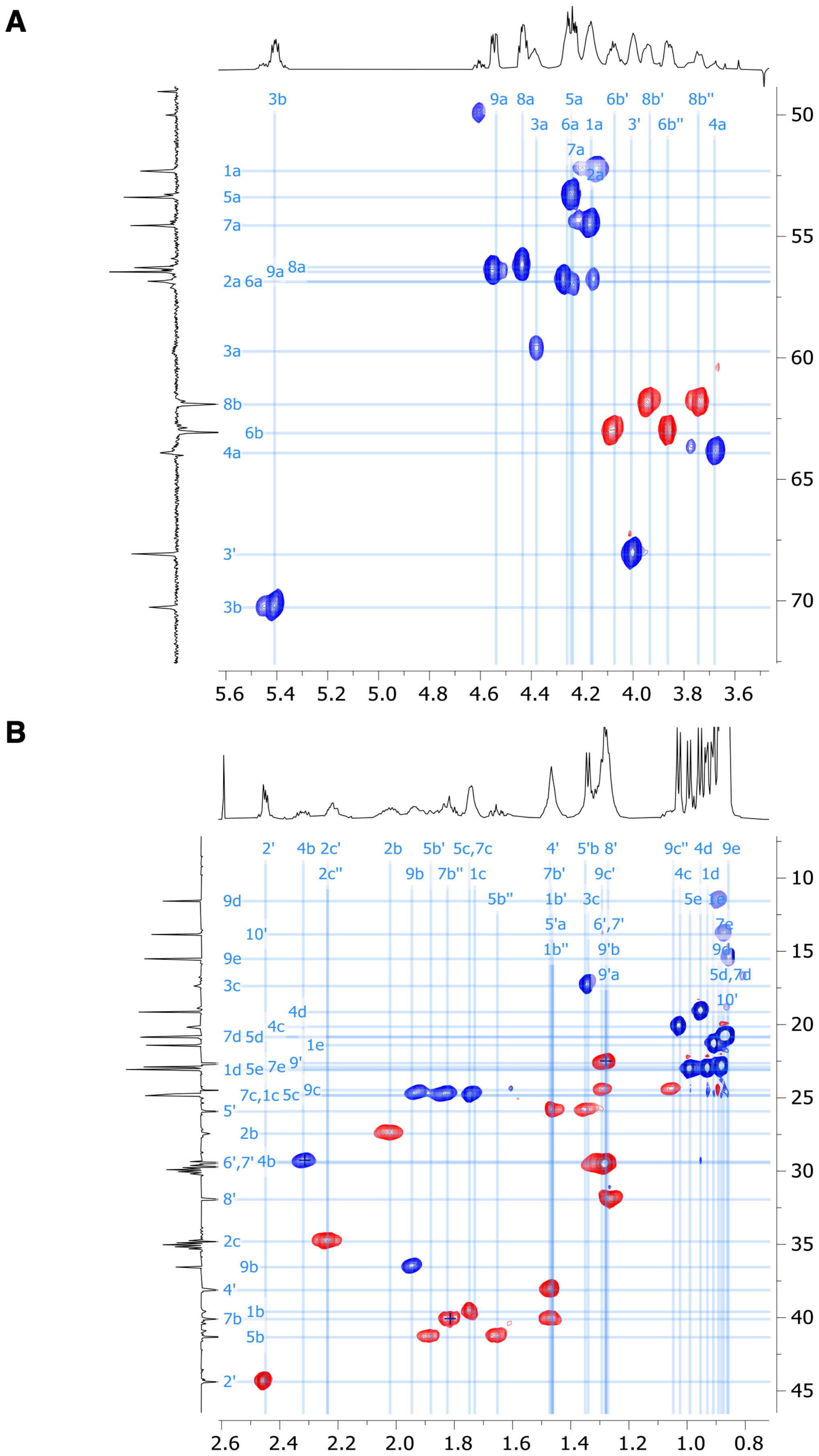


*Figure S25 continued overleaf. See next page for figure legend.*

**C**

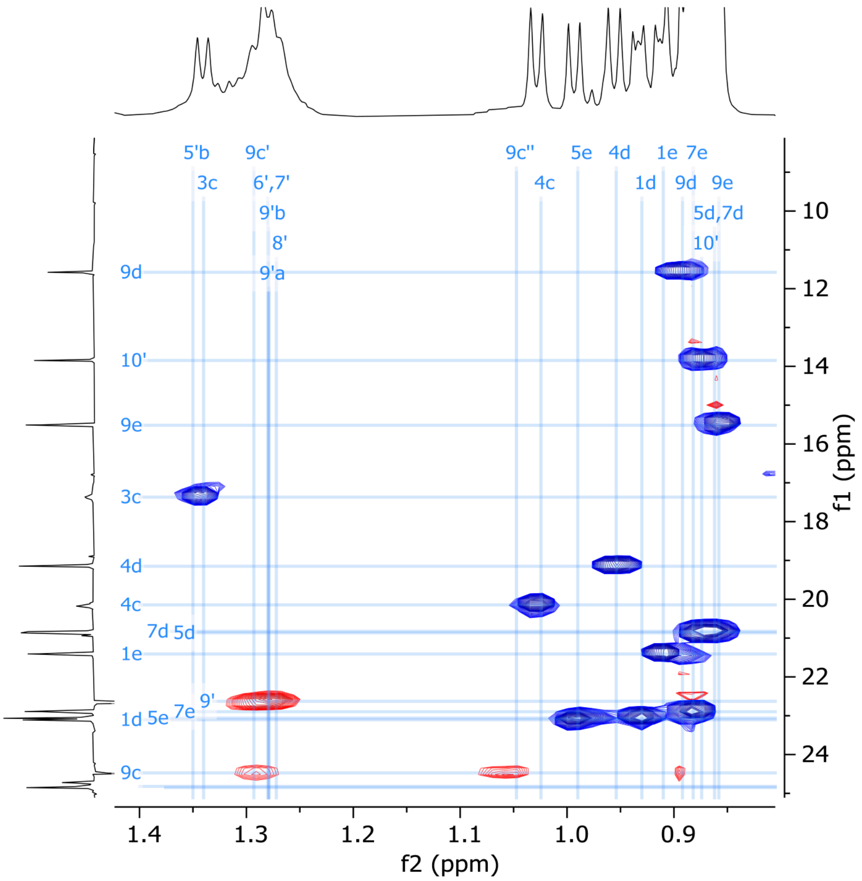


**Figure S25** Expansion of (A) α-C, (B) side chain and (C) Me-group regions of 2D HSQC-EDITED NMR spectrum of viscosin I (600 MHz, DMF-d7, 298K).

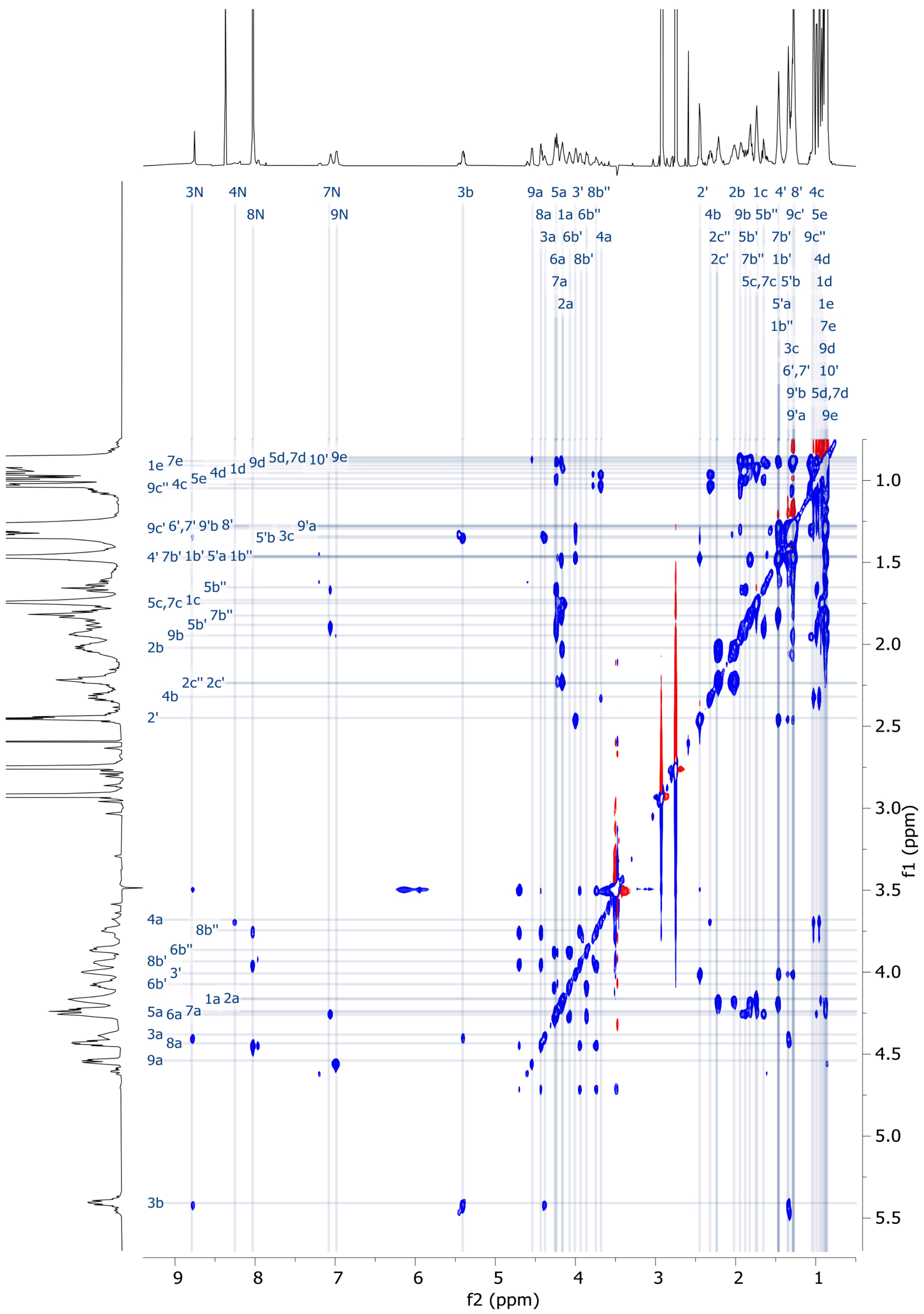


**Figure S26** 2D TOCSY NMR spectrum of viscosin I (600 MHz, DMF-d7, 298K).

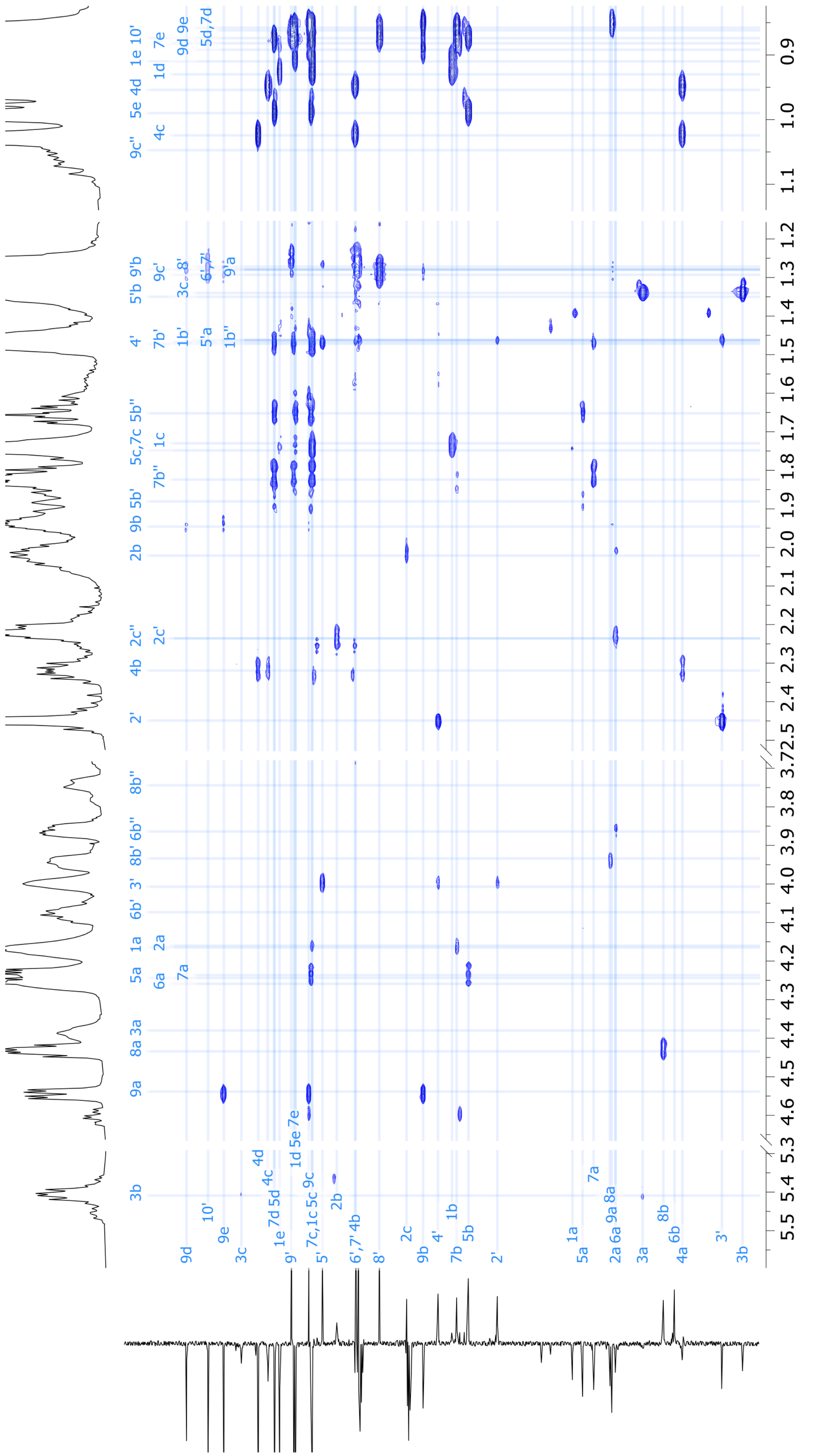


**Figure S27** 2D HMBC NMR spectrum of viscosin I (600 MHz, DMF-d7, 298K).

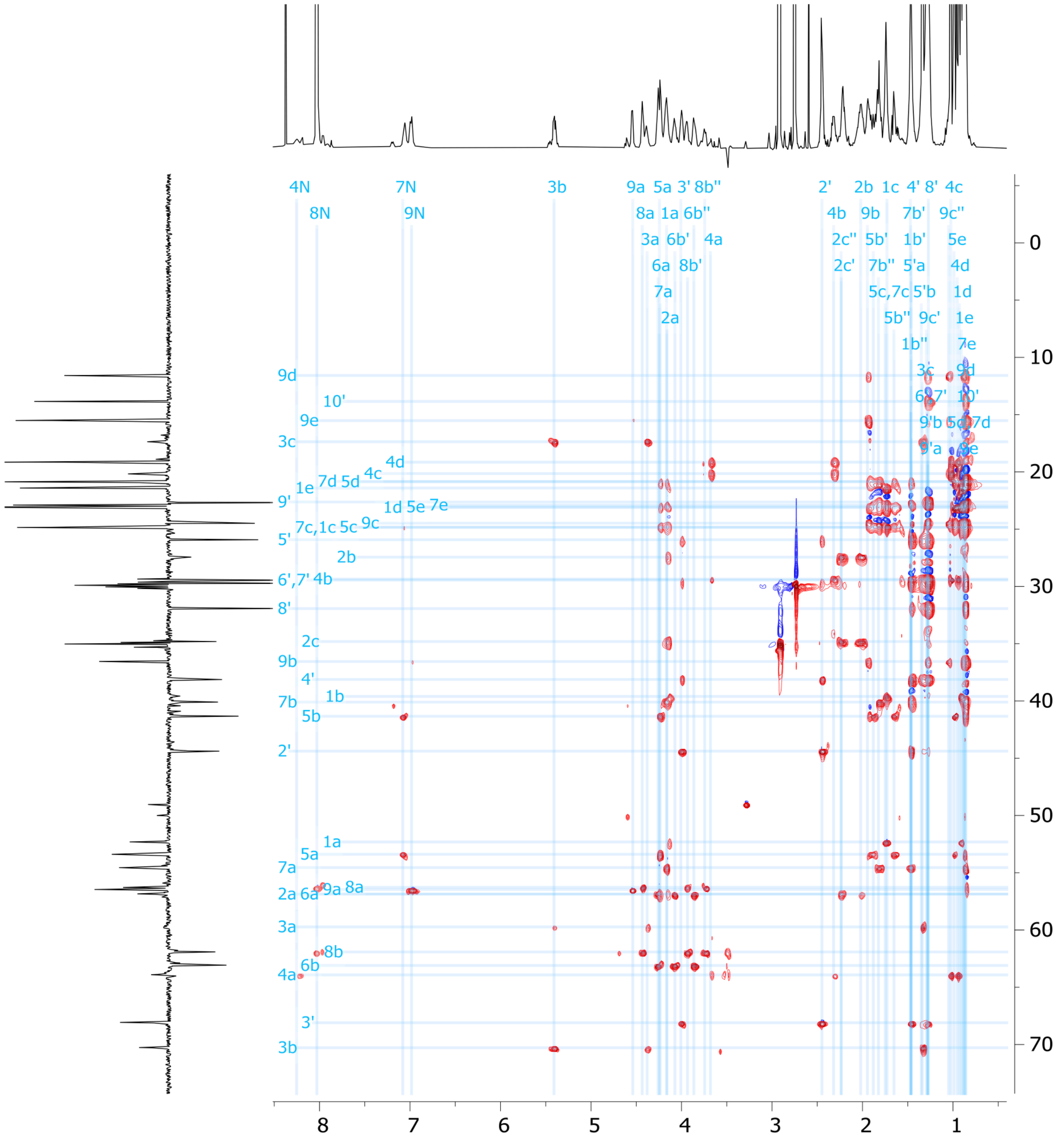


**Figure S28** 2D HSQC-TOCSY NMR spectrum of viscosin I (600 MHz, DMF-d7, 298K).

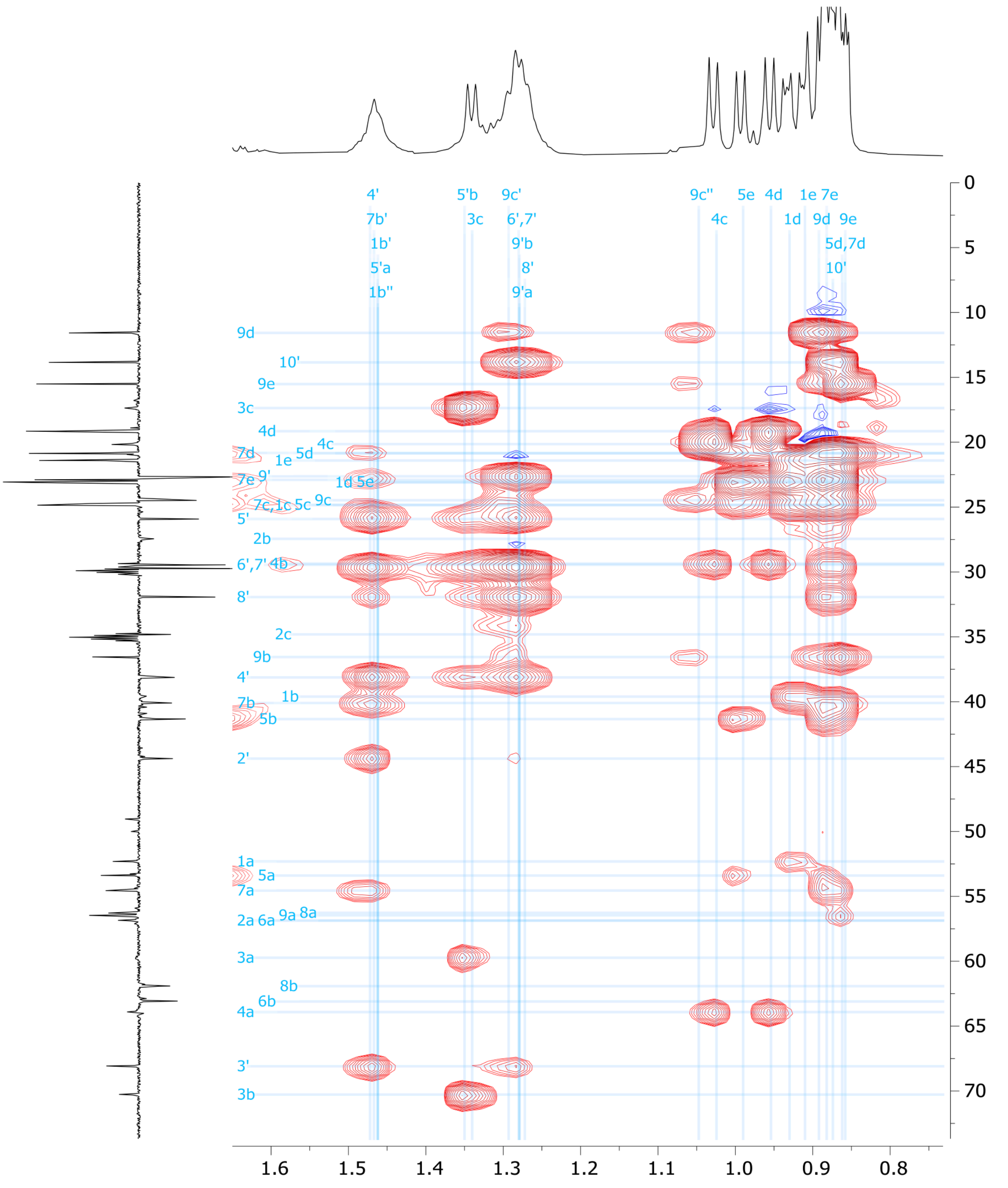

**Figure S29** High-field region of 2D HSQC-TOCSY NMR spectrum of viscosin I (600 MHz, DMF-d7, 298K).

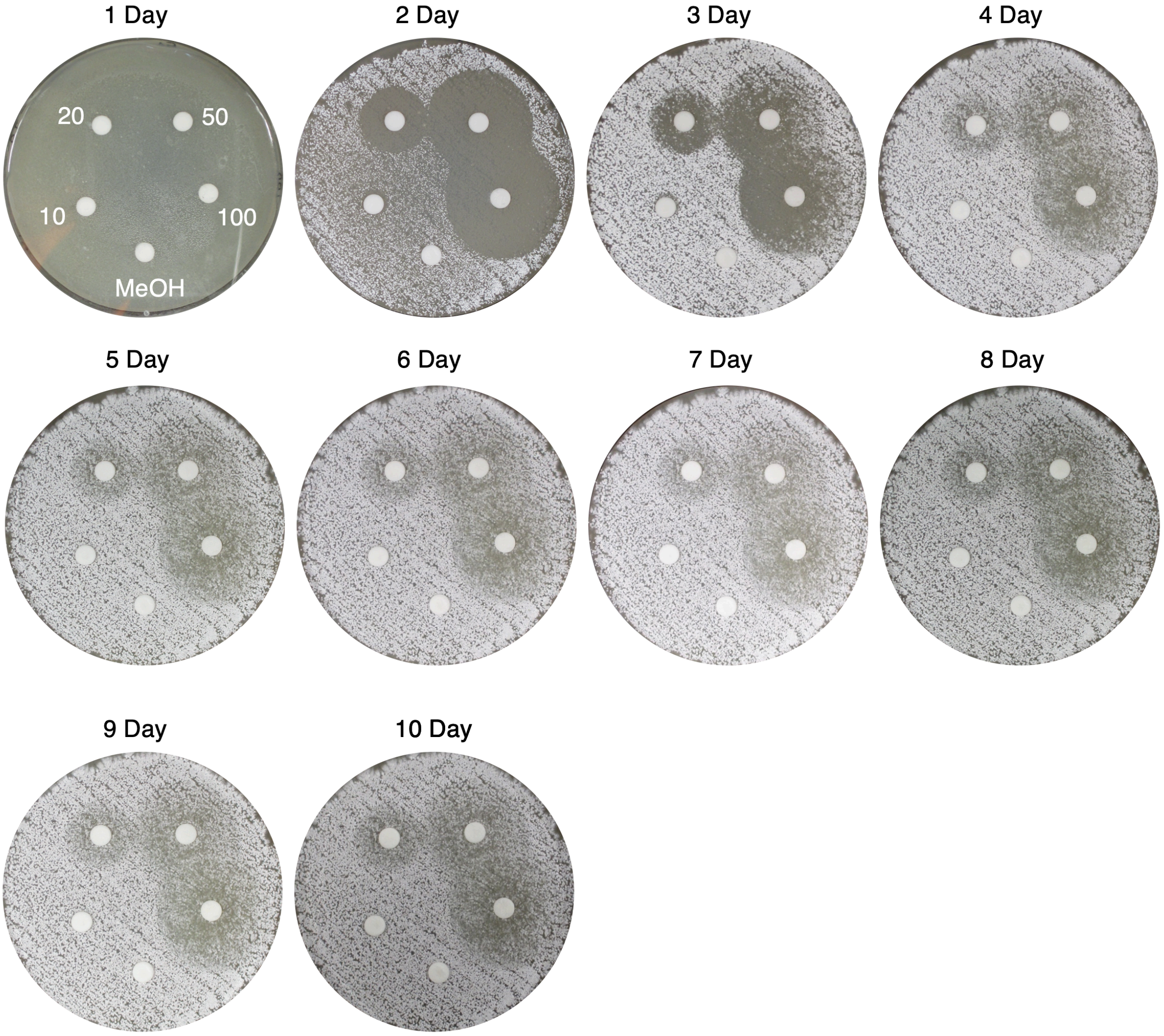


**Figure S30** Time course of viscosin I disk diffusion assays against *S. scabies* 87-22 on Instant Potato Medium (IPM). Viscosin I concentrations (µg/mL) are indicated, along with a methanol control.

| | Siderophore intensity | Protease activity | Motility | Congo red binding | Take-all inhibition | *S. scabies* inhibition | HCN production |
|---|---|---|---|---|---|---|---|
| **Siderophore intensity** | 1.00 | 0.44 | 0.25 | 0.33 | 0.24 | 0.30 | 0.33 |
| **Protease activity** | 0.44 | 1.00 | 0.24 | 0.21 | 0.18 | 0.45 | 0.14 |
| **Motility** | 0.25 | 0.24 | 1.00 | 0.28 | 0.35 | 0.52 | 0.39 |
| **Congo red binding** | 0.33 | 0.21 | 0.28 | 1.00 | 0.29 | 0.28 | 0.15 |
| **Take-all inhibition** | 0.24 | 0.18 | 0.35 | 0.29 | 1.00 | 0.31 | 0.31 |
| ***S. scabies* inhibition** | 0.30 | 0.45 | 0.52 | 0.28 | 0.31 | 1.00 | 0.52 |
| **HCN production** | 0.33 | 0.14 | 0.39 | 0.15 | 0.31 | 0.52 | 1.00 |

**Figure S31** Pearson correlation coefficients for phenotypes across both sequenced and unsequenced isolates. Isolates where one or more reliable phenotypes were not obtained (Supporting Dataset 1) were omitted from this correlation analysis.

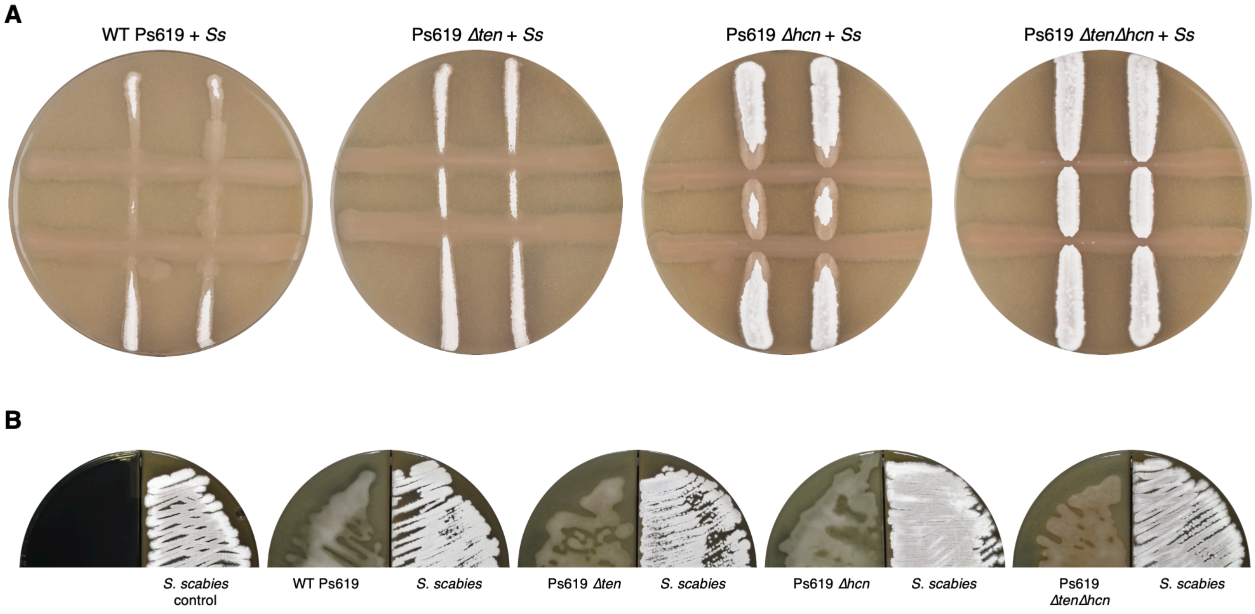


**Figure S32** Effect of Ps619 on *S. scabies* growth and development. (A) Cross streak assays (7 days post-inoculation) of Ps619 and associated mutants with *S. scabies* on drier conditions than in Fig. 6B. *S. scabies* (Ss) is streaked vertically first and then Ps619 is streaked horizontally. (B) Split plates (6 days post-inoculation) of *S. scabies* grown alongside Ps619 WT and mutant strains. Characteristic grey spore pigment of *S. scabies* (seen in control plate) is not present when grown alongside wild type or Δ*ten* strains.

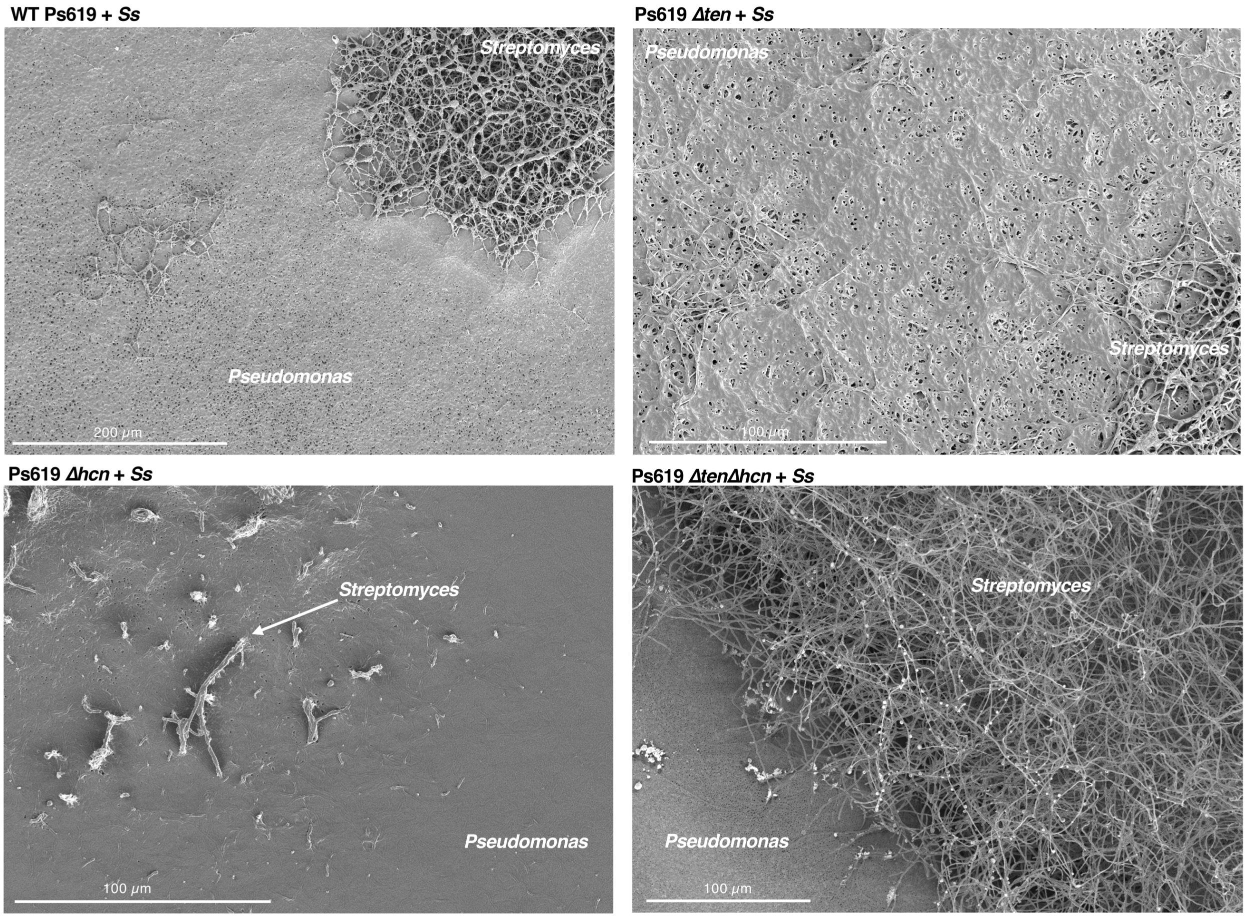


**Figure S33** Further cryo-SEM images of the interfacial region between the Ps619 strains and *S. scabies* (*Ss*) showing that a well-defined boundary only exists with the Ps619 Δ*ten*Δ*hcn* strain (bottom right).

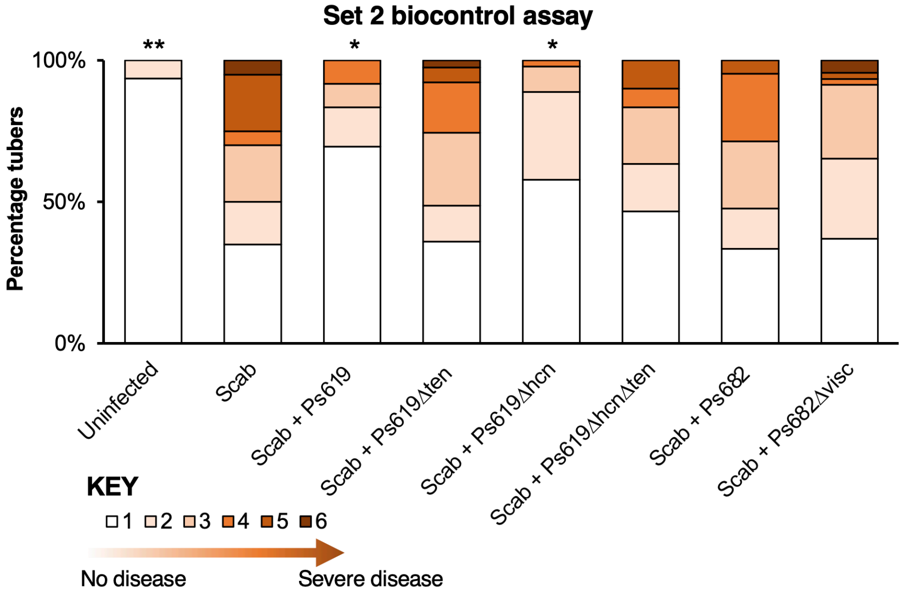


**Set 1 sampling summary**

| Condition | Number of plants | Number of tubers |
|---|---|---|
| Uninfected | 4 | 27 |
| Scab | 4 | 24 |
| Scab + Ps619 | 4 | 28 |
| Scab + Ps619Δ*ten* | 4 | 30 |
| Scab + Ps619Δ*hcn* | 4 | 27 |
| Scab + Ps619Δ*hcn*Δ*ten* | 4 | 33 |
| Scab + Ps682 | 4 | 31 |
| Scab + Ps682Δ*visc* | 4 | 30 |

**Set 1 statistical summary**

| Comparison | Mean score difference | *p* value | Statistical significance |
|---|---|---|---|
| Scab vs Uninfected | 1.013 | 0.0017 | ** |
| Scab vs Scab + Ps619 | 0.728 | 0.0291 | * |
| Scab vs Scab + Ps619Δ*ten* | 0.124 | 0.9936 | ns |
| Scab vs Scab + Ps619Δ*hcn* | 0.921 | 0.0043 | ** |
| Scab vs Scab + Ps619Δ*hcn*Δ*ten* | 0.435 | 0.3250 | ns |
| Scab vs Scab + Ps682 | 0.148 | 0.9831 | ns |
| Scab vs Scab + Ps682Δ*visc* | 0.088 | 0.9994 | ns |

**Set 2 sampling summary**

| Condition | Number of plants | Number of tubers |
|---|---|---|
| Uninfected | 3 | 30 |
| Scab | 3 | 20 |
| Scab + Ps619 | 4 | 35 |
| Scab + Ps619Δ*ten* | 4 | 39 |
| Scab + Ps619Δ*hcn* | 4 | 45 |
| Scab + Ps619Δ*hcn*Δ*ten* | 4 | 30 |
| Scab + Ps682 | 3 | 21 |
| Scab + Ps682Δ*visc* | 4 | 47 |

**Set 2 statistical summary**

| Comparison | Mean score difference | *p* value | Statistical significance |
|---|---|---|---|
| Scab vs Uninfected | 1.655 | 0.0014 | ** |
| Scab vs Scab + Ps619 | 1.165 | 0.0169 | * |
| Scab vs Scab + Ps619Δ*ten* | 0.041 | 0.9999 | ns |
| Scab vs Scab + Ps619Δ*hcn* | 1.168 | 0.0166 | * |
| Scab vs Scab + Ps619Δ*hcn*Δ*ten* | 0.647 | 0.3046 | ns |
| Scab vs Scab + Ps682 | 0.217 | 0.9859 | ns |
| Scab vs Scab + Ps682Δ*visc* | 0.592 | 0.3870 | ns |

**Figure S34** Results from set 2 of the potato scab biocontrol assay (set 1 data shown in main paper Figure 7) alongside statistical summaries of the two sets of experiments. This assay represents a full repeat of the set 1 experiment, which was started at a different time with a different set of plants. The bar chart shows the percentage of diseased tubers following infection with *S. scabies* ("Scab") along with treatment by Ps619, Ps682 and associated mutants. Tubers were scored using a disease severity index from 1 to 6 according to the method of Andrade *et al*. (86). Statistical analyses were calculated by using the average disease index of each plant (n = 3 or 4). *p* values were calculated using Dunnett's multiple comparison test, and asterisks indicate $p < 0.05$ (*), 0.01 (**) as compared to Scab treatment only (ns = not significant).

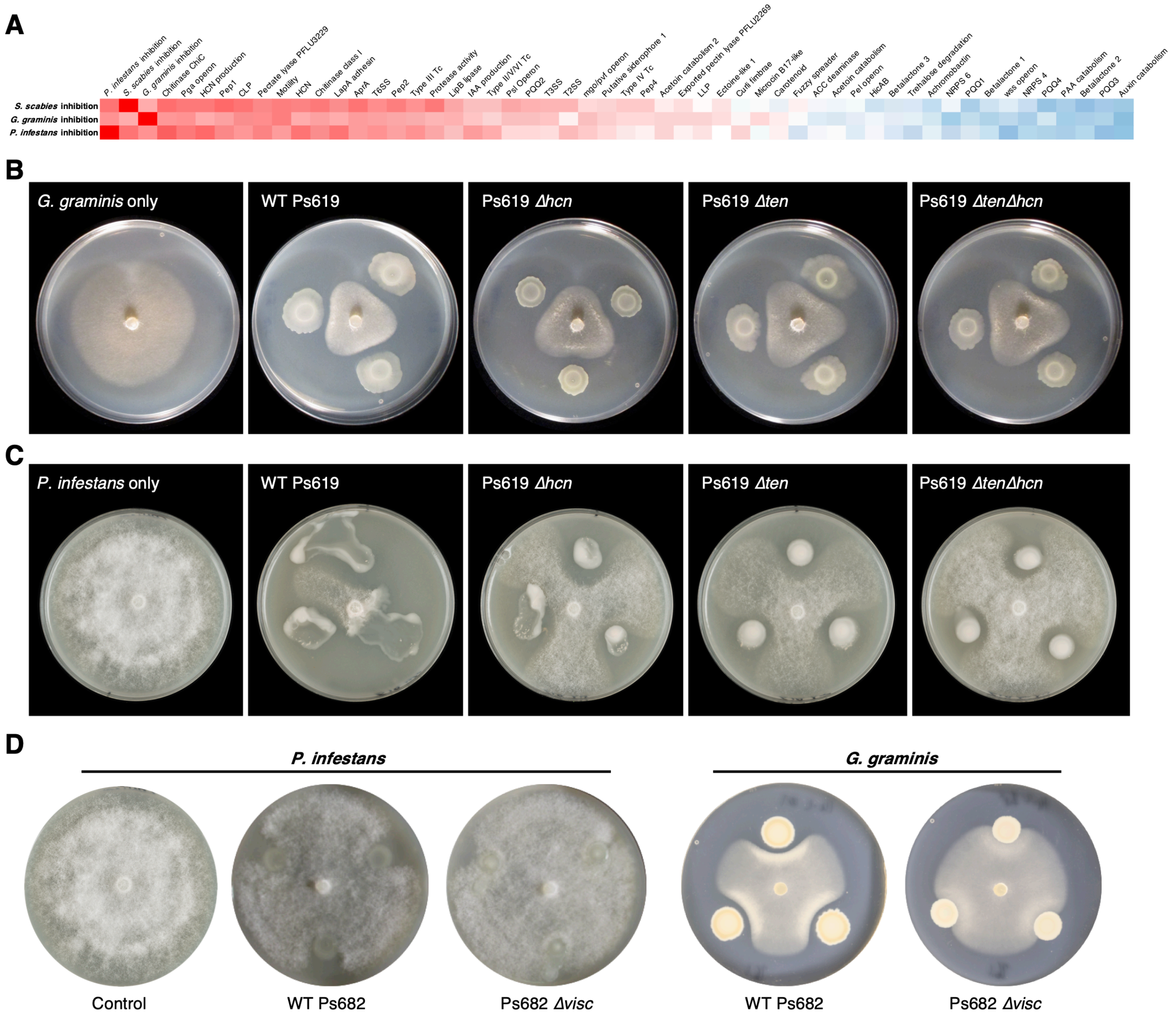


**Figure S35** Genotypes and phenotypes that correlate with the suppression of different plant pathogens. (A) Heatmap of Pearson correlation coefficients of pathogen inhibition versus genotypes and phenotypes (see Figure S8 for full correlations; the same colour scale is used). (B) On-plate inhibition of *G. graminis* growth by WT and mutant Ps619 strains. (C) On-plate inhibition of *P. infestans* growth by WT and mutant Ps619 strains. (D) WT and mutant Ps682 activity towards *P. infestans* and *G. graminis*.

### SUPPLEMENTARY REFERENCES


1. Blin, K., Shaw, S., Steinke, K., Villebro, R., Ziemert, N., Lee, S. Y., Medema, M. H. and Weber, T. (2019) antiSMASH 5.0: updates to the secondary metabolite genome mining pipeline. *Nucleic Acids Res.* **47**, W81–W87. DOI: 10.1093/nar/gkz310.
2. Kautsar, S. A., Blin, K., Shaw, S., Navarro-Muñoz, J. C., Terlouw, B. R., van der Hooft, J. J. J., van Santen, J. A., Tracanna, V., Suarez Duran, H. G., Pascal Andreu, V., Selem-Mojica, N., Alanjary, M., Robinson, S. L., Lund, G., Epstein, S. C., Sisto, A. C., Charkoudian, L. K., Collemare, J., Linington, R. G., Weber, T. and Medema, M. H. (2020) MIBiG 2.0: a repository for biosynthetic gene clusters of known function. *Nucleic Acids Res.* **48**, D454–D458. DOI: 10.1093/nar/gkz882.
3. Gross, H. and Loper, J. E. (2009) Genomics of secondary metabolite production by Pseudomonas spp. *Nat. Prod. Rep.* **26**, 1408–1446. DOI: 10.1039/b817075b.
4. Medema, M. H., Takano, E. and Breitling, R. (2013) Detecting sequence homology at the gene cluster level with MultiGeneBlast. *Mol. Biol. Evol.* **30**, 1218–1223. DOI: 10.1093/molbev/mst025.
5. Marchler-Bauer, A., Bo, Y., Han, L., He, J., Lanczycki, C. J., Lu, S., Chitsaz, F., Derbyshire, M. K., Geer, R. C., Gonzales, N. R., Gwadz, M., Hurwitz, D. I., Lu, F., Marchler, G. H., Song, J. S., Thanki, N., Wang, Z., Yamashita, R. A., Zhang, D., Zheng, C., Geer, L. Y. and Bryant, S. H. (2017) CDD/SPARCLE: functional classification of proteins via subfamily domain architectures. *Nucleic Acids Res.* **45**, D200–D203. DOI: 10.1093/nar/gkw1129.
6. Röttig, M., Medema, M. H., Blin, K., Weber, T., Rausch, C. and Kohlbacher, O. (2011) NRPSpredictor2–a web server for predicting NRPS adenylation domain specificity. *Nucleic Acids Res.* **39**, W362–W367. DOI: 10.1093/nar/gkr323.
7. Camacho, C., Coulouris, G., Avagyan, V., Ma, N., Papadopoulos, J., Bealer, K. and Madden, T. L. (2009) BLAST+: architecture and applications. *BMC Bioinformatics* **10**, 421. DOI: 10.1186/1471-2105-10-421.
8. Velasco, A., Acebo, P., Gomez, A., Schleissner, C., Rodríguez, P., Aparicio, T., Conde, S., Muñoz, R., la Calle, de, F., Garcia, J. L. and Sánchez-Puelles, J. M. (2005) Molecular characterization of the safracin biosynthetic pathway from *Pseudomonas fluorescens* A2-2: designing new cytotoxic compounds. *Mol. Microbiol.* **56**, 144–154. DOI: 10.1111/j.1365-2958.2004.04433.x.
9. Carrión, V. J., Gutiérrez-Barranquero, J. A., Arrebola, E., Bardaji, L., Codina, J. C., de Vicente, A., Cazorla, F. M. and Murillo, J. (2013) The mangotoxin biosynthetic operon (mbo) is specifically distributed within *Pseudomonas syringae* genomospecies 1 and was acquired only once during evolution. *Appl. Environ. Microbiol.* **79**, 756–767. DOI: 10.1128/AEM.03007-12.

10. Vallet Gely, I., Opota, O., Boniface, A., Novikov, A. and Lemaitre, B. (2010) A secondary metabolite acting as a signalling molecule controls *Pseudomonas entomophila* virulence. *Cell. Microbiol.* **12**, 1666–1679. DOI: 10.1111/j.1462-5822.2010.01501.x.
11. Kretsch, A. M., Morgan, G. L., Acken, K. A., Barr, S. A. and Li, B. (2021) Pseudomonas Virulence Factor Pathway Synthesizes Autoinducers That Regulate the Secretome of a Pathogen. *ACS Chem. Biol.* **16**, 501–509. DOI: 10.1021/acschembio.0c00901.
12. Cimermancic, P., Medema, M. H., Claesen, J., Kurita, K., Wieland Brown, L. C., Mavrommatis, K., Pati, A., Godfrey, P. A., Koehrsen, M., Clardy, J., Birren, B. W., Takano, E., Sali, A., Linington, R. G. and Fischbach, M. A. (2014) Insights into secondary metabolism from a global analysis of prokaryotic biosynthetic gene clusters. *Cell* **158**, 412–421. DOI: 10.1016/j.cell.2014.06.034.
13. Kato, J.-Y., Funa, N., Watanabe, H., Ohnishi, Y. and Horinouchi, S. (2007) Biosynthesis of gamma-butyrolactone autoregulators that switch on secondary metabolism and morphological development in *Streptomyces*. *Proc. Natl. Acad. Sci. U.S.A.* **104**, 2378–2383. DOI: 10.1073/pnas.0607472104.
14. Berti, A. D. and Thomas, M. G. (2009) Analysis of achromobactin biosynthesis by *Pseudomonas syringae* pv. *syringae* B728a. *J. Bacteriol.* **191**, 4594–4604. DOI: 10.1128/JB.00457-09.
15. Mercado-Blanco, J., van der Drift, K. M., Olsson, P. E., Thomas-Oates, J. E., van Loon, L. C. and Bakker, P. A. (2001) Analysis of the pmsCEAB gene cluster involved in biosynthesis of salicylic acid and the siderophore pseudomonine in the biocontrol strain *Pseudomonas fluorescens* WCS374. *J. Bacteriol.* **183**, 1909–1920. DOI: 10.1128/JB.183.6.1909-1920.2001.
16. Matthijs, S., Budzikiewicz, H., Schäfer, M., Wathelet, B. and Cornelis, P. (2008) Ornicorrugatin, a New Siderophore from *Pseudomonas fluorescens* AF76. *Zeitschrift für Naturforschung C* **63**, 8–12. DOI: 10.1515/znc-2008-1-202.
17. Cheng, X., de Bruijn, I., van der Voort, M., Loper, J. E. and Raaijmakers, J. M. (2013) The Gac regulon of *Pseudomonas fluorescens* SBW25. *Environ. Microbiol. Rep.* **5**, 608–619. DOI: 10.1111/1758-2229.12061.
18. Patel, H. M. and Walsh, C. T. (2001) In Vitro Reconstitution of the Pseudomonas aeruginosa Nonribosomal Peptide Synthesis of Pyochelin: Characterization of Backbone Tailoring Thiazoline Reductase and N-Methyltransferase Activities. *Biochemistry* **40**, 9023–9031. DOI: 10.1021/bi010519n.
19. Igarashi, Y., Asano, D., Sawamura, M., In, Y., Ishida, T. and Imoto, M. (2016) Ulbactins F and G, Polycyclic Thiazoline Derivatives with Tumor Cell Migration Inhibitory Activity from *Brevibacillus* sp. *Org. Lett.* **18**, 1658–1661. DOI: 10.1021/acs.orglett.6b00531.
20. Matthijs, S., Baysse, C., Koedam, N., Tehrani, K. A., Verheyden, L., Budzikiewicz, H., Schäfer, M., Hoorelbeke, B., Meyer, J.-M., De Greve, H. and Cornelis, P. (2004) The *Pseudomonas* siderophore quinolobactin is synthesized from xanthurenic acid, an

intermediate of the kynurenine pathway. *Mol. Microbiol.* **52**, 371–384. DOI: 10.1111/j.1365-2958.2004.03999.x.

21. Challis, G. L. (2005) A widely distributed bacterial pathway for siderophore biosynthesis independent of nonribosomal peptide synthetases. *ChemBioChem* **6**, 601–611. DOI: 10.1002/cbic.200400283.
22. San Millán, J. L., Kolter, R. and Moreno, F. (1985) Plasmid genes required for microcin B17 production. *J. Bacteriol.* **163**, 1016–1020.
23. Ghilarov, D., Stevenson, C. E. M., Travin, D. Y., Piskunova, J., Serebryakova, M., Maxwell, A., Lawson, D. M. and Severinov, K. (2019) Architecture of Microcin B17 Synthetase: An Octameric Protein Complex Converting a Ribosomally Synthesized Peptide into a DNA Gyrase Poison. *Mol. Cell* **73**, 749–762.e5. DOI: 10.1016/j.molcel.2018.11.032.
24. Metelev, M., Serebryakova, M., Ghilarov, D., Zhao, Y. and Severinov, K. (2013) Structure of microcin B-like compounds produced by *Pseudomonas syringae* and species specificity of their antibacterial action. *J. Bacteriol.* **195**, 4129–4137. DOI: 10.1128/JB.00665-13.
25. Santos-Aberturas, J., Chandra, G., Frattaruolo, L., Lacret, R., Pham, T. H., Vior, N. M., Eyles, T. H. and Truman, A. W. (2019) Uncovering the unexplored diversity of thioamidated ribosomal peptides in Actinobacteria using the RiPPER genome mining tool. *Nucleic Acids Res.* **47**, 4624–4637. DOI: 10.1093/nar/gkz192.
26. Kenney, G. E., Dassama, L. M. K., Pandelia, M.-E., Gizzi, A. S., Martinie, R. J., Gao, P., DeHart, C. J., Schachner, L. F., Skinner, O. S., Ro, S. Y., Zhu, X., Sadek, M., Thomas, P. M., Almo, S. C., Bollinger, J. M., Krebs, C., Kelleher, N. L. and Rosenzweig, A. C. (2018) The biosynthesis of methanobactin. *Science* **359**, 1411–1416. DOI: 10.1126/science.aap9437.
27. Sedkova, N., Tao, L., Rouvière, P. E. and Cheng, Q. (2005) Diversity of carotenoid synthesis gene clusters from environmental Enterobacteriaceae strains. *Appl. Environ. Microbiol.* **71**, 8141–8146. DOI: 10.1128/AEM.71.12.8141-8146.2005.
28. Pessi, G. and Haas, D. (2000) Transcriptional control of the hydrogen cyanide biosynthetic genes hcnABC by the anaerobic regulator ANR and the quorum-sensing regulators LasR and RhlR in *Pseudomonas aeruginosa*. *J. Bacteriol.* **182**, 6940–6949. DOI: 10.1128/JB.182.24.6940-6949.2000.
29. Carrión, O., Curson, A. R. J., Kumaresan, D., Fu, Y., Lang, A. S., Mercadé, E. and Todd, J. D. (2015) A novel pathway producing dimethylsulphide in bacteria is widespread in soil environments. *Nat. Commun.* **6**, 6579. DOI: 10.1038/ncomms7579.
30. Kinscherf, T. G. and Willis, D. K. (2005) The biosynthetic gene cluster for the beta-lactam antibiotic tabtoxin in *Pseudomonas syringae*. *J. Antibiot.* **58**, 817–821. DOI: 10.1038/ja.2005.109.
31. Palm, C. J., Gaffney, T. and Kosuge, T. (1989) Cotranscription of genes encoding indoleacetic acid production in *Pseudomonas syringae* subsp. *savastanoi*. *J. Bacteriol.* **171**, 1002–1009. DOI: 10.1128/jb.171.2.1002-1009.1989.

32. McClerklin, S. A., Lee, S. G., Harper, C. P., Nwumeh, R., Jez, J. M. and Kunkel, B. N. (2018) Indole-3-acetaldehyde dehydrogenase-dependent auxin synthesis contributes to virulence of *Pseudomonas syringae* strain DC3000. *PLoS Pathog.* **14**, e1006811. DOI: 10.1371/journal.ppat.1006811.
33. Laue, B. E., Jiang, Y., Chhabra, S. R., Jacob, S., Stewart, G. S., Hardman, A., Downie, J. A., O'Gara, F. and Williams, P. (2000) The biocontrol strain *Pseudomonas fluorescens* F113 produces the Rhizobium small bacteriocin, N-(3-hydroxy-7-cis-tetradecenoyl)homoserine lactone, via HdtS, a putative novel N-acylhomoserine lactone synthase. *Microbiology* **146**, 2469–2480. DOI: 10.1099/00221287-146-10-2469.
34. Kudo, F. and Eguchi, T. (2009) Biosynthetic genes for aminoglycoside antibiotics. *J. Antibiot.* **62**, 471–481. DOI: 10.1038/ja.2009.76.
35. Couch, R., O'Connor, S. E., Seidle, H., Walsh, C. T. and Parry, R. (2004) Characterization of CmaA, an Adenylation-Thiolation Didomain Enzyme Involved in the Biosynthesis of Coronatine. *J. Antibiot.* **186**, 35–42. DOI: 10.1128/JB.186.1.35-42.2004.
36. Bown, L., Li, Y., Berrué, F., Verhoeven, J. T. P., Dufour, S. C. and Bignell, D. R. D. (2017) Biosynthesis and evolution of coronafacoyl phytotoxin production in the common scab pathogen *Streptomyces scabies*. *Appl. Environ. Microbiol.* **83**, e01169–17. DOI: 10.1128/AEM.01169-17.
37. Kim, S. Y., Ju, K.-S., Metcalf, W. W., Evans, B. S., Kuzuyama, T. and van der Donk, W. A. (2012) Different biosynthetic pathways to fosfomycin in *Pseudomonas syringae* and *Streptomyces* species. *Antimicrob. Agents Chemother.* **56**, 4175–4183. DOI: 10.1128/AAC.06478-11.
38. Sagot, B., Gaysinski, M., Mehiri, M., Guigonis, J.-M., Le Rudulier, D. and Alloing, G. (2010) Osmotically induced synthesis of the dipeptide N-acetylglutaminylglutamine amide is mediated by a new pathway conserved among bacteria. *Proc. Natl. Acad. Sci. U.S.A.* **107**, 12652–12657. DOI: 10.1073/pnas.1003063107.
39. Robinson, S. L., Terlouw, B. R., Smith, M. D., Pidot, S. J., Stinear, T. P., Medema, M. H. and Wackett, L. P. (2020) Global analysis of adenylate-forming enzymes reveals β-lactone biosynthesis pathway in pathogenic Nocardia. *J. Biol. Chem.* **295**, 14826–14839. DOI: 10.1074/jbc.RA120.013528.
40. Seip, B., Galinski, E. A. and Kurz, M. (2011) Natural and engineered hydroxyectoine production based on the *Pseudomonas stutzeri* ectABCD-ask gene cluster. *Appl. Environ. Microbiol.* **77**, 1368–1374. DOI: 10.1128/AEM.02124-10.
41. Choi, O., Kim, J., Kim, J.-G., Jeong, Y., Moon, J. S., Park, C. S. and Hwang, I. (2008) Pyrroloquinoline quinone is a plant growth promotion factor produced by *Pseudomonas fluorescens* B16. *Plant Physiol.* **146**, 657–668. DOI: 10.1104/pp.107.112748.

42. Vior, N. M., Lacret, R., Chandra, G., Dorai-Raj, S., Trick, M. and Truman, A. W. (2018) Discovery and Biosynthesis of the Antibiotic Bicyclomycin in Distantly Related Bacterial Classes. *Appl. Environ. Microbiol.* **84**, e02828–17. DOI: 10.1128/AEM.02828-17.
43. Klapper, M., Götze, S., Barnett, R., Willing, K. and Stallforth, P. (2016) Bacterial Alkaloids Prevent Amoebal Predation. *Angew. Chem. Int. Ed.* **55**, 8944–8947. DOI: 10.1002/anie.201603312.
44. Uytterhoeven, B., Appermans, K., Song, L., Masschelein, J., Lathouwers, T., Michiels, C. W. and Lavigne, R. (2015) Systematic analysis of the kalimantacin assembly line NRPS module using an adapted targeted mutagenesis approach. *MicrobiologyOpen*. DOI: 10.1002/mbo3.326.
45. Schmidt, Y., van der Voort, M., Crüsemann, M., Piel, J., Josten, M., Sahl, H.-G., Miess, H., Raaijmakers, J. M. and Gross, H. (2014) Biosynthetic Origin of the Antibiotic Cyclocarbamate Brabantamide A (SB-253514) in Plant-Associated *Pseudomonas*. *ChemBioChem* **15**, 259–266. DOI: 10.1002/cbic.201300527.
46. Bauer, J. S., Ghequire, M. G. K., Nett, M., Josten, M., Sahl, H.-G., De Mot, R. and Gross, H. (2015) Biosynthetic Origin of the Antibiotic Pseudopyronines A and B in *Pseudomonas putida* BW11M1. *ChemBioChem* **16**, 2491–2497. DOI: 10.1002/cbic.201500413.
47. Delany, I., Sheehan, M. M., Fenton, A., Bardin, S., Aarons, S. and O'Gara, F. (2000) Regulation of production of the antifungal metabolite 2,4-diacetylphloroglucinol in *Pseudomonas fluorescens* F113: genetic analysis of phlF as a transcriptional repressor. *Microbiology* **146**, 537–543. DOI: 10.1099/00221287-146-2-537.
48. Mavrodi, D. V., Ksenzenko, V. N., Bonsall, R. F., Cook, R. J., Boronin, A. M. and Thomashow, L. S. (1998) A seven-gene locus for synthesis of phenazine-1-carboxylic acid by *Pseudomonas fluorescens* 2-79. *J. Bacteriol.* **180**, 2541–2548.
49. Hammer, P. E., Hill, D. S., Lam, S. T., van Pée, K. H. and Ligon, J. M. (1997) Four genes from *Pseudomonas fluorescens* that encode the biosynthesis of pyrrolnitrin. *Appl. Environ. Microbiol.* **63**, 2147–2154.
50. Calderón, C. E., Pérez-García, A., de Vicente, A. and Cazorla, F. M. (2013) The dar genes of *Pseudomonas chlororaphis* PCL1606 are crucial for biocontrol activity via production of the antifungal compound 2-hexyl, 5-propyl resorcinol. *Mol. Plant Microbe Interact.* **26**, 554–565. DOI: 10.1094/MPMI-01-13-0012-R.
51. Philmus, B., Shaffer, B. T., Kidarsa, T. A., Yan, Q., Raaijmakers, J. M., Begley, T. P. and Loper, J. E. (2015) Investigations into the Biosynthesis, Regulation, and Self-Resistance of Toxoflavin in *Pseudomonas protegens* Pf-5. *ChemBioChem* **16**, 1782–1790. DOI: 10.1002/cbic.201500247.
52. Lee, X., Fox, A., Sufrin, J., Henry, H., Majcherczyk, P., Haas, D. and Reimmann, C. (2010) Identification of the biosynthetic gene cluster for the *Pseudomonas aeruginosa* antimetabolite

L-2-amino-4-methoxy-trans-3-butenoic acid. *J. Bacteriol.* **192**, 4251–4255. DOI: 10.1128/JB.00492-10.

53. Jackson, K. D., Starkey, M., Kremer, S., Parsek, M. R. and Wozniak, D. J. (2004) Identification of psl, a Locus Encoding a Potential Exopolysaccharide That Is Essential for *Pseudomonas aeruginosa* PAO1 Biofilm Formation. *J. Bacteriol.* **186**, 4466–4475. DOI: 10.1128/JB.186.14.4466-4475.2004.
54. Spiers, A. J., Bohannon, J., Gehrig, S. M. and Rainey, P. B. (2003) Biofilm formation at the air–liquid interface by the *Pseudomonas fluorescens* SBW25 wrinkly spreader requires an acetylated form of cellulose. *Mol. Microbiol.* **50**, 15–27. DOI: 10.1046/j.1365-2958.2003.03670.x.
55. Friedman, L. and Kolter, R. (2004) Genes involved in matrix formation in Pseudomonas aeruginosa PA14 biofilms. *Mol. Microbiol.* **51**, 675–690.
56. Lind, P. A., Farr, A. D. and Rainey, P. B. (2017) Evolutionary convergence in experimental *Pseudomonas* populations. *ISME J.* **11**, 589–600. DOI: 10.1038/ismej.2016.157.
57. Maleki, S., Almaas, E., Zotchev, S., Valla, S. and Ertesvåg, H. (2016) Alginate Biosynthesis Factories in *Pseudomonas fluorescens*: Localization and Correlation with Alginate Production Level. *Appl. Environ. Microbiol.* **82**, 1227–1236. DOI: 10.1128/AEM.03114-15.
58. Freeman, B. C., Chen, C. and Beattie, G. A. (2010) Identification of the trehalose biosynthetic loci of *Pseudomonas syringae* and their contribution to fitness in the phyllosphere. *Environ. Microbiol.* **12**, 1486–1497. DOI: 10.1111/j.1462-2920.2010.02171.x.
59. Ferguson, G. C., Bertels, F. and Rainey, P. B. (2013) Adaptive divergence in experimental populations of *Pseudomonas fluorescens*. V. Insight into the niche specialist fuzzy spreader compels revision of the model *Pseudomonas* radiation. *Genetics* **195**, 1319–1335. DOI: 10.1534/genetics.113.154948.
60. Boyd, C. D., Smith, T. J., El-Kirat-Chatel, S., Newell, P. D., Dufrêne, Y. F. and O'Toole, G. A. (2014) Structural Features of the *Pseudomonas fluorescens* Biofilm Adhesin LapA Required for LapG-Dependent Cleavage, Biofilm Formation, and Cell Surface Localization. *J. Bacteriol.* **196**, 2775–2788. DOI: 10.1128/JB.01629-14.
61. de Bentzmann, S., Giraud, C., Bernard, C. S., Calderon, V., Ewald, F., Plésiat, P., Nguyen, C., Grunwald, D., Attree, I., Jeannot, K., Fauvarque, M.-O. and Bordi, C. (2012) Unique Biofilm Signature, Drug Susceptibility and Decreased Virulence in Drosophila through the *Pseudomonas aeruginosa* Two-Component System PprAB. *PLoS Pathog.* **8**, e1003052. DOI: 10.1371/journal.ppat.1003052.
62. Dueholm, M. S., Albertsen, M., Otzen, D. and Nielsen, P. H. (2012) Curli Functional Amyloid Systems Are Phylogenetically Widespread and Display Large Diversity in Operon and Protein Structure. *PLoS ONE* **7**, e51274. DOI: 10.1371/journal.pone.0051274.

63. Leveau, J. H. J. and Gerards, S. (2008) Discovery of a bacterial gene cluster for catabolism of the plant hormone indole 3-acetic acid. *FEMS Microbiol. Ecol.* **65**, 238–250. DOI: 10.1111/j.1574-6941.2008.00436.x.
64. Teufel, R., Mascaraque, V., Ismail, W., Voss, M., Perera, J., Eisenreich, W., Haehnel, W. and Fuchs, G. (2010) Bacterial phenylalanine and phenylacetate catabolic pathway revealed. *Proc. Natl. Acad. Sci. U.S.A.* **107**, 14390–14395. DOI: 10.1073/pnas.1005399107.
65. Saravanakumar, D. and Samiyappan, R. (2007) ACC deaminase from *Pseudomonas fluorescens* mediated saline resistance in groundnut (Arachis hypogea) plants. *J. Appl. Microbiol.* **102**, 1283–1292. DOI: 10.1111/j.1365-2672.2006.03179.x.
66. Huang, M., Oppermann, F. B. and Steinbüchel, A. (1994) Molecular characterization of the *Pseudomonas putida* 2,3-butanediol catabolic pathway. *FEMS Microbiol. Lett.* **124**, 141–150. DOI: 10.1111/j.1574-6968.1994.tb07276.x.
67. Scales, B. S., Erb-Downward, J. R., Huffnagle, I. M., LiPuma, J. J. and Huffnagle, G. B. (2015) Comparative genomics of *Pseudomonas fluorescens* subclade III strains from human lungs. *BMC Genomics* **16**, 1032. DOI: 10.1186/s12864-015-2261-2.
68. Rangel, L. I., Henkels, M. D., Shaffer, B. T., Walker, F. L., Davis, E. W., Stockwell, V. O., Bruck, D., Taylor, B. J. and Loper, J. E. (2016) Characterization of Toxin Complex Gene Clusters and Insect Toxicity of Bacteria Representing Four Subgroups of *Pseudomonas fluorescens*. *PLoS ONE* **11**, e0161120. DOI: 10.1371/journal.pone.0161120.
69. Li, G., Shen, M., Lu, S., Le, S., Tan, Y., Wang, J., Zhao, X., Shen, W., Guo, K., Yang, Y., Zhu, H., Rao, X., Hu, F. and Li, M. (2016) Identification and Characterization of the HicAB Toxin-Antitoxin System in the Opportunistic Pathogen *Pseudomonas aeruginosa*. *Toxins* **8**, 113. DOI: 10.3390/toxins8040113.
70. Powell, G. K. and Morris, R. O. (1986) Nucleotide sequence and expression of a *Pseudomonas savastanoi* cytokinin biosynthetic gene: homology with *Agrobacterium tumefaciens* tmr and tzs loci. *Nucleic Acids Res.* **14**, 2555–2565. DOI: 10.1093/nar/14.6.2555.
71. Borlee, B. R., Goldman, A. D., Murakami, K., Samudrala, R., Wozniak, D. J. and Parsek, M. R. (2010) *Pseudomonas aeruginosa* uses a cyclic-di-GMP-regulated adhesin to reinforce the biofilm extracellular matrix. *Mol. Microbiol.* **75**, 827–842. DOI: 10.1111/j.1365-2958.2009.06991.x.
72. Péchy-Tarr, M., Bruck, D. J., Maurhofer, M., Fischer, E., Vogne, C., Henkels, M. D., Donahue, K. M., Grunder, J., Loper, J. E. and Keel, C. (2008) Molecular analysis of a novel gene cluster encoding an insect toxin in plant-associated strains of *Pseudomonas fluorescens*. *Environ. Microbiol.* **10**, 2368–2386. DOI: 10.1111/j.1462-2920.2008.01662.x.
73. Schellenberger, U., Oral, J., Rosen, B. A., Wei, J.-Z., Zhu, G., Xie, W., McDonald, M. J., Cerf, D. C., Diehn, S. H., Crane, V. C., Sandahl, G. A., Zhao, J.-Z., Nowatzki, T. M., Sethi, A., Liu, L., Pan, Z., Wang, Y., Lu, A. L. and Wu, G. (2016) A selective insecticidal protein from

*Pseudomonas* for controlling corn rootworms. *Science*. **354**, 634–637 DOI: 10.1126/science.aaf6056.

74. Rainey, P. B. and Bailey, M. J. (1996) Physical and genetic map of the *Pseudomonas fluorescens* SBW25 chromosome. *Mol. Microbiol.* **19**, 521–533. DOI: 10.1046/j.1365-2958.1996.391926.x.
75. Mortishire-Smith, R. J., Nutkins, J. C., Packman, L. C., Rainey, P. B., Brodey, C. L., Johnstone, K. and Williams, D. H. (1991) Determination of the structure of an extracellular peptide produced by the mushroom saprotroph *Pseudomonas reactans*. *Tetrahedron* **47**, 3645–3654. DOI: 10.1016/S0040-4020(01)80877-2.
76. Scott, T. A., Heine, D., Qin, Z. and Wilkinson, B. (2017) An L-threonine transaldolase is required for L-threo-β-hydroxy-α-amino acid assembly during obafluorin biosynthesis. *Nat. Commun.* **8**, 15935. DOI: 10.1038/ncomms15935.
77. Choi, K.-H., Gaynor, J. B., White, K. G., Lopez, C., Bosio, C. M., Karkhoff-Schweizer, R. R. and Schweizer, H. P. (2005) A Tn7-based broad-range bacterial cloning and expression system. *Nat. Meth.* **2**, 443–448. DOI: 10.1038/nmeth765.
78. Garrido-Sanz, D., Meier-Kolthoff, J. P., Göker, M., Martín, M., Rivilla, R. and Redondo-Nieto, M. (2016) Genomic and Genetic Diversity within the *Pseudomonas fluorescens* Complex. *PLoS ONE* **11**, e0150183. DOI: 10.1371/journal.pone.0150183.
79. Edgar, R. C. (2004) MUSCLE: multiple sequence alignment with high accuracy and high throughput. *Nucleic Acids Res.* **32**, 1792–1797. DOI: 10.1093/nar/gkh340.
80. Stamatakis, A. (2014) RAxML version 8: a tool for phylogenetic analysis and post-analysis of large phylogenies. *Bioinformatics* **30**, 1312–1313. DOI: 10.1093/bioinformatics/btu033.
81. Letunic, I. and Bork, P. (2021) Interactive Tree Of Life (iTOL) v5: an online tool for phylogenetic tree display and annotation. *Nucleic Acids Res.* **49**, W293–W296. https://doi.org/10.1093/nar/gkab301.
82. Navarro-Muñoz, J. C., Selem-Mojica, N., Mullowney, M. W., Kautsar, S. A., Tryon, J. H., Parkinson, E. I., de Los Santos, E. L. C., Yeong, M., Cruz-Morales, P., Abubucker, S., Roeters, A., Lokhorst, W., Fernandez-Guerra, A., Cappelini, L. T. D., Goering, A. W., Thomson, R. J., Metcalf, W. W., Kelleher, N. L., Barona-Gómez, F. and Medema, M. H. (2020) A computational framework to explore large-scale biosynthetic diversity. *Nat. Chem. Biol.* **16**, 60–68. DOI: 10.1038/s41589-019-0400-9.
83. Olivares, P., Ulrich, E. C., Chekan, J. R., van der Donk, W. A. and Nair, S. K. (2016) Characterization of Two Late-Stage Enzymes Involved in Fosfomycin Biosynthesis in Pseudomonads. *ACS Chem. Biol.* **12**, 456–463. DOI: 10.1021/acschembio.6b00939.
84. Götze, S., Herbst-Irmer, R., Klapper, M., Görls, H., Schneider, K. R. A., Barnett, R., Burks, T., Neu, U. and Stallforth, P. (2017) Structure, Biosynthesis, and Biological Activity of the Cyclic Lipopeptide Anikasin. *ACS Chem. Biol.* **12**, 2498–2502. DOI: 10.1021/acschembio.7b00589.

85. Pauwelyn, E., Huang, C.-J., Ongena, M., Leclère, V., Jacques, P., Bleyaert, P., Budzikiewicz, H., Schäfer, M. and Höfte, M. (2013) New linear lipopeptides produced by *Pseudomonas cichorii* SF1-54 are involved in virulence, swarming motility, and biofilm formation. *Mol. Plant Microbe Interact.* **26**, 585–598. DOI: 10.1094/MPMI-11-12-0258-R.

86. Andrade, M. H. M. L., Niederheitmann, M., de Paula Ribeiro, S. R. R., Oliveira, L. C., Pozza, E. A. and Pinto, C. A. B. P. (2019) Development and validation of a standard area diagram to assess common scab in potato tubers. *Eur. J. Plant Pathol.* **154**, 739–750. DOI: 10.1007/s10658-019-01697-z.
